# The Splicing Factor XAB2 interacts with ERCC1-XPF and XPG for RNA-loop processing during mammalian development

**DOI:** 10.1101/2020.07.20.211441

**Authors:** Evi Goulielmaki, Maria Tsekrekou, Nikos Batsiotos, Mariana Ascensão-Ferreira, Eleftheria Ledaki, Kalliopi Stratigi, Georgia Chatzinikolaou, Pantelis Topalis, Theodore Kosteas, Janine Altmüller, Jeroen A. Demmers, Nuno L. Barbosa-Morais, George A. Garinis

**Affiliations:** Institute of Molecular Biology and Biotechnology, Foundation for Research and Technology-Hellas, GR70013, Heraklion, Crete, Greece; Department of Biology, University of Crete, Heraklion, Crete, Greece; Instituto de Medicina Molecular João Lobo Antunes, Faculdade de Medicina da Universidade de Lisboa, Avenida Professor Egas Moniz, 1649-028 Lisboa, Portugal; Cologne Center for Genomics (CCG), Institute for Genetics, University of Cologne, 50931, Cologne, Germany; Proteomics Center, Netherlands Proteomics Center, and Department of Biochemistry, Erasmus University Medical Center, the Netherlands

**Keywords:** XAB2, pre-mRNA splicing, DNA damage, Nucleotide Excision Repair, hepatic development

## Abstract

RNA splicing, transcription and the DNA damage response are intriguingly linked in mammals but the underlying mechanisms remain poorly understood. Using an *in vivo* biotinylation tagging approach in mice, we show that the splicing factor XAB2 interacts with the core spliceosome and that it binds to spliceosomal U4 and U6 snRNAs and pre-mRNAs in developing livers. XAB2 depletion leads to aberrant intron retention, R-loop formation and DNA damage in cells. Studies in illudin S-treated cells and *Csb^m/m^* developing livers reveal that transcription-blocking DNA lesions trigger the release of XAB2 from all RNA targets tested. Immunoprecipitation studies reveal that XAB2 interacts with ERCC1-XPF and XPG endonucleases outside nucleotide excision repair and that the trimeric protein complex binds RNA:DNA hybrids under conditions that favor the formation of R-loops. Thus, XAB2 functionally links the spliceosomal response to DNA damage with R-loop processing with important ramifications for mammalian development and transcription-coupled DNA repair disorders.

## Introduction

The spliceosome is a highly dynamic, ribonucleoprotein complex that comprises five small nuclear (sn) RNAs (the U1, U2, U4, U5, and U6 snRNAs) and a growing number of associated splicing factors that enable the selective intron excision of nascent pre-mRNA transcripts prior to mRNA translation ^1^. Splicing initiates with the binding of the U1 snRNP to the GU sequence at the 5’ splice site of an intron and the zinc finger protein splicing factor 1 (SF1) that binds to the intron branch point sequence. The U2 snRNP displaces SF1 and binds to the branch point sequence followed by ATP hydrolysis. The snRNPs U2 and U4/U6 position the 5′ end and the branch point in proximity. The latter stimulates snRNP U5 and the 3′ end of the intron is brought into proximity and joined to the 5′ end. The resulting lariat is released with U2, U5, and U6 bound to it ^2^. Pre-mRNA splicing is functionally coupled to transcription ^3,4^. The process of mRNA synthesis is known to regulate constitutive and alternative splicing (AS) through the physical association of splicing factors to the elongating RNAPII ^5^, the 3D chromatin structure ^6,7^ or through the fine-tuning of RNA polymerase II (RNAPII) elongation rates ^8^. Recent findings support the notion that DNA damage is inherently linked to splicing ^9-11^. The presence of irreparable DNA lesions may influence RNA splicing through the selective nuclear transport of splicing factors ^12,13^, the inhibition of the elongating RNAPII ^9^, the interaction of RNAPII- and spliceosome-associated factors ^14^ or the displacement of the spliceosome when RNAPII is stalled at DNA damage sites ^11^.

XPA-binding protein (XAB)-2 is the human homologue of the yeast pre-mRNA splicing factor Syf1 that contains 15 tetratricopeptide repeats shared by proteins involved in RNA processing ^15,16^. Disruption of the *Xab2* gene results in pre-implantation lethality in mice ^17^. Thus, there are currently no available data to address the functional role of XAB2 in transcript maturation and the DNA damage response (DDR) *in vivo*. Using a yeast two-hybrid screen and *in vitro* pull-down assays, XAB2 was shown to interact with the nucleotide excision repair (NER) factors XPA, CSA and CSB proteins and with RNAPII ^18^. The yeast homolog of XAB2, Syf1 was also shown to be a TREX-interacting factor ^19^. Microinjection of antibodies raised against XAB2 inhibited the recovery of RNA synthesis after UV irradiation and normal RNA synthesis in fibroblasts ^18^. Follow-up work showed that XAB2 interacts with PRP19 ^20^ an ubiquitin-protein ligase involved in pre-mRNA splicing and DNA repair ^21,22^ and Aquarius (AQR) an intron-binding spliceosomal factor that links pre-mRNA splicing to small nucleolar ribonucleoprotein biogenesis ^23^. Moreover, XAB2 is functionally involved in homology-directed repair and single-strand annealing ^24^ and it was recently shown to be directly involved in pre-mRNA splicing of many genes *in vitro*, including POLR2A ^25^.

To dissect the functional contribution of XAB2 during mammalian development, we established an *in vivo* biotinylation tagging approach in mice. Our findings provide evidence that XAB2 is essential for NER, the pre-mRNA splicing and for R-loop processing during mammalian development. We show that XAB2 is part of a core spliceosome complex that binds on UsnRNAs. We find that the accumulation of transcription-blocking DNA lesions in cells treated with illudin S or in *Csb^m/m^* developing livers carrying an inborn defect in the transcription-coupled subpathway of NER (TC-NER) trigger the release of XAB2 from all RNA targets tested. RNAi-mediated knockdown of XAB2 leads to decreased RNA synthesis, aberrant intron retention, R-loop accumulation and DNA damage. Using a series of immunoprecipitation approaches, we find that XAB2 interacts with ERCC1-XPF and XPG endonucleases, independently of NER and that the XAB2 complex is recruited to RNA:DNA hybrids under conditions that favor the accumulation of R-loops. We propose that XAB2 functionally links persistent DNA damage with the core spliceosome and the processing ofR-loops during mammalian development highlighting the causal contribution of transcription-blocking DNA lesions to the progeroid and developmental defects associated with TC-NER disorders.

## Results

### Generation of biotin-tagged XAB2 mice

Ablation of *Xab2* in the murine germline results in preimplantation lethality, indicating an essential role for XAB2 during mouse embryogenesis ^17^. To delineate the functional role of XAB2 during mammalian development, we generated knock-in animals expressing XAB2 fused with a tandem affinity purification tag comprising two affinity moieties, a 1X FLAG tag and a 15aa Avitag sequence, separated by a Tobacco Etch Virus (TEV) site for easy tag removal. The tag was inserted before the stop codon of the last exon 19 (**Figure 1A**). The targeting vector was transfected to 129/SV ES cells expressing the Protamine 1-Cre recombinase transgene, which efficiently excises the neomycin cassette in the male germ line ^26^. After selecting properly targeted clones (**Figure 1B**; as indicated), two independently transfected clones were used to generate germ line-transmitting chimeras. Avitag-fused heterozygous males (*aviXab2*^+/-^ mice) were backcrossed and maintained in a C57/BL6 background. Homozygous *aviXab2*^+/+^ knock-in animals were then crossed with mice ubiquitously expressing the HA-tagged BirA biotin ligase transgene ^27^. BirA is a bacterial ligase that specifically recognizes and successfully biotinylates the short 15aa biotinylation Avitag, thus creating a high affinity “handle” for the *in vivo* isolation of XAB2-bound protein complexes in aviXab2^+/+^;birA (from now on designated as bXAB2) mice by binding to streptavidin. Importantly, biotinylation of the inserted Avitag sequence does not interfere with murine development as bXAB2 animals are born at the expected Mendelian frequency (**Figure 1C**), grow normally (**Figure 1D**) and show no developmental defects or other pathological features (**Figure 1E**). Nuclear extracts of P15 livers from aviXAB2 or bXAB2 mice were probed with streptavidin-HRP (stp-HRP) confirming the biotinylation of bXAB2 *in vivo* (**Figure 1F**). The use of anti-FLAG and anti-HA confirmed the presence of the knock-in allele and the BirA transgene, respectively. Pulldowns with 0.6 mg of nuclear extracts from livers of P15 bXAB2^+/+^ mice revealed the percentage of biotinylated XAB2 in the pulldown (90%) and flow through (fth;10%) fraction (**Figure 1G**). Previous findings suggest that XAB2 is involved in the transcription-coupled repair (TC-NER) sub-pathway of nucleotide excision repair (NER) ^18^ and during the end resection step of homologous recombination repair ^24^. To test whether the Avitag sequence interferes with the putative function of XAB2 in DNA repair *in vivo*, we further exposed mouse embryonic fibroblasts (MEFs) to ultraviolet irradiation (UV) as well as to mitomycin C (MMC). MMC is a potent genotoxic agent which inhibits replication and transcription of DNA by introducing DNA inter-strand crosslinks that prevent dissociation of the strands. We find a similar percentage of UV-induced unscheduled DNA synthesis in bXAB2 cells compared to wild-type (wt.) controls (**Figure 1H**) and that bXAB2 MEFs are not hypersensitive to UV irradiation (**Figure 1I**) or to MMC (**Figure 1J**). Thus, bXAB2 animals develop normally to adulthood and are proficient at repairing UV-induced cyclobutane pyrimidine dimers (CPDs) and DNA interstrand cross-links (ICLs).

**Figure 1.**
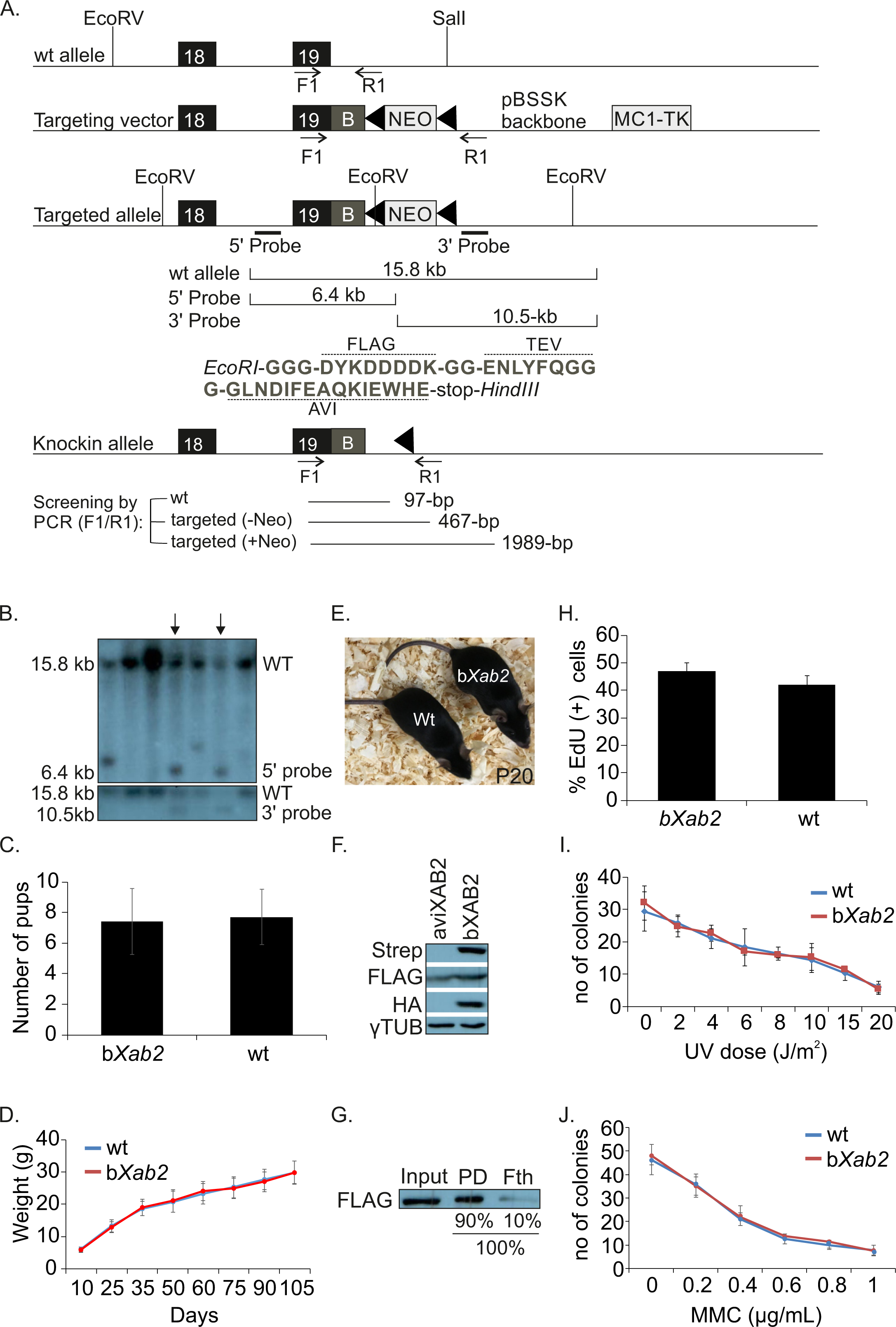
Generation of biotin-tagged XAB2 animals. (**A**). Schematic representation of knock-in mice expressing XAB2 fused with the 1xFLAG tag, a 15aa Avi tag sequence and a Tobacco Etch Virus (TEV) site separating the two tags. EcoRI and HindIII sites are synthetic; B: Flag-Tev-AVI fragment. (**B**). Two independently transfected ES clones (marked with arrow heads) were used to generate germline transmitting chimeras that were backcrossed with C57Bl/6 mice to generate *aviXab2*^+/-^ pups. Homozygous *aviXab2*^+/+^ knock-in animals (aviXAB2) were then crossed with mice ubiquitously expressing the HA-tagged BirA biotin ligase transgene (*aviXab2*^+/+^;birA, designated as bXAB2). (**C**). Number of wt. and bXAB2 pups born. (**D**). Weight (grams; g) of wt. and bXAB2 animals (n=8) at the indicated time points. (**E**). A photograph of P20 wt. and bXAB2 animals. (**F**). *In vivo* biotinylation of the short 15aa Avi-tag in bXAB2 animals. Nuclear extracts from P15 livers of mice expressing either only aviXAB2 (XAB2, MW: 110kDa) or aviXAB2 and BirA biotin ligase (bXAB2) were tested by Western blot. The blot was probed with streptavidin-HRP (stp-HRP), which confirms biotinylation of XAB2 *in vivo*, with anti-FLAG and anti-HA confirming the presence of the knock-in allele and of the BirA transgene, respectively. (**G**). Biotinylation efficiency in bXAB2 livers. The percentage of biotinylated XAB2 in the pulldown and (90%) and flow through fraction (10%; fth) was calculated by performing pull-down with 0.7 mg of nuclear extract derived from 15-day old bXAB2^+/+^;BirA livers and M-280 paramagnetic beads in excess. (**H**). % of EdU (+) MEFs derived from bXAB2 and wt. mice 2.5hrs after UV irradiation. (**I**). Survival of primary bXAB2 and wt. MEFs to UV (n ≥ 3 per time point) or (**J**). MMC at the indicated doses (n ≥ 3 per dose point). PD: Pulldown; Fth: Flow through. The images shown on Fig. 1F and 1G are representative of experiments that were repeated more than three times.

### A proteomics strategy reveals XAB2-bound protein partners involved in RNA processing and DNA repair

To isolate and characterize XAB2-associated protein complexes during postnatal hepatic development, we combined the *in vivo* biotinylation tagging approach ^27^ with a hypothesis-free, high-throughput proteomics strategy. To do this, we prepared nuclear extracts from livers of P15 bXAB2 and BirA mice using high-salt extraction conditions (**Figure 2A**). The liver is a relatively homogeneous organ that accurately depicts the growth status of the developing animal. Nuclear extracts were treated with benzonase and RNase A, to ensure that DNA or RNA does not mediate the identified protein interactions. Nuclear extracts were further incubated with streptavidin-coated beads and bound proteins were eluted and subjected to Western blot analysis confirming that bXAB2 can still interact with the pre-mRNA-processing factor 19 (PRP19) ^28^ (**Figure 2B**). Next, the proteome was separated by 1D SDS-PAGE (∼12 fractions) followed by in-gel digestion and peptides were analyzed with high-resolution liquid chromatography-tandem mass spectrometry (nLC MS/MS) on a hybrid linear ion trap Orbitrap instrument (**Figure 2C**). From three biological replicates, which comprised a total of 72 MS runs, we identified a total of 1167 proteins (**Table S1**) with 636 proteins (54,49%) shared between all three measurements under stringent selection criteria (**Figure 2D, Table S2**; see STAR Methods). To functionally characterize this dataset, we next subjected the 636-shared XAB2-bound proteins to gene ontology (GO) classification. Those biological pathways (**Figure 2E**) or processes (**Figure S1A**) containing a significantly disproportionate number of proteins relative to the murine proteome were flagged as significantly over-represented (FDR<0.05). At this confidence level, the over-represented biological processes and pathways involved 372 out of the initial 636 XAB2-bound core proteins; the latter set of proteins also showed a significantly higher number of known protein interactions (i.e. 958 interactions) than expected by chance (i.e. 531 interactions; **Figure 2F**) indicating a functionally relevant and highly interconnected protein network. Using this dataset, we were able to discern four major XAB2-associated protein complexes involved in i. pre-mRNA splicing (p≤3.2x10^-38^, e.g. AQR, BCAS2, PRP19, PRP8, SNRP40), ii. RNA transport (p≤1.8x10^-7^, e.g. NUP107, NUP133, NUP153, NUP160, NUP205, NUP210, NUP50, NUPl2), iii. Cell cycle (p≤7.1x10^-6^, e.g. ANAPC1, ANAPC 2, ANAPC 4, ANAPC 5, ANAPC 7, CDC23, CDC27, FZR1, RAD21, SMC1a, SMC3, STAG1, STAG2) and iv. Ribosome biogenesis (p≤4.8x10^-6^, e.g. BMS1, GNL3, MDN1, NOP58, UTP15, WDR36, WDR43, WDR75, XRN2). Together, these findings indicate that the great majority of XAB2-bound protein partners are functionally involved in RNA processing and genome utilization processes. The role of XAB2 in splicing *in vitro* is already well-documented. In support of that spliceosomal factors were highly enriched in the bXAB2-bound proteome of P15 livers. Nevertheless, as splicing factors are often non-specifically enriched in affinity enrichment-mass spectrometry approaches [33], we first challenged the specificity of XAB2 to the pre-mRNA splicing machinery by comparing the bXAB2 MS liver proteome to that derived previously from P15 biotin-tagged xeroderma pigmentosum complementation F (bXPF) livers under similar experimental conditions [22]. XPF is the obligatory interacting partner in the NER structure-specific ERCC1-XPF endonuclease complex [34]. Unlike bXPF, we find that bXAB2 co-immunoprecipitates several core spliceosome factors and more than half of them are proteins involved in the PRP19 and PRP19-related complex (nine out of 16 pre-mRNA splicing factors) (**Figure 2G**; colored in red).

**Figure 2.**
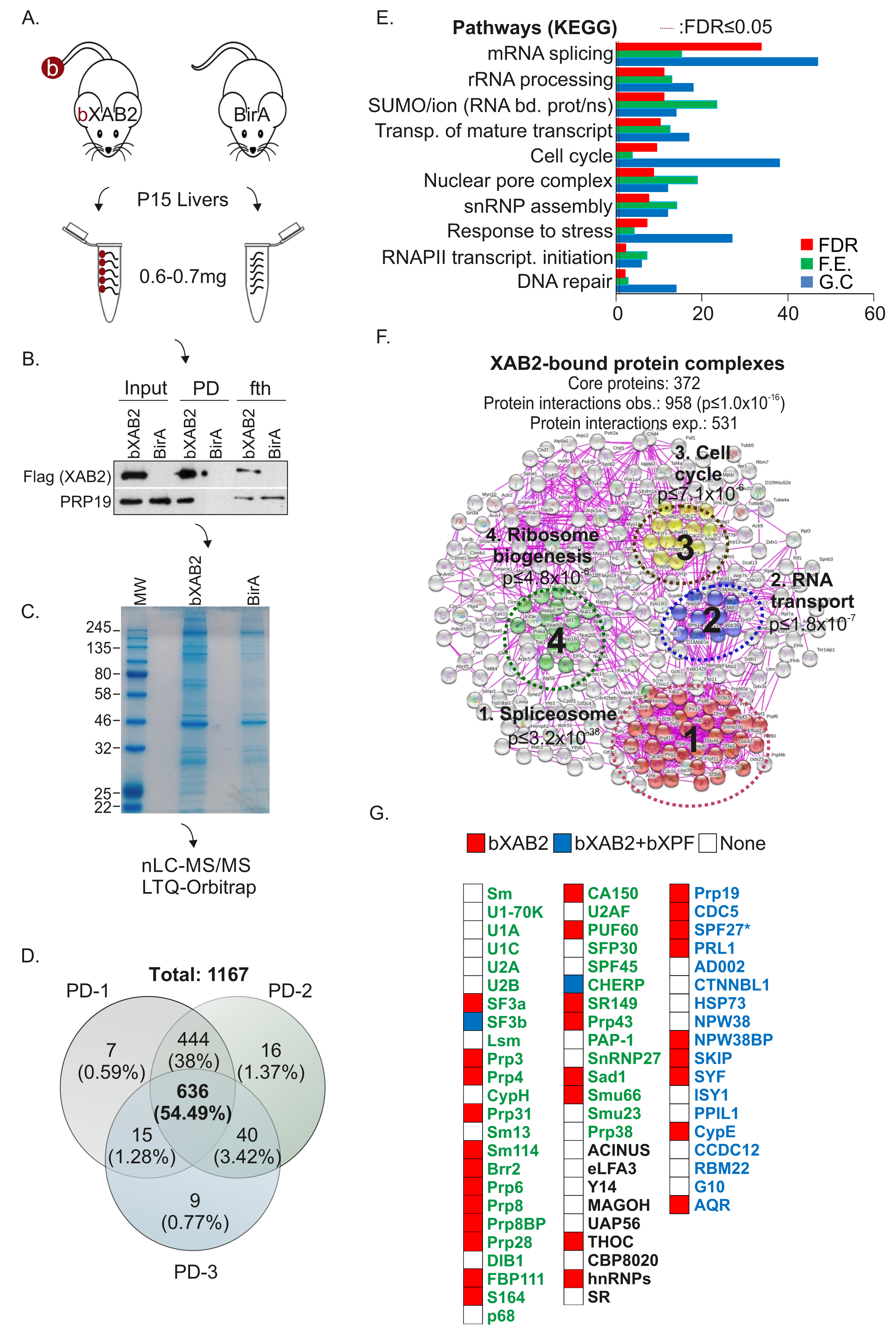
XAB2 interacts with protein complexes involved in transcript maturation. (**A**). Schematic representation of the high-throughput MS analysis in livers derived from P15 bXAB2 animals expressing the BirA transgene (n=8) and BirA transgenic control mice (n=8). (**B**). bXAB2 pulldowns (PD; n PD: Pulldown) and western blot with anti-FLAG (for XAB2) and anti-PRP19 in nuclear extracts derived from P15 bXab2 livers (n=3). (**C**). A representative SDS-PAGE gel of proteins extracts derived from P15 XAB2 and BirA transgenic mice. (**D**). Venn’s diagram of bXAB2-bound protein factors from three independent pull-downs (PD) and subsequent MS analyses; % depicts the percentage of XAB2-bound proteins involved in the indicated biological process over the total number of XAB2-bound proteins (**E**). List of significantly over-represented pathways (KEGG). The –log(FDR), which is calculated by Fisher’s exact test right-tailed, sorts the biological processes. The red-dotted line marks the threshold of significance at FDR ≤ 0.05. Count: the number of identified XAB2-bound protein factors involved in the indicated biological process. F.E. represents the ratio of XAB2-bound proteins involved in a process (sample frequency) to the total genes involved in the process (background frequency). (**F**). Number of observed (obs.) and expected (exp.) known protein interactions within the core 372 XAB2-bound protein set and schematic representation of the four significantly over-represented XAB2-bound protein complexes based on experimental evidence. (**G**) Heat map representation of bXAB2- and bXPF-bound proteins involved in the core spliceosome in P15 bXAB2 and bXPF livers. Constitutive factors of snRNPs are shown in green-colored; the PRP19 and the PRP19-related complex are shown in blue color. SPF27 (BCAS2) is marked with an “*” as it has been identified in two out of the 3 MS runs in P15 bXAB2 livers but has been further confirmed in downstream immunoprecipitation assays. The images shown on Fig. 2B and C are representative of experiments that were repeated more tha three times. Fth: flow-through; FDR: False Detection Rate; F.E: Fold Enrichment; GC: Gene count.

### Depletion of XAB2 leads to defective NER

Previous findings revealed that XAB2 is involved in the transcription-coupled sub-pathway of NER (TC-NER) ^18,29^. To test for the functional role of XAB2 in NER, we carried a series of pulldown assays in nuclear extracts of UV-irradiated (10J/m^2^) and control bXAB2 primary mouse embryonic fibroblasts (MEFs). We find that bXAB2 interacts with the DNA damage-binding protein-1 (DDB1) involved in UV-induced DNA damage recognition ^30^, and with a portion of xeroderma pigmentosum, complementation group A protein (XPA) known to assemble the NER incision complex at sites of DNA damage^31^, in control and UV-irradiated cells (**Figure S1B**). Confocal microscopy in UV-irradiated HEPA cells revealed that XAB2 does not co-localize with the UV-induced CPDs (**Figure S1C**). The great majority of XAB2 is distributed throughout the nucleoplasm but is excluded from the denser DAPI-stained heterochromatin regions or the nucleolus and forms a punctate staining that is distinctively adjacent to the DNA damage marker γH2Ax (**Figure S1D**). Knockdown of XAB2 in si*Xab2* HEPA cells (**Figure S1E**) results in increased cell death (**Figure S1F**), reduced proliferative capacity (**Figure S1G**) at 72h post-transfection and in the presence of irreparable UV-induced CPDs at 24h post-UV irradiation (**Figure 3A**). In agreement with the defective repair of UV-induced CPDs, si*Xab2* cells manifest a noticeable decrease in unscheduled DNA synthesis at 2.5h post-UV irradiation (measuring global genome NER; **Figure 3B**) ^32^. si*Xab2* cells also present with substantial delay in the recovery of RNA synthesis (assessing TC-NER; **Figure 3C)** as evaluated by the fluorescent detection of bromouridine (BrU) labelled nascent RNA at 4h post-UV irradiation. Knockdown of XAB2 reduced RNAPII (**Figure S1H)** but did not affect the protein levels of NER factors DDB1, XPA and ERCC1, (**Figure S1I**). Next, we investigated whether DDB1 and XPA are efficiently recruited at DNA damage sites in UV-irradiated si*Xab2* HEPA cells. We find that DDB1 co-localizes competently with UV-induced CPDs in si*Xab2* cells (**Figure 3D**). In contrast, XPA fails to form visible foci and properly recruit on UV-induced CPDs in these cells (**Figure 3E**). Finally, we find that treatment with Isoginkgetin, a pre-mRNA splicing inhibitor leads to severe splicing defects as documented by the accumulation of pre-mRNAs and the decrease of corresponding mRNA levels, (**Figure S1J- K)** but has no significant impact on the repair of UV-induced CPDs in HEPA cells indicating that splicing *per se* does not affect NER.

**Figure 3.**
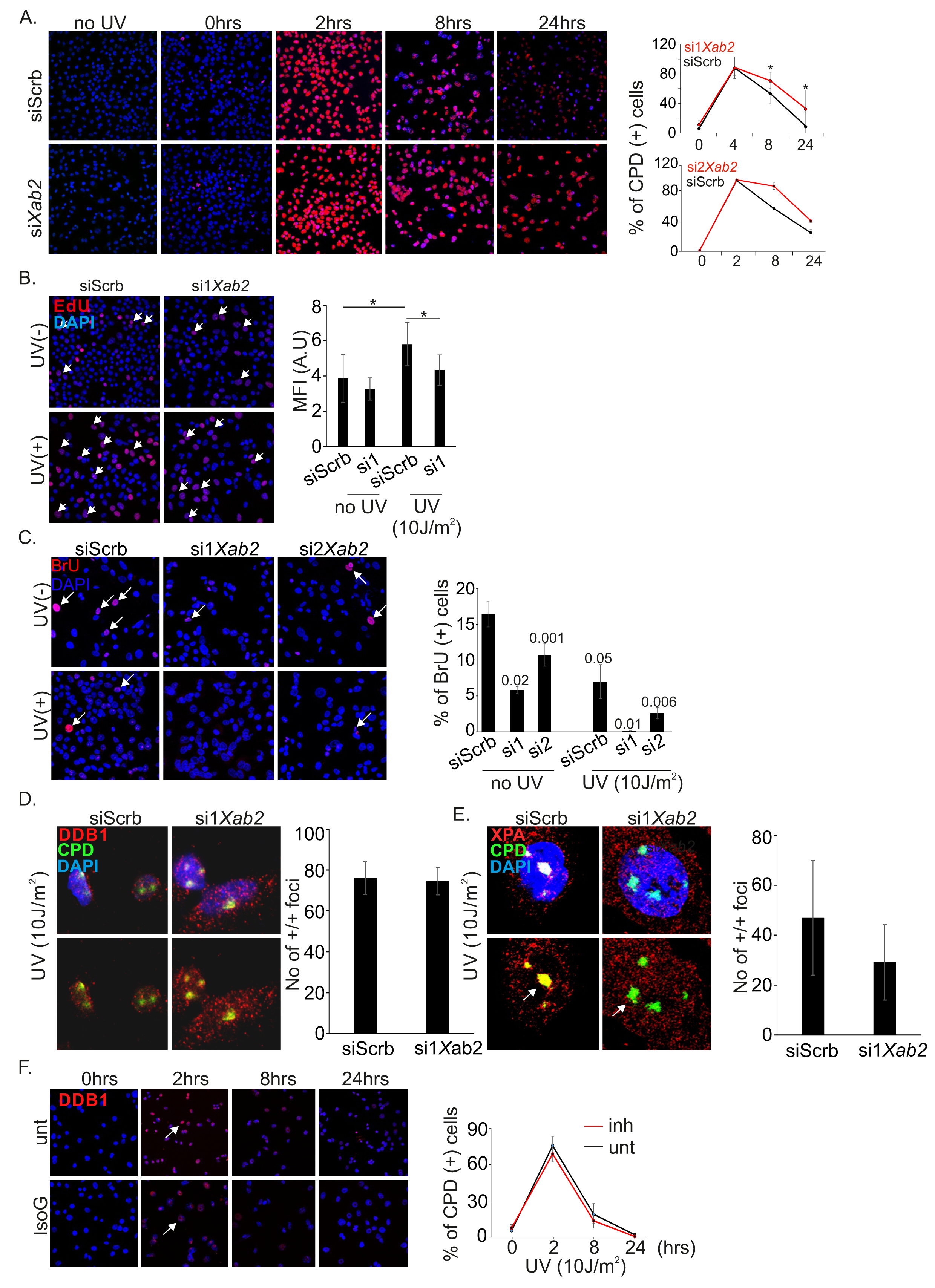
**XAB2 is required for NER A**. Immunofluorescence detection and quantification of CPDs in *siScrb* and si1 and si2*Xab2* treated cells at 4, 8 and 24h post-UV irradiation (10J/m^2^) (**B**). Immunofluorescence detection of unscheduled DNA synthesis in siScrb and si*Xab2* cells treated with 5-Ethynyl-2ʼ-deoxyuridine (EdU) at 48h post-transfection and 2.5h post UV irradiation (10J/m^2^). (**C**). Immunofluorescence detection of RNA synthesis recovery in siScrb and si*Xab2* cells treated with 5-Bromouridine (BrU) at 48h post-transfection with or without UV irradiation (10J/m^2^). (**D**). Immunofluorescence analysis of DDB1 and CPDs in siXab2 and siScrb cells exposed to UV irradiation (10J/m2). (**E**). Immunofluorescence detection of XPA and CPDs in siScrb and si*Xab2* cells exposed to UV irradiation (10J/m2). (**F**) Immunofluorescence detection and quantification of CPDs in untreated and isoginketin treated cells at 4, 8 and 24h post-UV irradiation (10J/m2). The asterisk “*” indicates a p-value ≤ 0.05 (n ≥ 3 per time point, treatment or genotype).

### XAB2 is part of a core spliceosome complex that binds on UsnRNAs

Follow-up pulldown experiments confirmed the mass spectrometry interactions and revealed that the endogenous bXAB2 interacts with Aquarius (AQR), PRP19 and BCAS2, an integral component of the spliceosome that is required for activating pre-mRNA splicing ^33^ in P15 livers (**Figure 4A**). The interaction of bXAB2 with core spliceosome factors in P15 livers prompted us to test whether XAB2 binds RNA *in vivo*. To do so, we carried out a series of RNA immunoprecipitation (RIP) experiments in P15 bXAB2 liver nuclear extracts incubated with streptavidin-coated beads (for bXAB2). The co-precipitated RNA was eluted, reversely transcribed and quantified for spliceosomal UsnRNAs by quantitative (q)PCR. We find that bXAB2 shows low specificity for U1 snRNA and no association with U2 snRNA or U3 small nucleolar RNAs (snoRNA). However, bXAB2 binds consistently to U4, U5 and U6 snRNAs (**Figure 4B**). Using U4 and U6 snRNAs in follow-up experiments, we next carried out a series of native RIP assays in nuclear extracts of HEPA cells. Unlike with XAB2 and AQR, we find that BCAS2 fails to recruit on U4 and U6 snRNAs (**Figure 4C**), indicating its indirect association with these targets. RNA-Seq analysis in si*Xab2* HEPA cells and mESCs 48 hrs post-transfection revealed 255 differentially expressed genes (124 genes were upregulated; 131 downregulated genes; fold change: +/-1.2, q-value ≤ 0.05) (**Table S3**) in si*Xab2* HEPA cells and 333 differentially expressed genes (41 upregulated and 291 downregulated; fold change: +/-1.2, q-value ≤ 0.05) (**Table S4**) in si*Xab2* mESCs, with a striking overrepresentation of ribosomal protein genes. Using this dataset, we next tested whether knockdown of *Xab2* has an effect on splicing decisions i.e. exon skipping (ES), intron retention (IR) and alternative selection of 5’ or 3’ splice sites (**Figure 4D**). Of the 126335 and the 56677 alternative spliced (AS) events detected in HEPA cells and mESCs respectively, 774 and 1134 were found differentially spliced in si*Xab2* cells (probability of differential splicing >0.8 and an absolute change in percent spliced-in (ΔPSI) greater than 5%). We find that the great majority (i.e. 805 out of 849, 94.82%, p≤10^-16^) of IR events with a significant probability of differential splicing are retained in si*Xab2* cells. Moreover 58,03% of alternative exons (i.e. 430 out of 741, p ≤ 10^-5^) are skipped when *Xab2* is ablated, indicating a global impairment of the splicing machinery in si*Xab2* cells (**Figure 4E-H**, **Figure S1M, Figure S2A-I, Table S5,** and **Table S6**).

**Figure 4.**
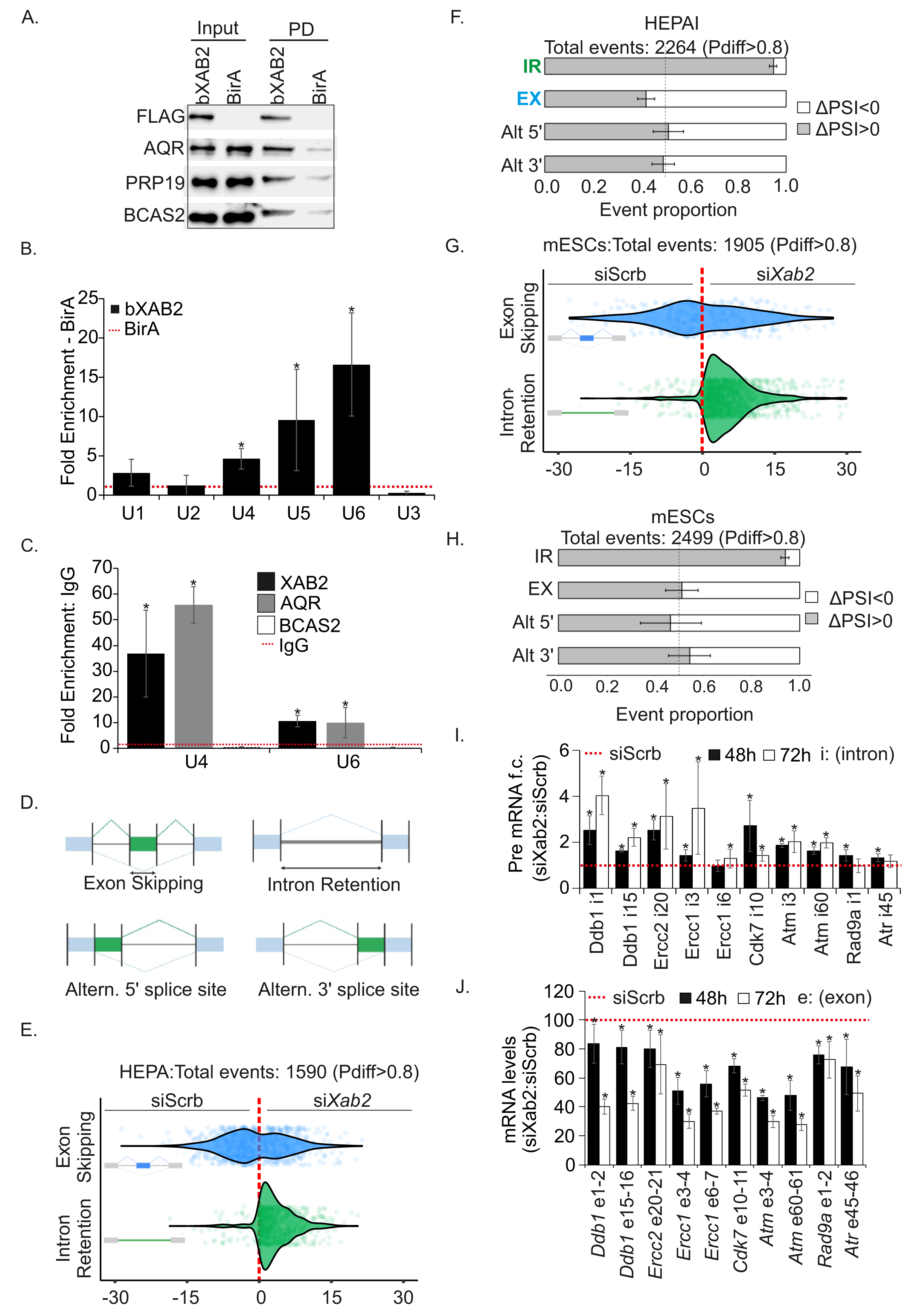
XAB2 is part of a core spliceosome complex. (**A**). bXAB2 pulldowns and western blot analysis of AQR, FLAG-tagged XAB2 (FLAG), PRP19 and BCAS2 proteins in P15 bXAB2 livers. (**B**). RNA pulldowns with bXAB2 and BirA (control) on U1-U6 U sn(o)RNAs (as indicated). (**C**). RNA immunoprecipitation with XAB2 and BCAS2 on U4 and U6 snRNAs (as indicated). (**D**). Schematic representation of splicing decisions i.e. exon skipping, intron retention and alternate (Altern.) selection of 5’ or 3’ splice sites. (**E and G**). Graph depicting the differences in Percent Spliced-In (PSI) between si*Xab2* and siScrb HEPA and mESCs cells respectively (ΔPSI > 0: increased alternative exon/intron inclusion in *siXab2* and ΔPSI < 0: decreased alternative exon/intron inclusion in *siXab2* compared to siScrb cells. (**F and H**). Distribution of the direction of alternative splicing changes between si*Xab2* and siScrb HEPA and mESCs cells respectively (**I**). Pre-mRNA levels and (**J**) mRNA levels of transcripts with retained introns in si*Xab2* and siScrb HEPA cells at 48h and 72h post-transfection (as indicated). The images shown on Fig. 3A, 3E, 3G are representative of experiments that were repeated more than three times. The asterisk “*” indicates a p-value ≤ 0.05 (n ≥ 3 per time point, treatment or genotype). FDR: False Detection Rate; F.E: Fold Enrichment; GC: Gene count.

To investigate which biological processes are mostly affected by XAB2 knockdown-mediated AS, we subjected the AS events found to GO classification. At the confidence level set in our analysis (FDR ≤ 0.05), we find that biological processes related, among others, to cell cycle, RNAPII-mediated transcription, DNA repair and RNA processing comprise a disproportional number of transcripts in both mESCs and HEPA cells relative to the murine transcriptome (**Figure S3A, S3B**). RNA immunoprecipitation in P15 livers revealed that XAB2 binds to the pre-mRNAs of several genes involved in e.g. DNA repair and the DDR (**Figure S3C**) as well as to the pre-mRNAs of genes known to be highly transcribed during postnatal hepatic development (e.g. *Ghr*, *Igf1*) (**Figure S3D**). Transcripts with retained introns often contain premature stop codons (PTCs), which leads to their cytoplasmic degradation by the nonsense-mediated RNA decay machinery ^34,35^ or, as in the case of XAB2-mediated intron retained transcripts to aberrant RNA-decay pathways ^25^. Otherwise, a subset of these transcripts may be stably retained within the nucleus for later processing ^36^. Beginning at 48h post-transfection, we find that the pre-mRNA levels of transcripts with retained introns gradually accumulate in HEPA cells depleted for XAB2 with two distinct siRNAs (**Figure 4I** and **Figure S3E**) whereas their corresponding mRNA levels progressively decrease (**Figure 4J**) indicating a defect in splicing rather than IR regulation in si*Xab2* cells. Thus, XAB2 is part of the core spliceosome complex and has a functional role in pre-mRNA splicing; it is recruited on UsnRNAs and pre-mRNAs and it is required for proper mRNA splicing early on during mammalian development.

### XAB2 is released from UsnRNAs and pre-mRNAs upon DNA damage and transcription blockage

UV-induced DNA lesions trigger the displacement of co-transcriptional mature catalytically active spliceosome (U2, U5 and U6) ^11^. In our work, we find that UV-induced DNA damage also triggers the rapid release of XAB2 and AQR from U4 and U6 snRNAs for at least 6 hours affecting the premature, catalytically active spliceosome (**Figure 5A**) and of XAB2 from all pre-mRNAs tested (**Figure 5B**). Importantly, the release of XAB2 from UsnRNAs is not accompanied by its disassociation from other protein components of the spliceosome machinery i.e. PRP8, PRP19, BCAS2, and SNRP40 (**Figure 5C**) and is consistent with the substantial increase in intron retention for several transcripts tested in UV-irradiated cells (**Figure S3F**). In contrast to the ATM-dependent UV-induced dis-association of mature spliceosome from nascent transcripts ^11^, selective inhibition of ATM (ATMi) with KU-55933 inhibitor ^37^ or of ATR (ATRi) with NU6027 in cells treated with UV did not rescue the release of XAB2 from U4 and U6 snRNAs (**Figure 5D**). These findings indicate that the DNA damage-driven release of XAB2 from RNA is not mediated through active DDR signaling. Next, we reasoned that DNA lesions altering the DNA double helix itself and/or interfering with transcription *in cis* are causal to the aberrant release of XAB2 from UsnRNAs or the pre-mRNAs. To test this, we made use of mouse embryonic fibroblasts (MEFs) derived from CPD photolyase transgenic mice ^38^. Photolyases directly revert photolesions into undamaged bases using visible light energy (i.e. photoreactivation; PR) and display substrate specificity for UV-induced 6-4PPs or CPDs ^39^. Using this system, CPD photolyase transgenic MEFs were exposed to UV irradiation and were subsequently kept in the dark (UV-induced CPDs remain in the mammalian genome) or exposed to PR light (UV-induced CPDs are repaired) for 60min post-UV irradiation (**Figure 5E**). Importantly, we find that the exposure of UV-irradiated MEFs to PR light substantially rescues the XAB2 RIP signals in U4 and U6 snRNAs (**Figure 5F**) and in pre-mRNAs tested (**Figure 5G**) indicating that the presence of DNA helix-distorting lesions inadvertently affects the process of pre-mRNA splicing in the mammalian genome. Likewise, we find an aberrant release of XAB2 from UsnRNAs in MEFs treated with illudin S known to induce DNA lesions that are ignored by global-genome NER and are exclusively processed by transcription- and replication-coupled repair pathways (**Figure 5H**) ^40^. The latter prompted us to test whether XAB2 binding to U4 and U6 snRNAs is also altered in *Csb^m/m^* mice carrying an inborn defect in TC-NER ^41^. Using P15 *bXab2*/*Csb^m/m^* livers, we find that the XAB2 RIP signals on U4 and U6 snRNAs as well as in the highly transcribed pre-mRNAs (*Ghr*, *Igf1*) are substantially decreased in the P15 livers of *bXab2*/*Csb^m/m^* animals (**Figure 5I** and **Figure S3G**).

**Figure 5.**
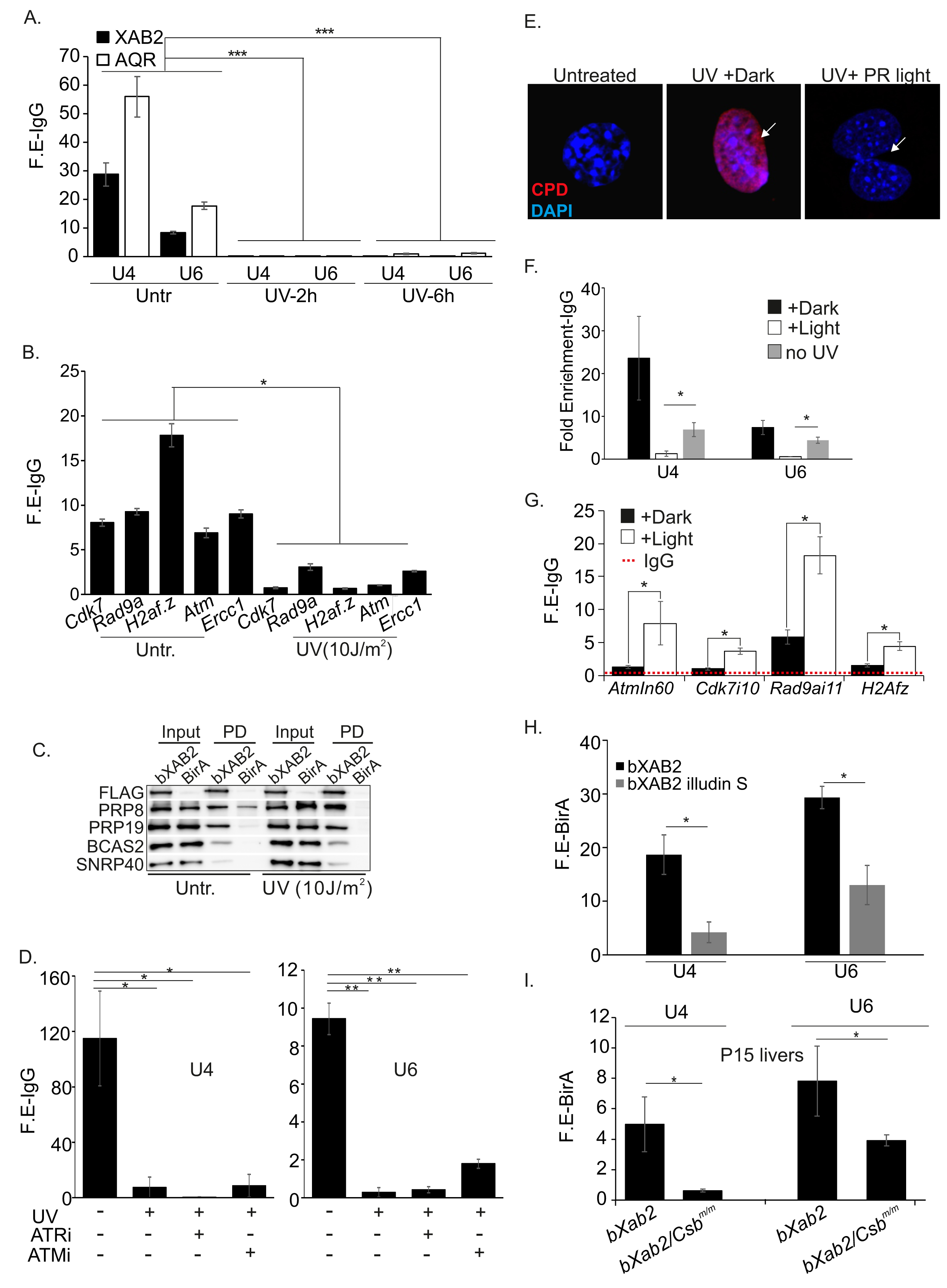
XAB2 is released from U snRNPs and pre-mRNAs upon DNA damage. (**A**). RNA immunoprecipitation of XAB2 and AQR on U4 and U6 snRNAs at 2 and 6h post-UV irradiation. (**B**). RNA immunoprecipitation of XAB2 on pre-mRNAs in untreated and UV-irradiated HEPA cells (as indicated). (**C**). bXAB2 pulldowns followed by western blot analysis with FLAG (for bXAB2), PRP8, PRP19, BCAS2 and SNRP40 in UV-irradiated (10J/m^2^) and control MEFs. (D). RNA immunoprecipitation of XAB2 on UsnRNAs in untreated, UV-irradiated cells and UV-irradiated cells pre-treated with inhibitors against ATM (ATMi) and ATR (ATRi; as indicated). (**E**). Immunofluorescence detection of UV-induced CPDs in UV-irradiated photolyase transgenic MEFs exposed to 1 hour of PR light (CPDs are repaired) or dark (CPDs remain) as indicated. (**F**). RNA immunoprecipitation of XAB2 on U4 and U6 snRNAs in UV-irradiated photolyase transgenic MEFs exposed to 1 hour of PR light (CPDs are repaired) or dark (CPDs remain) (**G**). RNA immunoprecipitation of XAB2 on pre-mRNAs in UV-irradiated photolyase transgenic MEFs exposed to 1 hour of PR light (CPDs are repaired) or dark (CPDs remain). (**H**) bXAB2 RNA pull downs on U4 and U6 snRNAs in untreated or illudin S-treated birA and bXAB2 MEFs. (**I**) bXAB2 RNA pull downs on U4 and U6 snRNAs in P15 bXAB2 or bXAB2;*Csb^m/m^* mouse livers (n ≥ 5 per genotype). The asterisk “*” indicates a p-value ≤ 0.05 (n ≥ 3 per time point, treatment or genotype).

### Abrogation of XAB2 promotes R-loop formation triggering DNA damage

R-loops are generated during transcription when nascent RNA exits RNA polymerase and pairs with its complementary DNA template to form a stable RNA–DNA hybrid that displaces single-stranded DNA (ssDNA) ^42^. Increased formation of R-loops is frequently observed in cells that are defective in DNA repair or RNA processing ^43-45^. In UV-irradiated MEFs, a DNA-RNA immunoprecipitation (DRIP) approach followed by treatment with RNase H (RNH; it digests RNA in DNA-RNA hybrids eliminating R-loops) revealed an enrichment of R-loops on *Cdk7* and *Ercc1* genetic loci whose pre-mRNAs were previously shown to remain bound by XAB2 (**Figure 6A**). Inhibition of ATM led to a detectable yet mild decrease in the formation of R-loops indicating that defective DDR signaling alone cannot fully justify the observed accumulation of R-loops in UV-irradiated cells (**Figure 6A**). This and our finding that R-loops accumulate substantially in MEFs treated with isoginketin (**Figure 6B**, **Figure S3G**) prompted us to further explore the functional role of XAB2 in R-loop processing. Similar to UV or isoginketin treatment in MEFs (**Figure 6A-B**), we find that knockdown of XAB2 triggers the formation of R-loops in cells; importantly, treatment of cells with RNase H post-fixation led to a substantial decrease in R-loops confirming the validity of these findings (**Figure 6C, Figure S4A**).

**Figure 6.**
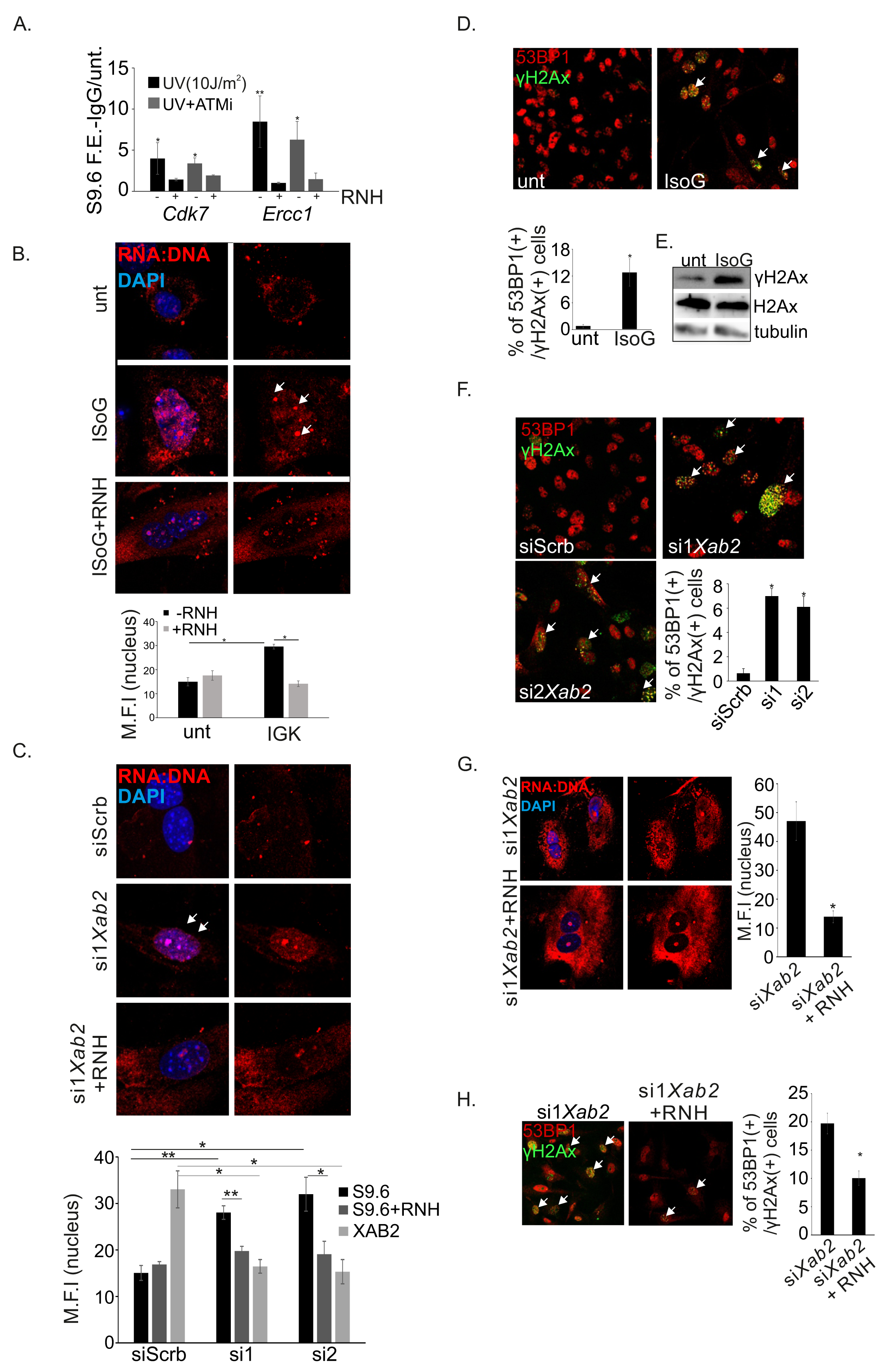
Abrogation of XAB2 leads to R-loop dependent genomic instability. (**A**) DRIP with an antibody against RNA:DNA hybrids on *Cdk7* and *Ercc1* gene loci in untreated and UV-irradiated MEFs treated or not with RNaseH1 (RNH) and pretreated with ATM inhibitor (ATMi), as indicated. (**B**). Representative immunofluorescence images and quantification of RNA:DNA hybrids (indicated by the white arrows) in untreated and isoginketin (IsoG)-treated wt. MEFs (see supplemental Figure S3H for lower magnifications). The graph depicts the mean S9.6 fluorescence intensity per nucleus (≥150 nuclei were analyzed per condition) from three representative experiments. (**C**) Immunofluorescence detection of R loops in siScrb and si*Xab2* MEFs treated (or not) with RNaseH1 (RNH). See supplemental Figure S4A for lower magnifications and representative images for the efficiency of Xab2 silencing in si2*Xab2* cells. The graph depicts the S9.6 fluorescence intensity per nucleus (≥150nuclei were analyzed per condition) from three representative experiment (**D**) Immunofluorescence detection of γ AX and 53BP1 (white arrowheads) in untreated and isoginketin (IsoG)-treated wt. MEFs. The graph represents the percentage of γH2AX+ 53BP1+ cells from three representative experiments (≥150nuclei were analyzed per condition). (**E**) Western blot analysis of phosphorylated and total H2AX levels in untreated and isoginketin (IsoG)-treated MEFs. (**F)** Immunofluorescence detection of γH2AX and 53BP1 (white arrowheads) in siScrb and si*Xab2* MEFs. The graph represents the percentage of γH2AX+ 53BP1+ cells in siScrb and siXab2 primary MEFs (see supplemental Figure S4A for the efficiency of *Xab2* silencing). (**G)** Immunofluorescence detection of RNA:DNA hybrids in si*Xab2* MEFs with or without RNaseH1 (RNH) protein transfection. The graph represents the S9.6 fluorescence intensity per nucleus (≥150 nuclei were analyzed per condition) from three representative experiments. (H) Immunofluorescence γH2AX and 53BP1 (white arrowheads) in *siXab2* MEFs with or without RNaseH1 (RNH) protein transfection. The graph represents the percentage of γH2AX+ 53BP1+ cells from three representative experiment (≥150 nuclei were analyzed per condition). The asterisk “*” indicates a p-value ≤ 0.05.

Treatment of cells with isoginketin also led to the substantial increase of γΗ2 x (+) 53BP1 (+) cells (**Figure 6D**) as well as to higher γH2Ax protein levels compared to untreated controls (**Figure 6E)**, indicating that a defect in splicing leads to persistent DNA damage accumulation. Likewise, RNAi-mediated ablation of XAB2 with two distinct siRNAs increased substantially the number of γΗ2Ax x (+) 53BP1 (+) cells (**Figure 6F**). As R-loops often lead to persistent DNA breaks, we next tested whether genome instability in si*Xab2* cells is a result of R-loop accumulation. Immunofluorescence studies revealed that protein transfection of RNase H in si*Xab2* cells leads to a decrease in R-loops (**Figure 6G)** and in the number of γΗ2 x (+) 53BP1 (+) cells (**Figure 6H)**. Consistent with our previous DRIP analysis on UV-irradiated cells (**Figure 6A**), confocal microscopy studies revealed that UV irradiation triggers the formation of R-loops in wt. cells that accumulate even further when XAB2 is silenced (**Figure S4B**); the latter leads to the substantially increase of γΗ2 x (+) 53BP1 (+) cells (**Figure S4C)**.

### XAB2 interacts with R-loop processing factors and is recruited to RNA:DNA hybrids under conditions that favor R-loop formation

Highly transcribed loci are prone to R-loop formation ^46,47^. To explore the direct association of XAB2 with R-loops and R-loop processing factors, primary MEFs were treated with all-trans-retinoic acid (tRA), a pleiotropic factor known to activate transcription during cell differentiation and embryonic development ^48^. In agreement, we find that transcription activation in tRA-treated MEFs leads to the substantial accumulation of R-loops in these cells, which becomes further pronounced when cells are treated with tRA and illudin S (**Figure 7A**). DRIP analysis with primers spanning the promoter or 3’end regions of genes (**Figure 7B**) where the frequency of RNA-DNA hybrids is expected to be higher revealed the significant accumulation of R-loops in tRA-induced gene targets i.e. *Rarb2*, *Stra6* and *Fibin* but not in the tRA non-inducible *Chordc* gene (**Figure 7B-C**). Using a series of ChIP assays, we coding region of the tRA non-inducible *Chordc1* gene (**Figure S4D**). Similar to the DNA damage-driven release of XAB2 from RNA targets (**Figure 6A-B**), XAB2 ChIP signals are substantially reduced when primary MEFs are treated with illudin S accumulating transcription-blocking DNA lesions. To confirm that XAB2 and R-loops co-occupy the tRA-responsive *Stra6* gene and the tRA non-inducible *Chordc* gene, we next employed a sequential native bioXAB2 pulldown approach followed by S9.6 DRIP in tRA- and illudin S-treated MEFs. This strategy revealed that bXAB2 is recruited preferentially on the 3’ end region of *Stra6* gene compared to the tRA non-inducible *Chordc* gene in tRA-treated MEFs and that bioXAB2/DRIP signals are higher when cells are treated with illudin S (**Figure 7D**). Importantly, we find that transfection of illudin S-treated MEFs with recombinant RNaseH restores the release of XAB2 from UsnRNAs (**Figure S4E**). R-loops are known to be actively processed by the NER structure-specific endonucleases XPF and XPG ^49,50^. PRP19 may act as a sensor of RPA-ssDNA moieties that emerge naturally when R-loops are formed during ongoing transcription ^21^. Using a series of bXAB2 pulldown and S9.6 DRIP assays under basal transcription conditions or upon tRA treatment in primary MEFs, we find that XAB2 interacts with XPF, XPG and PRP19 proteins (**Figure 7E**, **Figure S4F**) and that the XAB2 complex is recruited on R-loops (**Figure 7F**, **Figure S4G**). Unlike with PRP19, we find that the interaction of XAB2 with XPG or XPF is abolished when cells are treated with RNase H, indicating that the XAB2-XPF-XPG complex requires the presence of R-loops (**Figure 7F**, **Figure S4G**). Moreover, we find that XPF and XPG do not bind R-loops when XAB2 is silenced (**Figure 7G**) or when MEFs are exposed to UV irradiation; the latter indicates the high affinity of XPF and XPG endonucleases for UV-induced DNA lesions (**Figure 7H**). Since the recruitment of XPF and XPG on R-loops requires XAB2 and both endonucleases were previously shown to process R-loops into DNA breaks ^50,51^, it is surprising that *Xab2* knockdown leads to an increase in the number of γH2AX+ 53BP1+ cells. However, these findings could also reflect the known role of XAB2 in HR ^24^ or the fact

**Figure 7.**
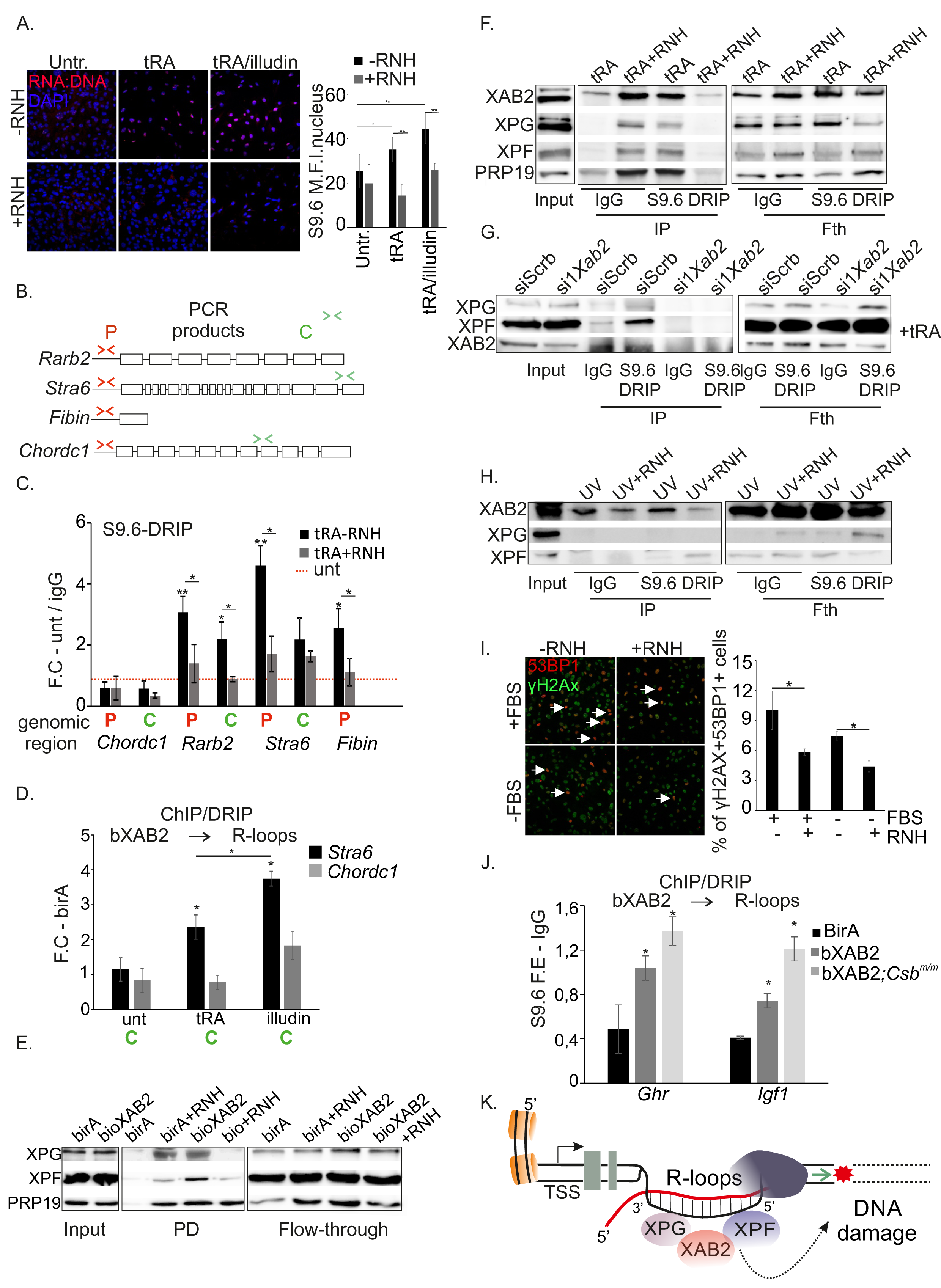
High transcription promotes XAB2 interaction with R-loops. (**A**) Immunofluorescence detection of R loops in untreated, tRA-treated (4 hours) and illudin S-treated (1.5 hours) MEFs. The graph depicts the S9.6 fluorescence intensity per nucleus (≥150 nuclei were analyzed per condition) from three representative experiment. (**B-C**) DRIP analysis of tRA-responsive (*Rarb2, Fibin* and *Stra6)* and non-responsive (*ChordC)* gene targets using the S9.6 antibody against RNA-DNA hybrids with or without RNaseH1 (RNH) treatment in untreated and tRA-treated MEFs. Amplification of promoter (red-colored; A) and coding (green-colored; B) regions are indicated. (**D**) Sequential native ChIP followed by DRIP on the coding region of tRA-induced *Stra6* gene and the tRA-non induced *Chordc1* gene in tRA and illudin S/tRA-treated birA and bXAB2 MEFs (**E**). bXAB2 pulldowns (PD: Pulldown) and western blot with anti-XPF, anti-XPG and anti-PRP19 in nuclear extracts derived from tRA-treated bXAB2 and BirA MEFs with or without RNaseH1(RNH) treatment. (**F**). S9.6 immunoprecipitation followed by western blot analysis for XAB2, XPG, XPF and PRP19 protein levels in the nuclear extracts of tRA-treated MEFs with or without RNaseH1 (RNH) treatment (**G**) S9.6 immunoprecipitation followed by western blot analysis for XPG, XPF and XAB2 protein levels in the nuclear extracts of tRA-treated siScrb or si1*Xab2* MEFs with or without RNaseH1 (RNH) treatment (**H**). S9.6 immunoprecipitation followed by western blot analysis for XPG, XPF and XAB2 protein levels in the nuclear extracts of UV-irradiated (10J/m^2^) MEFs with or without RNaseH1 (RNH) treatment. (**I**). Immunofluorescence detection of γH2AX and 53BP1 (white arrowheads) in wt. MEFs cultured in the presence (FBS+) or absence (FBS-) of fetal bovin serum, transfected (RNH+) or not (RNH-) with RNaseH1 protein. The graph represents the number of γH2AX+ 53BP1+ cells from three representative experiments (≥150 nuclei wer analyzed per condition). The asterisk “*” indicates a p-value ≤ 0.05 (**J**). S9.6 DRIP analysis with or without RNaseH1 (RNH) treatment on *Ercc1* and *Cdk7* gene targets in P15 wt. and *Csb^m/m^* mouse livers (n ≥ 3 per genotype). (**K**). A working model for XAB2 role in DNA repair, mRNA and R-loop processing. Under conditions that favor transcription, the splicing factor XAB2 interacts with components of the core spliceosome and is recruited on RNA targets for pre-mRNA processing. DNA damage inhibits RNA synthesis and triggers the release of XAB2 from snRNAs and pre-mRNA targets leading to aberrant intron retention and the accumulation of R-loops. The splicing factor XAB2 interacts with ERCC1-XPF and XPG endonucleases and the complex is recruited on RNA:DNA hybrids, thereby linking R-loop processing with the spliceosomal response to transcription-blocking DNA lesions during hepatic development. The asterisk “*” indicates a p-value ≤ 0.05 (n ≥ 3 per time point, treatment or genotype).

that the accumulation of R-loops in si*Xab2* cells could spontaneously lead to DNA breaks due to transcription-DNA replication conflicts in dividing cells ^52^. In support, we find that although the number of R-loops is comparable in tRA-treated MEFs cultured under serum-starved or serum-rich media, the number of γH2AX+ 53BP1+ cells (due to R-loops alone), is reduced in serum-starved MEFs compared to proliferating corresponding control cells (**Figure 7I**, **Figure S4H**, **Figure S4I**). To test whether XAB2 interaction with XPF and XPG on R-loops requires functional NER, we also performed S9.6 DRIP in wt. and totally NER-defective *Xpa^-/-^* MEFs (**Figure S5A-B**; as indicated) followed by western blotting for XPA (for wt cells), XPF and XPG. We find that XPA does not associate with R-loops and that recruitment of NER endonucleases on R-loops does not require XPA. Nevertheless, we find that R-loops accumulate in tRA-treated *Xpa^-/-^* MEFs compared to wt. controls (**Figure S5C**) likely reflecting the contributing role of irreparable DNA lesions (owing to the NER defect in *Xpa^-/-^* cells) on R-loop accumulation. In support, we find that an enrichment of R-loops on the promoters of highly transcribed *Igf1* and *Ghr* genes in P15 *Csb^m/m^* livers compared to littermate controls (**Figure S5D**). Similar to the reduction of XAB2 ChIP signals in illudin S-treated primary MEFs accumulating transcription-blocking DNA lesions (**Figure S4E**) we find that bXAB2 ChIP signals on the promoters of *Ghr* and *Igf1* genes are substantially reduced in P15 bXab2/*Csb^m/m^* livers but in age-matched bXAB2 controls (**Figure S5B**). Finally, a sequential native bioXAB2 pulldown followed by S9.6 DRIP in P15 bXAB2 and bXAB2/*Csb^m/m^* livers revealed that XAB2 interacts with R-loops on the promoters of *Igf1* and *Ghr* genes *in vivo*. Thus, XAB2 interacts conditions that promote R-loop formation (**Figure 7J)**.

## Discussion

Inborn defects in RNA processing or DNA repair associate with progeria ^53-55^, the metabolic syndrome ^56-58^, neurodegeneration ^59-62^ and cancer ^63-67^, arguing for functional links between genome maintenance and the splicing machinery in development or disease. Using an *in vivo* biotinylation tagging approach in mice, we find that XAB2 interacts with components of the core spliceosome as well as with DDB1 involved in UV-induced DNA damage recognition and XPA that plays a prominent role in the assembly of the NER incision complex at sites of DNA damage ^62,68^. As DDB1 is successfully recruited to UV-induced CPDs in si*Xab2* cells, the implication of XAB2:DDB1 interaction remains elusive. However, in si*Xab2* cells, XPA fails to form visible foci and to be recruited to UV-induced DNA lesions. In support of a defect in NER, si*Xab2* cells have decreased UDS and delayed RNA synthesis recovery following UV-induced DNA damage. Consistently, UV-induced CPDs remain unrepaired even 24h post-UV irradiation in si*Xab2* cells. Arguably, the defect in splicing does not interfere with the DNA repair defect in si*Xab2* cells because exposure of cells to a pre-mRNA splicing inhibitor has no detectable impact on the repair kinetics of UV-induced CPDs. We find that XAB2 is recruited to UsnRNAs and pre-mRNAs and that it is released from all RNA targets tested when cells are treated with UV-induced DNA damage or with the genotoxin illudin S. Intriguingly, the release of XAB2 from RNA following DNA damage is not rescued when ATM or ATR are inhibited but only when UV-induced CPDs are efficiently removed through PR-dependent repair. The latter supports previous observations showing that DNA damaging agents affect the subcellular localization of splicing factors or their association with nascent transcripts ^69-72^ and that ATM (and likely ATR) function downstream of the spliceosome displacement ^11^.

Nascent pre-mRNAs that associate with transcription complexes blocked at DNA lesions are released from late-spliceosome components and hybridize to the template strand of the melted DNA to form RNA:DNA duplexes (R-loops), a structure that slows transcription and activates DDR ^72^. Increased formation of R-loops is frequently observed in cells that are defective in DNA repair or RNA processing ^43-45^ leading to DNA breaks ^50^, transcription stalling and DDR activation ^72^. In support, we find that R-loops accumulate in primary MEFs following DNA damage or upon treatment with a pre-mRNA splicing inhibitor. Knockdown of *Xab2* leads to the substantial increase in R-loop formation and to an R-loop-dependent accumulation of X and 53BP1 in cells. Indeed, the number of γΗ2 X (+) 53BP1 (+) cells is significantly reduced when si*Xab2* cells are protein-transfected with RNaseH1 known to eliminate R-loops. Using a series of immunoprecipitation strategies, we find that XAB2 interacts with XPG and XPF endonucleases and that the XAB2 is required for the trimeric protein complex to be recruited on RNA:DNA hybrids under conditions that favor R-loop formation. The latter supports previous observations indicating that the NER endonucleases XPF and XPG actively process unscheduled R-loops into DNA breaks ^50,73^. The interaction of XAB2 with XPG or XPF as well as the recruitment of the XAB2 complex on RNA:DNA hybrids are also observed in NER-defective *Xpa^-/-^* MEFs, indicating that R-loop processing does not require functional NER. Similar to RNA, transcription-blocking DNA lesions trigger the release of XAB2 from DNA sites. However, we find that bXAB2 ChIP signals followed by DRIP are substantially higher on the 3’ end region of the tRA-inducible *Stra6* gene and increase even further when cells are also treated with illudin S. Likewise, DRIP signals followed by bXAB2 ChIP are higher on the R-loop-enriched promoter regions of *Igf1* and *Ghr* genes in P15 *Csb*^m/m^ livers indicating that XAB2 preferentially binds on DNA sites that associate with RNA. Taken together, our findings suggest that the splicing factor XAB2 is released from UsnRNAs and pre-mRNAs and together with XPG and XPF, it is recruited on RNA:DNA hybrids under conditions that promote R-loop formation *e.g.* transcription induction or DNA damage (**Figure 7K**). The finding that XAB2 is released from RNA targets in progeroid *Csb^m/m^* developing livers makes it attractive to test whether DNA damage-driven changes in RNA processing are causal to the premature onset of age-related pathological symptoms seen in TC-NER progeroid syndromes and during natural aging.

## Methods

### Generation of biotin-tagged XAB2 animals

To generate the targeting vector for the insertion/knock-in of the Avi tag cassette before the stop codon of the last exon of the XAB2 gene for the generation of the avXAB2 knock-in mice, PCR products were first amplified using Phusion High-Fidelity DNA Polymerase (NEB). The avi-tag was sub cloned in pBSSK (EcoRI/Hind III 0.18-kb). A triple ligation reaction was set up using the fragments: 5’ homology sub cloned in two fragments (XbaI/BamHI 2.4kb and BamHI/ExoRI 1.2kb); avi tag (EcoRI/Hind III 0.18-kb); pBSSK-avi tag (XbaI/ExoRI 2.9kb). The lox-neomycin-lox cassette (HindIII/SalI 1.5-kb) was subsequently cloned into the vector followed by cloning of the 3’ homology region (SalI fragment 3.3kb). Finally, the MC1-TK gene (SacII 1.8-kb) was inserted into the vector for negative selection. The final targeting vector was linearized using NotI and used for embryonic stem cell electroporation. 129/SV embryonic stem cells carrying the Protamine 1-Cre transgene were maintained in their undifferentiated state (LIF-ESGRO 10^7^ units) and grown on a feeder layer of gamma-irradiated (3.500 rads) G418^r^ primary mouse embryonic fibroblasts. Electroporation (400V, 25 μF) of 0.8x10^7^ embryonic stem cells with 50 μg of NotI linearized targeting vector (2mg mL^-1^) was performed and homologous recombined clones were selected with G418 (300μg mL^-1^) and ganciclovir (2μM). G418-resistant embryonic stem cell clones were subjected to Southern blot analysis and hybridized with 5’ and 3’ probes from their homology region. Genomic DNA from embryonic stem cell clones was digested overnight with EcoRV (MINOTECH Biotechnology) and resolved on 1% agarose gels. Samples were immobilized on Hybond-NC nylon membranes (Amersham Bioscience) and hybridized with probes with [^32^P] dCTP (Izotop). 5’ (1.1kb NcoI/EcoRI) and 3’ (1.2-kb BglII/Hind III) specific probes flanking the last exon of the Xab2 gene were used to identify the targeted (6.4-kb or 7.5-kb) and wild-type allele (15.8-kb). Clones with the correct homologous recombination were expanded to confirm their integrity and karyotyped to verify their euploid karyotype. Positive clones tested negative for mycoplasma (Venor GeM) were used for C57/BL6 blastocyst injection to generate chimeric mice. Chimeric males were bred to C57BL/6 wild-type females for germline transmission. Offspring were screened by PCR for neo-deletion using primers F1: 5’-AAGAACTGTCGCTCCCTGATGAAC-3’ and R1 5’-CCTGGGGGGAAAGAATGAATTGCT-3’ (Fig. 1a). Expression of Protamine-1 Cre transgene in the male germ line resulted in the deletion of the floxed neomycin gene in all the first pups born, leaving behind a single loxP site after the avi tag cassette. The Cre recombinase transgene, derived from the PC3 embryonic stem cell background, was bred out in the process of backcrossing to C57BL/6 mice. Biotin-tag XAB2 knock-in mice were further crossed to transgenic BirA transgenic mice ^27^. Mice were kept on a regular diet and housed at the IMBB animal house, which operates in compliance with the ‘Animal Welfare Act’ of the Greek government, using the ‘Guide for the Care and Use of Laboratory Animals’ as its standard. As required by Greek law, formal permission to generate and use genetically modified animals was obtained from the responsible local and national authorities. The independent Animal Ethical Committee at FORTH approved all animal studies.

### Mass Spectrometry studies

Proteins eluted from the beads were separated by SDS/PAGE electrophoresis on an 10% polyacrylamide gel and stained with Colloidal blue silver (ThermoFisher Scientific, USA; ^74^. SDS-PAGE gel lanes were cut into 2-mm slices and subjected to in-gel reduction with dithiothreitol, alkylation with iodoacetamide and digested with trypsin (sequencing grade; Promega), as described previously ^75,76^. Nanoflow liquid chromatography tandem mass spectrometry (nLC-MS/MS) was performed on an EASY-nLC coupled to an Orbitrap Fusion Tribid mass spectrometer (Thermo) operating in positive mode. Peptides were separated on a ReproSil-C18 reversed-phase column (Dr. Maisch; 15 cm × 50μ m) using a linear gradient of 0–80% acetonitrile (in 0.1% formic acid) during 90 min at a rate of 200 nl/min. The elution was directly sprayed into the electrospray ionization (ESI) source of the mass spectrometer. Spectra were acquired in continuum mode; fragmentation of the peptides was performed in data-dependent mode by HCD. Raw mass spectrometry data were analyzed with the MaxQuant software suite ^77^ (version 1.6.0.16) as described previously ^76^ with the additional options ‘LFQ’ and ‘iBAQ’ selected. The A false discovery rate of 0.01 for proteins and peptides and a minimum peptide length of seven amino acids were set. The Andromeda search engine was used to search the MS/MS spectra against the Uniprot database (taxonomy: *Mus musculus*, release September 2017), concatenated with the reversed versions of all sequences. A maximum of two missed cleavages was allowed. The peptide tolerance was set to 10 ppm and the fragment ion tolerance was set to 0.6Da for HCD spectra. The enzyme specificity was set to trypsin and cysteine carbamidomethylation was set as a fixed modification. Both the PSM and protein FDR were set to 0.01. In case the identified peptides of two proteins were the same or the identified peptides of one protein included all peptides of another protein, these proteins were combined by MaxQuant and reported as one protein group. The generated ‘proteingroups.txt’ table was filtered for contaminants and reverse hits. For interactor identification, t-test-based statistics was applied on LFQ. First, the logarithm (log 2) of the LFQ values were taken, resulting in a Gaussian distribution of the data. This allowed imputation of missing values by normal distribution (width=0.3, shift=1.8), assuming these proteins were close to the detection limit. Statistical outliers for the XAB2 samples compared to BirA control were then determined using two-tailed t-test. Multiple testing correction was applied by using a permutation-based false discovery rate (FDR) method in Perseus.

### Cells, colony formation and unscheduled DNA synthesis assays

Mouse embryonic stem cells JM8A3N.1 were cultured on gelatinized tissue culture dishes in medium containing Dulbecco’s modified Eagle’s medium (DMEM) supplemented with 15% fetal bovine serum (FBS), 50μg mL^-1^ streptomycin, 50 U mL^-1^ penicillin (Gibco), 2mM L-glutamine (Gibco), 1% non-essential amino acids (Gibco), 0.1mM β-Mercaptoethanol (Applichem), 1μM MEK inhibitor PD0325901 (Selleck), 3μM GSK inhibitor CHIR99021 (Selleck). Primary MEFs (P4) and HEPA cells were cultured in standard medium containing Dulbecco’s modified Eagle’s medium (DMEM) supplemented with 10% fetal bovine serum (FBS), 50μg mL^-1^ streptomycin, 50 U mL^-1^ penicillin (Gibco), 2mM L-glutamine (Gibco). Serum starved MEFs were grown in DMEM supplemented with 1% FBS for 24hrs before treatments. To knockdown *Xab2* an oligo RNA was designed at the position 1685 (si1) and at position 1460 (si2) of the cDNA (Invitrogen). Mouse embryonic stem cells were transfected in suspension with Lipofectamine 2000 (Invitrogen) and subsequently plated in a 60mm plate. HEPA cells were transfected with Polyplus Jet prime according to the manufacturer’s protocol. As a non-targeting control, AllStars negative (Qiagen) was used. MEFs were transfected with Amaxa Mouse Embryonic Fibroblast Nucleofector Kit. Both siRNAs were used at a final concentration of 50nM. MEFs were protein transfected with recombinant RNase H (NEB) using Project Reagent Transfection Kit (Thermo, 89850). 5units of RNase H were used per 24well plate. Cells were transfected 1hr before tRA/tRA illudin treatments. Cells were rinsed with PBS, exposed to UVC irradiation at the indicated doses, MMC (10 μg mL^-1^) (AppliChem), tRA (10 μM) (Sigma-Aldrich), illudin S (50ngr/ml), isoginketin (30-6μΜ) and cultured at 37°C for 4 to 12h prior to subsequent experiments. Pre-incubation with isoginketin (30-60μM), ATM inhibitor (10μM) and ATR inhibitor (10μM), started 6hrs (isoginketin) or 1 h (ATM, ATR inhibitors) before UVC irradiation and lasted throughout the experiment. For cell survival experiments, a total of 200-300 HEPA cells 24h post knock-down or primary MEFs or were seeded in 10 cm Petri dishes. The next day, MEFs were exposed to MMC treatment for 4 h or to UVC irradiation and incubated for 10 days. Colonies were stained with Coomassie blue (0.2% Coomassie blue, 50% methanol, 7% acetic acid), and the number of colonies was counted and expressed as a percentage of the treated cells relative to that of the untreated control. Three dishes per dose were used and at least three independent survival experiments were performed. For viability experiments, Trypan blue inclusion was used at selected time points post transfection. Culture medium was collected and centrifuged to precipitate dead cells. HEPA cells were detached from the tissue culture dish by trypsin-0.5% EDTA, resuspended in culture medium and merged with the fraction of cells from the medium. The cell suspension was subsequently diluted 1:5 with 0.4% Trypan blue and the number of viable and dead cells was counted in a Neubauer haemocytometer under the microscope. The number of viable cells was expressed as the percentage of viable cells relative to that of the control cells. At least three independent viability experiments per time point were performed. DNA repair synthesis was determined by 5-ethynyl-2’-deoxyuridine (EdU) incorporation. Primary MEFs grown on coverslips were globally UVC irradiated and incubated for 2.5 h in medium supplemented with 10 mM EdU. After EdU incorporation, cells were washed with PBS followed by fixation with 2% formaldehyde in PBS. Coverslips were blocked for 30 min with 10% FBS in PBS, followed by 1 h incubation with 10mM sodium ascorbate and 4mM CuSO4 containing Alexa Fluor594 azide (ThermoFischer Scientific A10270) and DAPI staining. The number of EdU-positive cells among at least 200 cells was counted, and the percentage of EdU-positive cells relative to the total number of cells was calculated. DNA transcription sites were labelled as follows. MEFs were grown on coverslips. After treatments cells were washed with ice-cold TBS buffer (10mMTris-HCl, 150mMNaCl, 5mMMgCl_2_) and further washed with glycerol buffer (20mMTris-HCl, 25%glycerol, 5mMMgCl_2_, 0,5mMEGTA) for 10min on ice. Washed cells were permeabilised with 0,5% TritonX-100 in glycerol buffer (with 25U/ml RNase inhibitor) on ice for 3min and immediately incubated at RT for 30min with nucleic acid synthesis buffer (50mMTris-HCl pH7.4, 10mM MgCl_2_, 150mM NaCl, 25%glycerol, 25U/ml RNase inhibitor, protease inhibitors, supplemented with 0,5mM ATP, CTP, GTP and 0,2mMBrUTP. After incorporation, cells were fixed with 4% formaldehyde in PBS on ice for 10min. Immunofluorescence with a-BrdU antibody was performed as described below.

### Immunofluorescence, Antibodies, Westerns blots and FACS

Immunofluorescence experiments were performed as previously described ^27,58^. Briefly, cells were fixed in 4% formaldehyde, permeabilized with 0.5% Triton-X and blocked with 1% BSA. After one-hour incubation with primary antibodies, secondary fluorescent antibodies were added and DAPI was used for nuclear counterstaining. Samples were imaged with SP8 confocal microscope (Leica). For local DNA damage infliction, cells were UV-irradiated (60 J m^-2^) through isopore polycarbonate membranes containing 3-μm-diameter pores (Millipore). For CPD immunodetection, nuclear DNA was denatured with 1M HCl for 30min. For S9.6 immunofluorescence cells were fixed with ice-cold methanol at -20^°^C for 10min.RNase H treatment was performed post-fixation at 37^°^C in PBS supplemented with 10-15 Units RNaseH for 30min. For whole cell extract preparations, cell pellets were resuspended in 150mM NaCl, 50mM Tris pH=7.5, 5% Glycerol, 1% NP-40, 1mM MgCl) and incubated on ice for 30 min. For cell cycle analysis cells were fixed with 70% ethanol for 30min, washed with PBS, RNase A treated (1mgr/ml) at 37^°^C for 30min and stained with propidium iodide (RT,20mgr/ml) for 1hrs.Antibodies against XAB2 (wb: 1:2000, IF: 1:1000), PRP19 (wb: 1:1000, IF: 1:500), β-TUB (wb: 1:2000) were from Abcam. BCAS2 (wb: 1:5000, IF: 1:1000), DDB1 (IF: 1:500) were from Novus. HA (Y-11, western blotting (wb): 1:500), ERCC1 (D-10, wb: 1:500), RAD9A (wb: 1:300), POLII (wb: 1:500, IF: 1:50), XPA (wb: 1:500), XPF (F-11, wb: 1:500) and XPG (sc12558, wb: 1:200) were from Santa Cruz Biotechnology. γH2Ax (05-636, IF: 1:12.000) and S9.6 (MABE1095, IF: 1:300) was from Millipore. DDB1 (wb: 1:5000), XPC (wb: 1:1000) and CSB (wb: 1:1000) were from Bethyl Laboratories. Streptavidin-HRP (wb: 1:12,000) was from Upstate Biotechnology. FLAGM2 (F3165, wb 1:2.000) was from Sigma-Aldrich. CPD (IF: 1:50) was Cosmo Bio Ltd (TDM2). BrdU (1:300, IF) was from BD Pharmingen.

### Co-immunoprecipitation assays

Nuclear protein extracts from 15-day-old livers or cells were prepared as previously described ^27^ using the high-salt extraction method (10mM HEPES-KOH pH 7.9, 380mM KCl, 3mM MgCl2, 0.2mM EDTA, 20% glycerol and protease inhibitors). For immunoprecipitation (IP) assays, nuclear lysates were diluted threefold by adding ice-cold HENG buffer (10mM HEPES-KOH pH 7.9, 1.5mM MgCl2, 0.25mM EDTA, 20% glycerol) and precipitated with antibodies overnight at 4°C followed by incubation for 2 h with protein G Sepharose beads (Millipore). Normal mouse or rabbit IgG (Santa Cruz) was used as a negative control. Immunoprecipitates were washed five times (10mM HEPES-KOH pH7.9, 300mM KCl, 0.3% NP40, 1.5mM MgCl2, 0.25mM EDTA, 20% glycerol and protease inhibitors), eluted and resolved on 10% SDS-PAGE. Pulldowns were performed with 0.6-0.7mg of nuclear extracts using M-280 paramagnetic streptavidin beads (Invitrogen) as previously described23.

### Differential alternative splicing analysis

Global quality of FASTQ files with raw RNA-seq reads was analyzed using *fastqc* v.0.11.5 (https://www.bioinformatics.babraham.ac.uk/projects/fastqc/). *Vast-tools* ^78,79^ aligning and read processing software was used for quantification of alternative sequence inclusion levels from FASTQ files using VASTD-DB annotation ^80^ for mouse genome assembly mm9. To ensure sufficient read coverage for alternative splicing quantification, we considered only *vast-tools*’ events with a minimum mapability-corrected read coverage score of “VLOW” across all samples. The relative inclusion of an alternative sequence (exon or intron), henceforth called alternative splicing event, is based on the number of junction reads supporting inclusion (#inc) and exclusion (#exc), used to quantify percent spliced-in (PSI) ^81^ values. The beta distribution (conjugate prior probability distribution for the binomial), constrained in the]0,1[interval and characterized by a mean value of α/(α + β) from the distribution’s shape parameters α and β, was exploited in modelling the precision of each PSI from its supporting coverage (#inc and #exc). We used R function *rbeta* to emit, for each sample, 500 values from a beta distribution with α = #inc + 1 and β = #exc + 1 where 1 is added to ensure that both the shape parameters, α and β, are different from zero. Since beta distributions get narrower with increasing shape parameters (for the same PSI value), this parametrization allows the scattering of emitted values to serve as a surrogate for that PSI’s dispersion (given the original read coverage supporting it), while the distribution’s mean value, (#inc + 1) / (#inc + 1 + #exc + 1), is an approximation of the empirical PSI. For each event, *rbeta*-emitted values were grouped per condition (Control and Xab2 siRNA) and the median of all emitted values per group was used to determine global PSIs per condition and the difference between these, ΔPSI = PSI_Xab2 siRNA_ - PSI_Control_. For each alternative splicing event, the beta distribution’s emitted values were used in calculating the significance of its ΔPSI. First, the difference between randomly ordered Xab2 siRNA and Control vectors of emitted values was calculated. The significance of each ΔPSI was set as the ratio between the number of differences that are greater than zero and the total number of differences, reflecting the probability of PSI_Xab2 siRNA_ being greater than PSI_Control_. Differentially spliced events were considered as those with a probability of a |ΔPSI| > 0 greater than 0.8 and an absolute ΔPSI greater than 5% (Table EV3). To assess the enrichment of differentially spliced events in positive or negative ΔPSI values within exon skipping and intron retention events, we tested the null hypothesis that the proportion of positive ΔPSI values (i.e. PSI_Xab2 siRNA_ > PSI_Control_) was equal to 0.5 using R function *prop.test*.

### RNA immunoprecipitation studies

RNA immunoprecipitation in HEPA cells and MEFs was performed as previously described ^82^ with a few modifications. In brief, cells (4x150mm plate) were harvested by trypsinization and the cell pellet was resuspended in NP-40 lysis buffer for 10min on ice. Nuclei were washed once in NP-40 Lysis buffer and subsequently resuspended in 1 ml RIP buffer (150mM KCl, 25mM Tris pH 7.4, 5mM EDTA, 0.5mM DTT, 0.5% NP40, 1mM PMSF, 40U/mL RNaseOut; Invitrogen). Resuspended nuclei were mechanically sheared using a syringe (26G) with 5-7 strokes. Nuclear membranes and debris were pelleted by centrifugation at 13,000 RPM for 10 min. Antibodies (5µg) were added to supernatant and incubated overnight at 4°C with gentle rotation. Fifty microliters of protein G Sepharose beads (Millipore) were added and incubated for 2hr at 4°C with gentle rotation. Beads were pelleted at 6.000 RPM for 3 min, the supernatant was removed, and beads were washed three times in 1 mL wash buffer (280mM KCl, 25mM Tris pH 7.4, 5mM EDTA, 0.5mM DTT, 0.5% NP40, 1mM PMSF, 40 U/mL RNaseOut; Invitrogen) for 10min at 4oC, followed by one wash in PBS. Beads were resuspended in 1 ml of Trizol and co-precipitated RNAs were isolated according to the manufacturer’s protocol. RNA was precipitated with Ethanol/Sodium acetate in the presence of Glycoblue at -20°C overnight. Isolated RNA was treated with DNase I (Promega) followed by reverse transcription with random primers (Invitrogen). For RNA immunoprecipitation in liver tissue, livers from two P15 mice were minced and subsequently cross-linked with 0.1% formaldehyde for 10 min at room temperature. After addition of 0.25 M glycine for 5 min, cells were harvested and lysed with RIPA buffer (50mM Tris-HCl [pH 7.4], 1% NP-40, 0.5% Na deoxycholate 0.05% SDS, 1mM EDTA, and 150mM NaCl) followed by sonication at 4°C. Nuclear membrane and debris were pelleted by centrifugation at 13,000 RPM for 10 min. Antibodies (5μ were added to supernatant and incubated overnight at 4°C with gentle rotation. Fifty microliters of protein G Sepharose beads (Millipore) were added and incubated for 2 hours at 4°C with gentle rotation. Beads were pelleted at 6.000 RPM for 3 min, the supernatant was removed, and beads were washed three times in 1 mL wash buffer (50mM Tris-HCl [pH 7.4], 1% NP-40, 0.5% Na deoxycholate, 0.05% SDS, 1mM EDTA, and 350mM NaCl) for 10min at 4°C, followed by one wash with PBS. Crosslinks were reversed by adding 100μ elution buffer (50mM Tris-HCl pH 6.5, 5mM EDTA, 1% SDS, and 10mM DTT) and heating for 45 min at 70°C. RNA was purified with TRIzol reagent, treated with DNase I, and used for first-strand cDNA synthesis.

### RNA-Seq and Quantitative PCR studies

Total RNA was isolated from cells using a Total RNA isolation kit (Qiagen) as described by the manufacturer. For RNA-Seq studies, libraries were prepared using the Illumina® TruSeq® mRNA stranded sample preparation Kit for HEPA cells and for mESCs. Library preparation started with 1µ g total RNA. After poly-A selection (using poly-T oligo-attached magnetic beads), mRNA was purified and fragmented using divalent cations under elevated temperature. The RNA fragments underwent reverse transcription using random primers. This is followed by second strand cDNA synthesis with DNA polymerase I and RNase H. After end repair and A-tailing, indexing adapters were ligated. The products were then purified and amplified to create the final cDNA libraries. After library validation and quantification (Agilent 2100 Bioanalyzer), equimolar amounts of all 12 libraries were pooled. The pool was quantified by using the Peqlab KAPA Library Quantification Kit and the Applied Biosystems 7900HT Sequence Detection System. The pool was sequenced by using an Illumina HiSeq 4000 sequencer with a paired-end (2x 75 cycles) protocol. Quantitative PCR (Q-PCR) was performed with a Biorad 1000-series thermal cycler according to the instructions of the manufacturer (Biorad) as previously described ^27^. All relevant data and primer sequences for the genes tested by qPCR are available upon request.

### DIP

DRIP analysis was based on CHIP analysis with some modifications. DIP analysis was performed without a cross-linking step. Nuclei were isolated using 0,5% NP-40 buffer. Isolated nuclei were resuspended in TE buffer supplemented with 0,5%SDS and 100mgr proteinase K. Genomic DNA was isolated after addition of KoAc (1M) and isopropanol precipitation. DNA was sonicated on ice 3 min using Covaris S220 Focused ultrasonicator. Samples were treated with RNase H (10units/5µg DNA) at 37°C overnight. Samples were immunoprecipitated with S9.6 antibodies (8 μ antibody/ 5μ DNA) overnight at 4°C followed by incubation for 3 hours with protein G-Sepharose beads (Millipore) and washed sequentially. The complexes were eluted and purified DNA fragments were analyzed by qPCR using sets of primers targeting different regions of related genes.

### DRIP western analysis

DRIP western analysis was performed as described previously ^73^. Non-crosslinked cells were lysed in 0.5% NP40 buffer for 10 min on ice. Pelleted nuclei were lysed in RSB buffer (10 mM Tris-HCl pH 7.5, 200 mM NaCl, 2.5 mM MgCl2) with 0.2% sodium deoxycholate [NaDOC, 0.1% SDS and 0.5% Triton X-100, and extracts were sonicated for 10 min (Diagenode Bioruptor). Extracts were then diluted 1:4 in RSB with 0.5% Triton X-100 (RSB + T) and subjected to IP with the S9.6 antibody (8 μg DNA), bound to protein A dynabeads (Invitrogen), and pre-blocked with 1mg/ml BSA/PBS for 1 hr. IgG2a antibodies were used as control. RNase H (PureLink, Invitrogen) was added before IP as in DRIP. Beads were washed 4x with RSB + T; 2x with RSB; and eluted in 1x Laemmli.

### Sequential native ChIP analysis

Non-cross-linked cells were lysed using 0,5% NP-40 buffer. Chromatin was digested with MNase (50 Units/0.5mgDNA) at 37^0^C for 10min. S1 chromatin fraction was isolated. S2 chromatin fraction was dialyzed against Tris-EDTA buffer for 2hrs and isolated. S1 and S2 chromatin fractions were used for pulldown using M280 paramagnetic streptavidin beads (Invitrogen). After sequential washes complexes were eluted. A fraction of them was kept for qPCR analysis and the rest was used for S9.6 immunoprecipitation as described above.

### Primer sequences

Primer: ATM (intron) Forward: TGGCTGCATCTACCTGTGAC Reverse: TGCTAGTCAGCCCTCCTCAT, Primer: ATM (exon) Forward: ACCCAGGTCTATGTGCACAA Reverse: CCCCTGTTCAAAAGCCACTC, Primer: ERCC2 (exon) Forward: GGCTGTGATCATGTTTGGAG, Reverse: CAGCAAAGACCATGAGTCCA, Primer: ERCC2 (intron) Reverse: CCCTGACAGCACTGTTTCC, Primer: U4 snRNA Forward: GCAGTGGCAGTATCGTAGCC Reverse: AAAATTGCCAATGCCGACTA, Primer: U6 snRNA Forward: CGCTTCGGCAGCACATATAC Reverse: ATGGAACGCTTCACGAATTT, Primer: Rad9 (exon) Forward: GTAGTAGCTGCTGGGACTCA Reverse: TTCGGGATAGCGAATGGACA, Primer Rad9 (intron) Forward: GGCAACGTGAAGGGTAAGTT Reverse: -CGGGGAGGAGAACAGAAAGT, Primer Ercc1 (exon) Forward: AAAGATCCCCAGCAGGCTC Reverse: ATAAGGAGGTCGGCTGGCTT, Primer Ercc1 (intron) Forward: CTAAGCCAAGCATGGTGACA Reverse: TGGGGAGAACAGAACAAACC, Primer CDK7 (exon) Forward: ACTGCAGCACATCTTCATCG Reverse: GGGGCGGTTACTGAAGTACT, Primer DDB1 (exon) Forward: CCTCGAATCCATCCTGATGAC, Reverse: AGAGCAAGCAAAGACGTTGG, Primer DDB1 (intron) Forward: CCTCGAATCCATCCTGATGAC, Reverse: AGCTGCTTGGTTAAGGCTCA, Primer H2Afz (exon) Forward: TAAAGCGTATCACCCCTCG Reverse: TCAGCGATTTGTGGATGTGT, Primer H2Afz (intron) Forward: CGTATCACCCCTCGTCACTT Reverse: ATGACATACCACCACCAGCA, Primer Rarb2 (P) Forward: GGGAGTTTTTAAGCGCTGTG Reverse: ACCACTTCTGTCACACGGAAT, Primer Rarb2 (C): Forward: ATCTCTTGAAAAAGGTGCCGAACGT Reverse: GGAAATGTCTCACTGCAGCAGTGGT, Stra6 (P) Forward: AGGCACCCTTTTAAGGAGGA Reverse: TTCCACACCTCACAAAGACG, Stra6 (C) Forward: ACCACACATACCAAAACTTCCTG Reverse: CGGGGTAAAGACGTACCTTC, ChordC (P) Forward: GCAGTCCGGTAGGAAATCTG Reverse: CCGGTACTGCTTCAGGAATTT, ChordC (C) Forward: TTCAAGCCCCTAAGCCAGTA, Reverse: TACACGAGTGGACACTGCAA

### Quantification and Statistical analysis

A two-tailed t-test was used to extract the statistically significant data by means of the IBM SPSS Statistics 19 (IBM) and the *R* software for statistical computing (www.r-project.org). Significant over-representation of pathways and gene networks was determined by Gene Ontology (http://geneontology.org/). Data analysis is discussed also in the Method Details section. Experiments were repeated at least 3 times. The data exhibited normal distribution (where applicable). There was no estimation of group variation before experiments. Error bars indicate standard deviation unless stated otherwise (standard error of the mean; s.e.m.). For animal studies, each biological replicate consists of 3-5 mouse tissues or cell cultures per genotype per time point or treatment. No statistical method was used to predetermine sample size. None of the samples or animals was excluded from the experiment. The animals or the experiments were non-randomized. The investigators were not blinded to allocation during experiments and outcome assessment.

### Data availability

The mass spectrometry proteomics data have been deposited to the ProteomeXchange Consortium (http://proteomecentral.proteomexchange.org) via the PRIDE partner repository with the dataset identifier PXD014084. The RNA-Seq data are deposited in ArrayExpress (https://www.ebi.ac.uk/arrayexpress/), (E-MTAB-8035). All other data and reagents are available from the authors upon reasonable request.

## Acknowledgements

The Horizon 2020 ERC Consolidator grant “DeFiNER” (GA64663), the Horizon 2020 Marie Curie ITNs “Chromatin3D” (GA622934), “aDDRess” (GA812829) and “HealthAge” “(GA812830), the Hellenic Foundation for Research and Innovation (HFRI) and the General Secretariat for Research and Technology (GSRT) under grant agreement HFRI-1059 and HFRI-FM17-631 and the “Fondation Santé” grant supported this work. GC is supported by the IKY postdoctoral research fellowship program (MIS: 5001552), co-financed by the European Social Fund-ESF and the Greek government. NLB-M is supported by an EMBO Installation Grant (3057) and an Investigador FCT (Fundação para a Ciência e a Tecnologia) Starting Grant (IF/00595/2014); MA-F is supported by an FCT PhD fellowship (PD/BD/128283/2017) and Fundação AstraZeneca.

## Author Contributions

EG, MT, NB, MA-F, EL, KS, GC, PT, TK and NLB-M performed the experiments and/or analyzed data. GG interpreted data and wrote the manuscript. All relevant data are available from the authors.

## References

1. Jurica, M. S. & Moore, M. J. Pre-mRNA splicing: awash in a sea of proteins. Mol Cell 12, 5–14 (2003).

2. Hoskins, A. A. & Moore, M. J. The spliceosome: a flexible, reversible macromolecular machine. Trends Biochem Sci 37, 179–188, doi:10.1016/j.tibs.2012.02.009 (2012).

3. Bentley, D. L. Rules of engagement: co-transcriptional recruitment of pre-mRNA processing factors. Curr Opin Cell Biol 17, 251–256, doi:10.1016/j.ceb.2005.04.006 (2005).

4. Kornblihtt, A. R., de la Mata, M., Fededa, J. P., Munoz, M. J. & Nogues, G. Multiple links between transcription and splicing. RNA 10, 1489–1498, doi:10.1261/rna.7100104 (2004).

5. de la Mata, M. & Kornblihtt, A. R. RNA polymerase II C-terminal domain mediates regulation of alternative splicing by SRp20. Nat Struct Mol Biol 13, 973–980, doi:10.1038/nsmb1155 (2006).

6. Alexander, R. & Beggs, J. D. Cross-talk in transcription, splicing and chromatin: who makes the first call? Biochem Soc Trans 38, 1251–1256, doi:10.1042/BST0381251 (2010).

7. Zhou, H. L., Luo, G., Wise, J. A. & Lou, H. Regulation of alternative splicing by local histone modifications: potential roles for RNA-guided mechanisms. Nucleic Acids Res 42, 701–713, doi:10.1093/nar/gkt875 (2014).

8. de la Mata, M. et al. A slow RNA polymerase II affects alternative splicing in vivo. Mol Cell 12, 525–532 (2003).

9. Munoz, M. J. et al. DNA damage regulates alternative splicing through inhibition of RNA polymerase II elongation. Cell 137, 708–720, doi:10.1016/j.cell.2009.03.010 (2009).

10. Paronetto, M. P., Minana, B. & Valcarcel, J. The Ewing sarcoma protein regulates DNA damage-induced alternative splicing. Mol Cell 43, 353–368, doi:10.1016/j.molcel.2011.05.035 (2011).

11. Tresini, M. et al. The core spliceosome as target and effector of non-canonical ATM signalling. Nature 523, 53–58, doi:10.1038/nature14512 (2015).

12. Lenzken, S. C., Loffreda, A. & Barabino, S. M. RNA splicing: a new player in the DNA damage response. Int J Cell Biol 2013, 153634, doi:10.1155/2013/153634 (2013).

13. Montecucco, A. & Biamonti, G. Pre-mRNA processing factors meet the DNA damage response. Front Genet 4, 102, doi:10.3389/fgene.2013.00102 (2013).

14. Dutertre, M. et al. Cotranscriptional exon skipping in the genotoxic stress response. Nat Struct Mol Biol 17, 1358–1366, doi:10.1038/nsmb.1912 (2010).

15. Zeytuni, N. & Zarivach, R. Structural and functional discussion of the tetra-trico-peptide repeat, a protein interaction module. Structure 20, 397–405, doi:10.1016/j.str.2012.01.006 (2012).

16. Ben-Yehuda, S. et al. Genetic and physical interactions between factors involved in both cell cycle progression and pre-mRNA splicing in Saccharomyces cerevisiae. Genetics 156, 1503–1517 (2000).

17. Yonemasu, R. et al. Disruption of mouse XAB2 gene involved in pre-mRNA splicing, transcription and transcription-coupled DNA repair results in preimplantation lethality. DNA Repair (Amst) 4, 479–491, doi:10.1016/j.dnarep.2004.12.004 (2005).

18. Nakatsu, Y. et al. XAB2, a novel tetratricopeptide repeat protein involved in transcription-coupled DNA repair and transcription. J Biol Chem 275, 34931–34937, doi:10.1074/jbc.M004936200 (2000).

19. Chanarat, S., Seizl, M. & Strasser, K. The Prp19 complex is a novel transcription elongation factor required for TREX occupancy at transcribed genes. Genes Dev 25, 1147–1158, doi:10.1101/gad.623411 (2011).

20. Kuraoka, I. et al. Isolation of XAB2 complex involved in pre-mRNA splicing, transcription, and transcription-coupled repair. J Biol Chem 283, 940–950, doi:10.1074/jbc.M706647200 (2008).

21. Marechal, A. et al. PRP19 transforms into a sensor of RPA-ssDNA after DNA damage and drives ATR activation via a ubiquitin-mediated circuitry. Mol Cell 53, 235–246, doi:10.1016/j.molcel.2013.11.002 (2014).

22. Song, E. J. et al. The Prp19 complex and the Usp4Sart3 deubiquitinating enzyme control reversible ubiquitination at the spliceosome. Genes Dev 24, 1434–1447, doi:10.1101/gad.1925010 (2010).

23. Hirose, T. et al. A spliceosomal intron binding protein, IBP160, links position-dependent assembly of intron-encoded box C/D snoRNP to pre-mRNA splicing. Mol Cell 23, 673–684, doi:10.1016/j.molcel.2006.07.011 (2006).

24. Onyango, D. O., Howard, S. M., Neherin, K., Yanez, D. A. & Stark, J. M. Tetratricopeptide repeat factor XAB2 mediates the end resection step of homologous recombination. Nucleic Acids Res 44, 5702–5716, doi:10.1093/nar/gkw275 (2016).

25. Hou, S. et al. XAB2 depletion induces intron retention in POLR2A to impair global transcription and promote cellular senescence. Nucleic Acids Res 47, 8239–8254, doi:10.1093/nar/gkz532 (2019).

26. O’Gorman, S., Dagenais, N. A., Qian, M. & Marchuk, Y. Protamine-Cre recombinase transgenes efficiently recombine target sequences in the male germ line of mice, but not in embryonic stem cells. Proc Natl Acad Sci U S A 94, 14602–14607 (1997).

27. Chatzinikolaou, G. et al. ERCC1-XPF cooperates with CTCF and cohesin to facilitate the developmental silencing of imprinted genes. Nat Cell Biol 19, 421–432, doi:10.1038/ncb3499 (2017).

28. Chan, S. P., Kao, D. I., Tsai, W. Y. & Cheng, S. C. The Prp19p-associated complex in spliceosome activation. Science 302, 279–282, doi:10.1126/science.1086602 (2003).

29. Hou, S. et al. XAB2 functions in mitotic cell cycle progression via transcriptional regulation of CENPE. Cell Death Dis 7, e2409, doi:10.1038/cddis.2016.313 (2016).

30. Bondar, T. et al. Cul4A and DDB1 associate with Skp2 to target p27Kip1 for proteolysis involving the COP9 signalosome. Mol Cell Biol 26, 2531–2539, doi:10.1128/MCB.26.7.2531-2539.2006 (2006).

31. Scharer, O. D. Nucleotide excision repair in eukaryotes. Cold Spring Harb Perspect Biol 5, a012609, doi:10.1101/cshperspect.a012609 (2013).

32. Kelly, C. M. & Latimer, J. J. Unscheduled DNA synthesis: a functional assay for global genomic nucleotide excision repair. Methods Mol Biol 291, 303–320 (2005).

33. Grote, M. et al. Molecular architecture of the human Prp19/CDC5L complex. Mol Cell Biol 30, 2105–2119, doi:10.1128/MCB.01505-09 (2010).

34. Lareau, L. F., Brooks, A. N., Soergel, D. A., Meng, Q. & Brenner, S. E. The coupling of alternative splicing and nonsense-mediated mRNA decay. Adv Exp Med Biol 623, 190–211 (2007).

35. de Lima Morais, D. A. & Harrison, P. M. Large-scale evidence for conservation of NMD candidature across mammals. PLoS One 5, e11695, doi:10.1371/journal.pone.0011695 (2010).

36. Boutz, P. L., Bhutkar, A. & Sharp, P. A. Detained introns are a novel, widespread class of post-transcriptionally spliced introns. Genes Dev 29, 63–80, doi:10.1101/gad.247361.114 (2015).

37. Ding, J., Miao, Z. H., Meng, L. H. & Geng, M. Y. Emerging cancer therapeutic opportunities target DNA-repair systems. Trends in pharmacological sciences 27, 338–344, doi:10.1016/j.tips.2006.04.007 (2006).

38. Garinis, G. A. et al. Transcriptome analysis reveals cyclobutane pyrimidine dimers as a major source of UV-induced DNA breaks. The EMBO journal 24, 3952–3962, doi:10.1038/sj.emboj.7600849 (2005).

39. Garinis, G. A., Jans, J. & van der Horst, G. T. Photolyases: capturing the light to battle skin cancer. Future Oncol 2, 191–199, doi:10.2217/14796694.2.2.191 (2006).

40. Jaspers, N. G. et al. Anti-tumour compounds illudin S and Irofulven induce DNA lesions ignored by global repair and exclusively processed by transcription- and replication-coupled repair pathways. DNA Repair (Amst) 1, 1027–1038, doi:10.1016/s1568-7864(02)00166-0 (2002).

41. van der Horst, G. T. et al. Defective transcription-coupled repair in Cockayne syndrome B mice is associated with skin cancer predisposition. Cell 89, 425–435, doi:10.1016/s0092-8674(00)80223-8 (1997).

42. Roberts, R. W. & Crothers, D. M. Stability and properties of double and triple helices: dramatic effects of RNA or DNA backbone composition. Science 258, 1463–1466, doi:10.1126/science.1279808 (1992).

43. Skourti-Stathaki, K., Proudfoot, N. J. & Gromak, N. Human senataxin resolves RNA/DNA hybrids formed at transcriptional pause sites to promote Xrn2-dependent termination. Mol Cell 42, 794–805, doi:10.1016/j.molcel.2011.04.026 (2011).

44. Li, X. & Manley, J. L. Inactivation of the SR protein splicing factor ASF/SF2 results in genomic instability. Cell 122, 365–378, doi:10.1016/j.cell.2005.06.008 (2005).

45. Huertas, P. & Aguilera, A. Cotranscriptionally formed DNA:RNA hybrids mediate transcription elongation impairment and transcription-associated recombination. Mol Cell 12, 711–721, doi:10.1016/j.molcel.2003.08.010 (2003).

46. Wahba, L., Costantino, L., Tan, F. J., Zimmer, A. & Koshland, D. S1-DRIP-seq identifies high expression and polyA tracts as major contributors to R-loop formation. Genes Dev 30, 1327-1338, doi:10.1101/gad.280834.116 (2016).

47. Stork, C. T. et al. Co-transcriptional R-loops are the main cause of estrogen-induced DNA damage. Elife 5, doi:10.7554/eLife.17548 (2016).

48. Bastien, J. & Rochette-Egly, C. Nuclear retinoid receptors and the transcription of retinoid-target genes. Gene 328, 1–16, doi:10.1016/j.gene.2003.12.005 (2004).

49. Brustel, J., Kozik, Z., Gromak, N., Savic, V. & Sweet, S. M. M. Large XPF-dependent deletions following misrepair of a DNA double strand break are prevented by the RNA:DNA helicase Senataxin. Sci Rep 8, 3850, doi:10.1038/s41598-018-21806-y (2018).

50. Sollier, J. et al. Transcription-coupled nucleotide excision repair factors promote R-loop-induced genome instability. Mol Cell 56, 777–785, doi:10.1016/j.molcel.2014.10.020 (2014).

51. Cristini, A. et al. Dual Processing of R-Loops and Topoisomerase I Induces Transcription-Dependent DNA Double-Strand Breaks. Cell Rep 28, 3167–3181 e3166, doi:10.1016/j.celrep.2019.08.041 (2019).

52. Hamperl, S., Bocek, M. J., Saldivar, J. C., Swigut, T. & Cimprich, K. A. Transcription-Replication Conflict Orientation Modulates R-Loop Levels and Activates Distinct DNA Damage Responses. Cell 170, 774–786 e719, doi:10.1016/j.cell.2017.07.043 (2017).

53. Schumacher, B., Garinis, G. A. & Hoeijmakers, J. H. Age to survive: DNA damage and aging. Trends in genetics : TIG 24, 77–85, doi:10.1016/j.tig.2007.11.004 (2008).

54. Garinis, G. A., van der Horst, G. T., Vijg, J. & Hoeijmakers, J. H. DNA damage and ageing: new-age ideas for an age-old problem. Nat Cell Biol 10, 1241–1247, doi:10.1038/ncb1108-1241 (2008).

55. Lopez-Mejia, I. C. et al. A conserved splicing mechanism of the LMNA gene controls premature aging. Hum Mol Genet 20, 4540–4555, doi:10.1093/hmg/ddr385 (2011).

56. Sen, S., Jumaa, H. & Webster, N. J. Splicing factor SRSF3 is crucial for hepatocyte differentiation and metabolic function. Nat Commun 4, 1336, doi:10.1038/ncomms2342 (2013).

57. Wong, C. M., Xu, L. & Yau, M. Y. Alternative mRNA Splicing in the Pathogenesis of Obesity. Int J Mol Sci 19, doi:10.3390/ijms19020632 (2018).

58. Karakasilioti, I. et al. DNA damage triggers a chronic autoinflammatory response, leading to fat depletion in NER progeria. Cell Metab 18, 403–415, doi:10.1016/j.cmet.2013.08.011 (2013).

59. Hutton, M. et al. Association of missense and 5’-splice-site mutations in tau with the inherited dementia FTDP-17. Nature 393, 702–705, doi:10.1038/31508 (1998).

60. Cartegni, L. & Krainer, A. R. Disruption of an SF2/ASF-dependent exonic splicing enhancer in SMN2 causes spinal muscular atrophy in the absence of SMN1. Nat Genet 30, 377–384, doi:10.1038/ng854 (2002).

61. Mordes, D. et al. Pre-mRNA splicing and retinitis pigmentosa. Mol Vis 12, 1259–1271 (2006).

62. Kamileri, I., Karakasilioti, I. & Garinis, G. A. Nucleotide excision repair: new tricks with old bricks. Trends in genetics : TIG 28, 566–573, doi:10.1016/j.tig.2012.06.004 (2012).

63. Quesada, V. et al. Exome sequencing identifies recurrent mutations of the splicing factor SF3B1 gene in chronic lymphocytic leukemia. Nat Genet 44, 47–52, doi:10.1038/ng.1032 (2011).

64. Su, H. et al. SHQ1 regulation of RNA splicing is required for T-lymphoblastic leukemia cell survival. Nat Commun 9, 4281, doi:10.1038/s41467-018-06523-4 (2018).

65. Fei, D. L. et al. Impaired hematopoiesis and leukemia development in mice with a conditional knock-in allele of a mutant splicing factor gene U2af1. Proc Natl Acad Sci U S A 115, E10437–E10446, doi:10.1073/pnas.1812669115 (2018).

66. Black, K. L. et al. Aberrant splicing in B-cell acute lymphoblastic leukemia. Nucleic Acids Res, doi:10.1093/nar/gky1231 (2018).

67. Hoeijmakers, J. H. DNA damage, aging, and cancer. N Engl J Med 361, 1475–1485, doi:10.1056/NEJMra0804615 (2009).

68. Apostolou, Z., Chatzinikolaou, G., Stratigi, K. & Garinis, G. A. Nucleotide Excision Repair and Transcription-Associated Genome Instability. Bioessays 41, e1800201, doi:10.1002/bies.201800201 (2019).

69. Guil, S., Long, J. C. & Caceres, J. F. hnRNP A1 relocalization to the stress granules reflects a role in the stress response. Mol Cell Biol 26, 5744–5758, doi:10.1128/MCB.00224-06 (2006).

70. Llorian, M. et al. Nucleocytoplasmic shuttling of the splicing factor SIPP1. J Biol Chem 280, 38862–38869, doi:10.1074/jbc.M509185200 (2005).

71. Busa, R., Geremia, R. & Sette, C. Genotoxic stress causes the accumulation of the splicing regulator Sam68 in nuclear foci of transcriptionally active chromatin. Nucleic Acids Res 38, 3005–3018, doi:10.1093/nar/gkq004 (2010).

72. Shkreta, L. & Chabot, B. The RNA Splicing Response to DNA Damage. Biomolecules 5, 2935–2977, doi:10.3390/biom5042935 (2015).

73. Cristini, A., Groh, M., Kristiansen, M. S. & Gromak, N. RNA/DNA Hybrid Interactome Identifies DXH9 as a Molecular Player in Transcriptional Termination and R-Loop-Associated DNA Damage. Cell Rep 23, 1891–1905, doi:10.1016/j.celrep.2018.04.025 (2018).

74. Candiano, G. et al. Blue silver: a very sensitive colloidal Coomassie G-250 staining for proteome analysis. Electrophoresis 25, 1327–1333, doi:10.1002/elps.200305844 (2004).

75. Wilm, M. et al. Femtomole sequencing of proteins from polyacrylamide gels by nano-electrospray mass spectrometry. Nature 379, 466–469, doi:Doi 10.1038/379466a0 (1996).

76. Schwertman, P. et al. UV-sensitive syndrome protein UVSSA recruits USP7 to regulate transcription-coupled repair. Nat Genet 44, 598–602, doi:10.1038/ng.2230 (2012).

77. Cox, J. et al. A practical guide to the MaxQuant computational platform for SILAC-based quantitative proteomics. Nat Protoc 4, 698–705, doi:10.1038/nprot.2009.36 (2009).

78. Irimia, M. et al. A highly conserved program of neuronal microexons is misregulated in autistic brains. Cell 159, 1511–1523, doi:10.1016/j.cell.2014.11.035 (2014).

79. Braunschweig, U. et al. Widespread intron retention in mammals functionally tunes transcriptomes. Genome Res 24, 1774–1786, doi:10.1101/gr.177790.114 (2014).

80. Tapial, J. et al. An atlas of alternative splicing profiles and functional associations reveals new regulatory programs and genes that simultaneously express multiple major isoforms. Genome Res 27, 1759–1768, doi:10.1101/gr.220962.117 (2017).

81. Barbosa-Morais, N. L. et al. The evolutionary landscape of alternative splicing in vertebrate species. Science 338, 1587–1593, doi:10.1126/science.1230612 (2012).

82. Rinn, J. L. et al. Functional demarcation of active and silent chromatin domains in human HOX loci by noncoding RNAs. Cell 129, 1311–1323, doi:10.1016/j.cell.2007.05.022 (2007).

