## Supplemental information for "The Splicing Factor XAB2 interacts with ERCC1-XPF and XPG for RNA-loop processing during mammalian development"

**Supplemental figure legends**

**Figure S1.** (**A**). List of significantly over-represented biological processes of 636 shared XAB2-bound proteins. Data are shown as in Figure 2E. (**B**). bXAB2 pulldowns followed by western blot analysis with DDB1 or XPA in UV-irradiated (10J/m2) and control primary MEFs. (**C**). Immunofluorescence detection of XAB2 with UV-induced CPDs; white arrowheads indicate the immunodetection of XAB2 and CPDs, respectively (D). Immunofluorescence detection of XAB2 with γH2Ax in UV-irradiated HEPA cells (10J/m2). (**E**) Western blot analysis of XAB2 protein levels in si*Xab2* cells and si Scramble (Scrb) cells at 48 hours (h) post-transfection. (F). % of live si*Xab2* cells (Trypan blue inclusion) at the indicated time points post transfection; the red dotted line indicates the 100% survival. (**G**). Number (no) of colonies of si*Xab2* cells at seven days post-transfection (**H).** Western blot analysis of RNAPII in siScrb and si*Xab2* cells at 48h post-transfection. (**I**) Western blot analysis of DDB1, XPA and ERCC1 in untreated, UV-irradiated and MMC-treated siScrb and si*Xab2* cells. (**J**).mRNA levels and (**K**) Pre-mRNA levels of transcripts with retained introns in untreated and 30 or 60μΜ isoG treated MEF cells at 12hrs post-inhibitor treatment (as indicated). (**L**). A table depicting the data from all alternative splicing events considered and the differentially spliced events in si*Xab2* and siScrb HEPA cells. The images shown on Figure S1B-I are representative of experiments that were repeated more than three times.

**Figure S2.** (**A**). Distribution of all alternative splicing events considered and (**B**) differentially spliced events in si*Xab2* and si Scramble (Scrb) HEPA cells. (**C**). Volcano plots of all exon skipping events and (**D**). Intron retention events considered in si*Xab2* and si Scramble (Scrb) HEPA cells. Intron retention (IR); Exon Skipping (EX); Alternative 3’ spliced sites (Alt3’); Alternative 5’ spliced sites (Alt5’); Percent spliced-in (PSI). (**E**). A table depicting the data of all alternative splicing events considered and the differentially spliced events in si*Xab2* and si Scramble (Scrb) mESCs. (**F**). Distribution of all alternative splicing events considered and (**G**) differentially spliced events in si*Xab2* and si Scramble (Scrb) mESCs. (**H**). Volcano plots of all exon skipping events and (**I**). Intron retention events considered in si*Xab2* and siScrb mESCs.

**Figure S3** (**A**). Over-representation of biological processes that are significantly affected by alternative splicing in si*Xab2* HEPA cells. Intron retention (IR); Exon Skipping (EX); Alternative 3’ spliced sites (Alt3’); Alternative 5’ spliced sites (Alt5’); Percent spliced-in (PSI); FDR: False Detection Rate; F.E: Fold Enrichment; GC: Gene count. . (**B**). Over-representation of biological processes that are significantly affected by alternative splicing in si*Xab2* mESCs. (**C-D**)**.** bXAB2 pulldowns for the indicated pre-mRNAs in P15 livers (**E**). Pre-mRNA levels of transcripts with retained introns in si2*Xab2* and siScrb HEPA cells at 48h post-transfection. (**F**). Pre-mRNA levels of UV-irradiated cells compared to untreated controls. (**G**) bXAB2 RNA pull downs on pre-mRNAs in P15 bXAB2 or bXAB2;*Csb^m/m^* mouse livers (n ≥ 3 per genotype). (**H**) RNA:DNA hybrid immunofluorescence detection in untreated and isoginketin treated B6 MEFs *H*igher magnifications of cells in white box and quantifications are presented in Figure 6B.

**Figure S4.** (**A)** Basal levels of R loops were monitored and quantitated by immunofluorescence with S9.6 antibody in si*Scrb* and si*Xab2* transfected B6 MEFs. Higher magnifications of cells in white box are presented in Figure 6C in the case of si1*Xab2* or in insets in the case of si2*Xab2*. Quantifications of immunofluorescence are shown in figure 6C. (**B**). Representative images and quantification of RNA:DNA hybrid immunofluorescence detection in siScrb and si*Xab2* MEFs without UV-irradiation or 8hrs post-UV irradiation. Graph depicts mean S9.6 fluorescence intensity per nucleus from three representative experiment (≥150 nuclei were analyzed per condition). (**C**). Immunofluorescence detection of γH2AX and 53BP1 (white arrowheads) in siScrb and si*Xab2* B6 MEFs with or without UV-irradiation. The graph represents the number of γH2AX+ 53BP1+ cells from three representative experiment (≥150nuclei were analyzed per condition). (**D**) bXAB2 RNA pull downs on U4 and U6 snRNAs in tRA or illudin-tRA treated birA and bXAB2 MEFs with or without RNase H protein transfection. (E) bXAB2 ChIP signals from BirA and tRA- and illudin S/tRA-treated bXAB2 MEFs on the coding region of tRA-induced *Stra6* gene and the tRA-non induced *Chordc1* gene. (**F**). Western blot of RNA/DNA hybrid IP in nuclear extracts derived from B6 MEFs with or without RNaseH treatment, using indicated antibodies. Isotype-matched IgG antibody were used as controls. (**G**). bXAB2 pulldowns (PD: Pulldown) and western blot with anti-XPF, anti-XPG and anti-PRP19 in nuclear extracts derived from bXAB2 and BirA MEFs with or without RNaseH treatment. (**H**). Cell cycle profiling and representative images of FACS analysis of wt. cells cultured in the presence (FBS+) or absence (FBS-) of fetal bovine serum. (**I**) Representative images and quantification of RNA:DNA hybrid immunofluorescence detection of wt. cells cultured in the presence (FBS+) or absence (FBS-) of fetal bovine serum transfected (RNH+) or not (RNH-) with RNaseH1 protein.

**Figure S5**. (A). Western blot of RNA/DNA hybrid IP in nuclear extracts derived from wt. MEFs with or without RNaseH treatment, using the indicated antibodies. Isotype-matched IgG antibody were used as controls. (**B**).Western blot of RNA:DNA hybrid IP in nuclear extracts derived from *Xpa*^-/-^ MEFs. Isotype-matched IgG antibody were used as controls. (C). Representative images and quantification of RNA:DNA hybrid immunofluorescence detection of *Xpa*^-/-^ cells. (D). S9.6 DRIP analysis on the promoters of highly transcribed genes i.e. *Igf1* and *Ghr* with or without RNaseH1 (RNH) in wt. and *Csb*^m/m^ P15 livers. (E). bXAB2 ChIP signals from BirA, bXAB2 and bXAB2;*Csb*^m/m^ P15 livers on promoter regions of highly transcribed genes (*Igf1* and *Ghr*).

**Supplemental Table legends**

**Table S1. A list of 1167 bXAB2‐bound proteins identified in the P15 bXAB2 biological replicates compared to BirA control livers.** UP: Unique peptides.

**Table S2. A list of the 636 bXAB2‐bound proteins identified in P15 livers shared across the three bXAB2 biological replicates compared to BirA control livers.** UP: Unique peptides.

**Table S3. A list of 255 differentially expressed genes in HEPA cells transfected with dsRNA targeting the Xab2 transcript vs. scramble control cells.** FC: Fold change.

**Table S4. A list of 333 differentially expressed genes in mESCs transfected with dsRNA targeting the *Xab2* transcript vs. scramble control cells.** FC: Fold change.

**Table S5. Xab2 -induced differential splicing events in HEPA cells.** COORD: Coordinates.

**Table S6. Xab2 -induced differential splicing events in mESCs.** COORD: Coordinates.

 *. P ≤ 0.05. **. P ≤ 0.01. ***. P ≤ 0.001.
