## Supplemental table 2 for "The Splicing Factor XAB2 interacts with ERCC1-XPF and XPG for RNA-loop processing during mammalian development"

**Table S2. A list of the 636 XAB2-bound proteins identified in P15 livers shared across the three bXAB2 biological replicates compared to BirA control livers.**

| Access. no | Symbol | bXab UP1 | bXab_UP3 | bXab_UP2 | BirA UP1 | BirA UP3 | BirA UP2 |
|---|---|---|---|---|---|---|---|
| Q9QXS1 | Actn4 | 188 | 197 | 202 | 44 | 62 | 193 |
| Q5YD48 | A1cf | 8 | 8 | 16 | 1 | 0,01 | 6 |
| O35452 | Tnxb | 116 | 130 | 137 | 30 | 58 | 124 |
| Q68FG2 | Sptbn2 | 103 | 111 | 118 | 31 | 49 | 101 |
| E9PZ16 | Hspg2 | 97 | 115 | 115 | 27 | 44 | 105 |
| E9QP46 | Syne2 | 97 | 187 | 235 | 6 | 38 | 154 |
| F6XCT0 | Macf1 | 40 | 48 | 44 | 0,01 | 0,01 | 0,01 |
| Q921L6 | Cttn | 23 | 24 | 27 | 0,01 | 0,01 | 0,01 |
| A0A0R4J0V1 | Ints10 | 8 | 10 | 11 | 0,01 | 0,01 | 0,01 |
| E9Q360 | Ints10 | 8 | 10 | 11 | 0,01 | 0,01 | 0,01 |
| A2A9P6 | Cdk11b | 7 | 10 | 9 | 0,01 | 0,01 | 0,01 |
| Q9JKX4 | Aatf | 6 | 10 | 14 | 0,01 | 0,01 | 7 |
| E9PWQ3 | Col6a3 | 60 | 67 | 66 | 34 | 26 | 52 |
| A0A087WR50 | Fn1 | 59 | 72 | 74 | 23 | 32 | 64 |
| Q3UHL6 | Fn1 | 59 | 72 | 74 | 23 | 32 | 64 |
| B7ZNH7 | Col14a1 | 57 | 63 | 71 | 33 | 46 | 62 |
| A0A1D5RLS1 | Eps15l1 | 4 | 8 | 8 | 0,01 | 0,01 | 0,01 |
| D3Z315 | Cope | 6 | 7 | 6 | 0,01 | 0,01 | 0,01 |
| E9PVU0 | Myo6 | 49 | 50 | 57 | 11 | 20 | 49 |
| F6RJ39 | Acin1 | 25 | 33 | 43 | 6 | 20 | 33 |
| Q5SYD0 | Myo1d | 47 | 55 | 55 | 16 | 26 | 51 |
| Q80YX1 | Tnc | 44 | 51 | 54 | 16 | 15 | 48 |
| Q8R2S9 | Actr8 | 20 | 24 | 23 | 2 | 10 | 19 |
| Q9QYC0 | Add1 | 14 | 21 | 23 | 3 | 5 | 17 |
| Q9Z103 | Adnp | 8 | 16 | 21 | 0,01 | 0,01 | 13 |
| Q9QZQ1 | Afdn | 40 | 53 | 59 | 5 | 9 | 0,01 |
| Q9QZQ1 | Afdn | 40 | 53 | 59 | 5 | 9 | 46 |
| Q8CJF7 | Ahctf1 | 32 | 59 | 61 | 5 | 15 | 48 |
| P53995 | Anapc1 | 19 | 31 | 41 | 0,01 | 8 | 27 |
| Q91W96 | Anapc4 | 14 | 13 | 18 | 0,01 | 2 | 14 |
| O35488 | Slc27a2 | 34 | 32 | 35 | 28 | 22 | 31 |
| Q8BTZ4 | Anapc5 | 9 | 13 | 17 | 0,01 | 4 | 10 |
| Q8BTZ4 | Anapc5 | 9 | 13 | 17 | 0,01 | 4 | 10 |
| Q9WVM3 | Anapc7 | 17 | 17 | 23 | 0,01 | 6 | 16 |
| A0A0R4J1N7 | Ank1 | 11 | 20 | 21 | 0,01 | 0,01 | 9 |
| Q8C8R3 | Ank2 | 11 | 25 | 28 | 0,01 | 0,01 | 14 |
| P26231 | Ctnna1 | 30 | 41 | 42 | 19 | 19 | 36 |
| P97449 | Anpep | 5 | 10 | 15 | 0,01 | 0,01 | 10 |
| Q8CFQ3 | Aqr | 32 | 40 | 39 | 1 | 14 | 29 |
| A2AQA7 | Aqr | 32 | 40 | 39 | 1 | 14 | 29 |
| E9Q7E2 | Arid2 | 13 | 20 | 17 | 0,01 | 0,01 | 9 |
| Q9WTL8 | Arntl | 7 | 7 | 12 | 0,01 | 0,01 | 5 |
| Q9WTL8 | Arntl | 7 | 7 | 12 | 0,01 | 0,01 | 5 |
| Q8BI84 | Mia3 | 26 | 54 | 40 | 0,01 | 4 | 19 |

| | | | | | | | |
|---|---|---|---|---|---|---|---|
| P98203 | Arvcf | 19 | 23 | 23 | 0,01 | 3 | 17 |
| P98203 | Arvcf | 19 | 23 | 23 | 0,01 | 3 | 17 |
| P98203 | Arvcf | 19 | 23 | 23 | 0,01 | 3 | 17 |
| P98203 | Arvcf | 19 | 23 | 23 | 0,01 | 3 | 17 |
| Q80YQ1 | Thbs1 | 25 | 34 | 35 | 6 | 11 | 31 |
| A0A1L1STC6 | Syne1 | 25 | 70 | 77 | 0,01 | 2 | 40 |
| Q9EP89 | Lactb | 25 | 24 | 26 | 10 | 12 | 22 |
| Q3U741 | Ddx17 | 24 | 28 | 30 | 13 | 17 | 25 |
| E9PZJ8 | Ascc3 | 13 | 18 | 14 | 0,01 | 0,01 | 10 |
| Q925I1 | Atad3 | 32 | 32 | 29 | 10 | 12 | 24 |
| M0QWP1 | Agrn | 22 | 40 | 43 | 3 | 3 | 39 |
| P56480 | Atp5b | 19 | 19 | 21 | 14 | 11 | 16 |
| A2AS05 | Helz2 | 21 | 12 | 11 | 0,01 | 0,01 | 3 |
| A2AS03 | Helz2 | 21 | 12 | 11 | 0,01 | 0,01 | 3 |
| G3X973 | Stab1 | 21 | 21 | 24 | 0,01 | 0,01 | 18 |
| A2AKD7 | Snta1 | 5 | 8 | 8 | 0,01 | 0,01 | 2 |
| Q03265 | Atp5a1 | 20 | 20 | 21 | 15 | 12 | 15 |
| Q8R4H2 | Arhgef12 | 11 | 15 | 14 | 0,01 | 0,01 | 4 |
| F8VQN6 | Arhgef12 | 11 | 15 | 14 | 0,01 | 0,01 | 4 |
| P08032 | Spta1 | 17 | 13 | 13 | 0,01 | 0,01 | 5 |
| Q8K019 | Bclaf1 | 9 | 25 | 27 | 0,01 | 5 | 19 |
| Q8K019 | Bclaf1 | 9 | 25 | 27 | 0,01 | 5 | 19 |
| Q3TL44 | Nlrx1 | 9 | 12 | 12 | 1 | 0,01 | 3 |
| Q8CCJ3 | Ufl1 | 8 | 11 | 9 | 0,01 | 0,01 | 3 |
| Q99JY9 | Actr3 | 19 | 19 | 21 | 10 | 10 | 17 |
| Q6PAL0 | Bend3 | 25 | 29 | 34 | 0,01 | 8 | 29 |
| P35441 | Thbs1 | 24 | 33 | 35 | 6 | 10 | 0,01 |
| Q6PGF5 | Bms1 | 17 | 19 | 23 | 0,01 | 2 | 19 |
| Q921S7 | Mrpl37 | 8 | 11 | 8 | 0,01 | 0,01 | 4 |
| E9Q6A7 | Bptf | 19 | 36 | 39 | 0,01 | 3 | 23 |
| E9QQ10 | Akap9 | 7 | 12 | 10 | 0,01 | 0,01 | 4 |
| Q8C5W0 | Clmn | 10 | 12 | 9 | 0,01 | 0,01 | 5 |
| Q8C5W0 | Clmn | 10 | 12 | 9 | 0,01 | 0,01 | 5 |
| Q8C5W0 | Clmn | 10 | 12 | 9 | 0,01 | 0,01 | 5 |
| E9Q397 | Sptb | 27 | 24 | 20 | 0,01 | 0,01 | 11 |
| O35646 | Capn6 | 11 | 12 | 19 | 3 | 5 | 13 |
| Q9CVB6 | Arpc2 | 16 | 21 | 19 | 10 | 8 | 15 |
| E9Q2E4 | Hectd4 | 16 | 19 | 18 | 0,01 | 0,01 | 9 |
| Q8CH18 | Ccar1 | 14 | 22 | 32 | 0,01 | 4 | 14 |
| Q8VDP4 | Ccar2 | 15 | 18 | 18 | 0,01 | 2 | 12 |
| Q5SV80 | Myo19 | 6 | 9 | 8 | 0,01 | 0,01 | 4 |
| P82198 | Tgfbi | 15 | 18 | 18 | 9 | 7 | 13 |
| A0A1D5RLQ9 | Cdc42bpa | 12 | 21 | 18 | 0,01 | 0,01 | 8 |
| D3YYN8 | Cdc42bpa | 12 | 21 | 18 | 0,01 | 0,01 | 8 |
| A2A6Q5 | Cdc27 | 9 | 14 | 17 | 1 | 4 | 9 |
| H7BX44 | Cdc42bpa | 12 | 21 | 18 | 0,01 | 0,01 | 8 |
| E9PVY0 | Cdc42bpa | 12 | 21 | 18 | 0,01 | 0,01 | 8 |

| | | | | | | | |
|---|---|---|---|---|---|---|---|
| P61161 | Actr2 | 15 | 15 | 18 | 6 | 9 | 14 |
| Q3UU96 | Cdc42bpa | 12 | 21 | 18 | 0,01 | 0,01 | 8 |
| Q80TN4 | Dnajc16 | 7 | 13 | 19 | 0,01 | 0,01 | 7 |
| J3JS23 | Zfp62 | 14 | 23 | 23 | 0,01 | 6 | 19 |
| A2A6Q5 | Cdc27 | 9 | 14 | 17 | 1 | 4 | 9 |
| Q9DC48 | Cdc40 | 23 | 26 | 26 | 0,01 | 6 | 21 |
| S4R1B9 | Taf1 | 14 | 30 | 30 | 0,01 | 6 | 17 |
| Q6A068 | Cdc5l | 26 | 31 | 37 | 5 | 12 | 27 |
| B1Q2W7 | Taf1 | 14 | 30 | 30 | 0,01 | 6 | 17 |
| P24788 | Cdk11b | 7 | 10 | 9 | 0,01 | 0,01 | 0,01 |
| E9Q3L2 | Pi4ka | 6 | 8 | 9 | 0,01 | 0,01 | 4 |
| A0A140T8I9 | Pi4ka | 6 | 8 | 9 | 0,01 | 0,01 | 4 |
| A2AWT5 | Ubtf | 14 | 24 | 28 | 4 | 8 | 20 |
| A2ADB1 | Spen | 13 | 26 | 35 | 0,01 | 2 | 26 |
| P68368 | Tuba4a | 13 | 13 | 14 | 0,01 | 0,01 | 10 |
| O35218 | Cpsf2 | 13 | 22 | 29 | 1 | 5 | 19 |
| A0A0A0MQA! | Tuba4a | 13 | 13 | 14 | 0,01 | 0,01 | 10 |
| Q9CRB9 | Chchd3 | 13 | 16 | 15 | 8 | 6 | 11 |
| P40201 | Chd1 | 24 | 36 | 39 | 0,01 | 8 | 29 |
| Q920A7 | Afg3l1 | 6 | 11 | 14 | 0,01 | 0,01 | 5 |
| Q5SUH7 | Clint1 | 9 | 14 | 11 | 0,01 | 0,01 | 7 |
| Q6PDQ2 | Chd4 | 40 | 56 | 60 | 4 | 16 | 50 |
| E9QAS5 | Chd4 | 40 | 56 | 60 | 4 | 16 | 50 |
| E9QAS4 | Chd4 | 40 | 56 | 60 | 4 | 16 | 50 |
| A2AJK6 | Chd7 | 13 | 29 | 33 | 0,01 | 0,01 | 21 |
| F8WHU7 | Stag1 | 12 | 23 | 26 | 0,01 | 4 | 9 |
| Q8CGZ0 | Cherp | 14 | 18 | 20 | 0,01 | 4 | 14 |
| A0A1D5RL92 | Cherp | 14 | 18 | 20 | 0,01 | 4 | 14 |
| Q8C147 | Dock8 | 12 | 23 | 22 | 0,01 | 0,01 | 11 |
| Q5SUH6 | Clint1 | 9 | 14 | 11 | 0,01 | 0,01 | 7 |
| B1AYU7 | Cwc22 | 16 | 17 | 21 | 0,01 | 0,01 | 12 |
| E9QMQ3 | Triobp | 8 | 16 | 14 | 0,01 | 0,01 | 9 |
| G5E8J2 | Ank1 | 11 | 20 | 21 | 0,01 | 0,01 | 9 |
| G8JL84 | Ank1 | 11 | 20 | 21 | 0,01 | 0,01 | 9 |
| Q02357 | Ank1 | 11 | 20 | 21 | 0,01 | 0,01 | 9 |
| Q8R1B4 | Eif3c | 9 | 17 | 13 | 1 | 0,01 | 7 |
| A0A1W2P711 | Mia2 | 11 | 25 | 16 | 0,01 | 0,01 | 6 |
| B7ZW98 | Ank1 | 11 | 20 | 21 | 0,01 | 0,01 | 9 |
| E9QNT8 | Ank1 | 11 | 20 | 21 | 0,01 | 0,01 | 9 |
| Q91ZV0 | Mia2 | 11 | 25 | 16 | 0,01 | 0,01 | 6 |
| D3YTV8 | Ank1 | 11 | 20 | 21 | 0,01 | 0,01 | 9 |
| Q0VGY9 | Ank1 | 11 | 20 | 21 | 0,01 | 0,01 | 9 |
| Q02357 | Ank1 | 11 | 20 | 21 | 0,01 | 0,01 | 9 |
| Q02357 | Ank1 | 11 | 20 | 21 | 0,01 | 0,01 | 9 |
| Q8C196 | Cps1 | 37 | 32 | 28 | 18 | 10 | 18 |
| Q9R0Q6 | Arpc1a | 11 | 9 | 13 | 6 | 0,01 | 8 |
| Q9EPU4 | Cpsf1 | 27 | 39 | 49 | 0,01 | 11 | 34 |

| | | | | | | | |
|---|---|---|---|---|---|---|---|
| Q810A7 | Ddx42 | 6 | 9 | 10 | 0,01 | 0,01 | 6 |
| Q569Z5 | Ddx46 | 5 | 11 | 12 | 0,01 | 0,01 | 5 |
| E0CYH7 | Filip1l | 9 | 20 | 20 | 0,01 | 1 | 16 |
| Q5F2E8 | Taok1 | 12 | 11 | 11 | 0,01 | 0,01 | 7 |
| P31001 | Des | 32 | 32 | 36 | 19 | 17 | 30 |
| O35286 | Dhx15 | 28 | 33 | 36 | 8 | 20 | 29 |
| O55111 | Dsg2 | 8 | 15 | 13 | 3 | 0,01 | 8 |
| A2ACQ1 | Dhx35 | 10 | 15 | 17 | 0,01 | 2 | 10 |
| Q8C8Z9 | Dtnb | 10 | 13 | 13 | 0,01 | 2 | 7 |
| O70585 | Dtnb | 10 | 13 | 13 | 0,01 | 2 | 7 |
| A2A4P0 | Dhx8 | 19 | 20 | 29 | 0,01 | 4 | 16 |
| A0A087WPL5 | Dhx9 | 50 | 53 | 62 | 26 | 0,01 | 0,01 |
| Q8C6K9 | Col6a6 | 8 | 11 | 8 | 2 | 0,01 | 4 |
| Q91X77 | Cyp2c50 | 8 | 8 | 4 | 0,01 | 0,01 | 0,01 |
| Q811D0 | Dlg1 | 11 | 16 | 13 | 2 | 2 | 8 |
| Q811D0 | Dlg1 | 11 | 15 | 13 | 2 | 2 | 8 |
| Q3UNN4 | Smarcc1 | 8 | 17 | 16 | 3 | 0,01 | 10 |
| P11531 | Dmd | 13 | 23 | 23 | 0,01 | 4 | 13 |
| P63037 | Dnaja1 | 10 | 9 | 14 | 1 | 2 | 7 |
| H3BLH0 | Smarca2 | 29 | 34 | 35 | 0,01 | 0,01 | 25 |
| E9Q6A6 | Col6a6 | 8 | 11 | 8 | 2 | 0,01 | 4 |
| Q7TT50 | Cdc42bpb | 40 | 48 | 42 | 3 | 5 | 30 |
| A0A0G2JDM7 | Anapc5 | 9 | 13 | 17 | 0,01 | 4 | 10 |
| A0A0G2JE03 | Anapc5 | 9 | 13 | 17 | 0,01 | 4 | 10 |
| J3KMH5 | Mia3 | 8 | 22 | 17 | 0,01 | 0,01 | 0,01 |
| Q3TWF7 | Anapc5 | 9 | 13 | 17 | 0,01 | 4 | 10 |
| B2LVG5 | Cpsf4 | 7 | 10 | 11 | 0,01 | 1 | 7 |
| G5E8K2 | Ank3 | 7 | 19 | 23 | 0,01 | 3 | 13 |
| G5E8K5 | Ank3 | 7 | 19 | 23 | 0,01 | 3 | 13 |
| A0A0R4J2B6 | Rbbp5 | 7 | 13 | 12 | 1 | 2 | 5 |
| G5E8K5 | Ank3 | 7 | 19 | 23 | 0,01 | 3 | 13 |
| K3W4P2 | Ints1 | 8 | 12 | 19 | 0,01 | 0,01 | 12 |
| A0A0G2JH17 | Ints1 | 8 | 12 | 19 | 0,01 | 0,01 | 12 |
| G5E8K5 | Ank3 | 7 | 19 | 23 | 0,01 | 3 | 13 |
| O08810 | Eftud2 | 33 | 44 | 45 | 18 | 23 | 36 |
| P23116 | Eif3a | 19 | 29 | 24 | 0,01 | 4 | 14 |
| D3Z7I0 | Gtf3c2 | 14 | 21 | 21 | 0,01 | 2 | 15 |
| P60229 | Eif3e | 6 | 11 | 10 | 0,01 | 0,01 | 5 |
| Q8QZY1 | Eif3l | 8 | 9 | 11 | 0,01 | 0,01 | 5 |
| G5E8K3 | Ank3 | 7 | 19 | 23 | 0,01 | 3 | 13 |
| Q8BYC6 | Taok3 | 11 | 12 | 13 | 2 | 0,01 | 9 |
| Q8CHI8 | Ep400 | 4 | 10 | 13 | 0,01 | 0,01 | 8 |
| Q8CHI8 | Ep400 | 4 | 10 | 13 | 0,01 | 0,01 | 8 |
| Q8CHI8 | Ep400 | 4 | 10 | 13 | 0,01 | 0,01 | 8 |
| Q8CHI8 | Ep400 | 4 | 10 | 13 | 0,01 | 0,01 | 8 |
| Q66JV4 | Rbm12b2 | 20 | 23 | 32 | 0,01 | 0,01 | 23 |
| O70318 | Epb41l2 | 22 | 34 | 31 | 0,01 | 3 | 23 |

| | | | | | | | |
|---|---|---|---|---|---|---|---|
| Q8R0W0 | Eppk1 | 16 | 22 | 18 | 0,01 | 4 | 14 |
| Q60902 | Eps15l1 | 4 | 8 | 8 | 0,01 | 0,01 | 0,01 |
| Q60902 | Eps15l1 | 4 | 8 | 8 | 0,01 | 0,01 | 0,01 |
| Q60902 | Eps15l1 | 4 | 8 | 8 | 0,01 | 0,01 | 0,01 |
| Q60902 | Eps15l1 | 4 | 8 | 8 | 0,01 | 0,01 | 0,01 |
| A0A1W2P812 | Ank3 | 7 | 19 | 23 | 0,01 | 3 | 13 |
| Q3V1V3 | Esf1 | 9 | 14 | 14 | 0,01 | 4 | 10 |
| Q0KL02 | Trio | 19 | 21 | 28 | 0,01 | 3 | 17 |
| Q0KL02 | Trio | 19 | 21 | 28 | 0,01 | 3 | 17 |
| S4R2K9 | Ank3 | 7 | 19 | 23 | 0,01 | 3 | 13 |
| P56960 | Exosc10 | 18 | 20 | 22 | 1 | 3 | 18 |
| Q8VBX6 | Mpdz | 6 | 26 | 27 | 0,01 | 0,01 | 15 |
| Q9Z1T2 | Thbs4 | 12 | 12 | 15 | 0,01 | 2 | 11 |
| Q69ZR9 | Fam208a | 27 | 44 | 54 | 0,01 | 10 | 44 |
| P48725 | Pcnt | 6 | 12 | 11 | 0,01 | 0,01 | 5 |
| A0A1L1SQR4 | Dock6 | 30 | 40 | 48 | 0,01 | 0,01 | 35 |
| Q7TPD1 | Fbxo11 | 6 | 9 | 7 | 0,01 | 3 | 0,01 |
| Q8CGB6 | Tns2 | 17 | 27 | 37 | 0,01 | 5 | 24 |
| Q8CGB6 | Tns2 | 17 | 27 | 37 | 0,01 | 5 | 24 |
| Q8CGB6 | Tns2 | 17 | 27 | 37 | 0,01 | 5 | 24 |
| Q8CGB6 | Tns2 | 17 | 27 | 37 | 0,01 | 5 | 24 |
| Q6P6L0 | Filip1l | 9 | 20 | 20 | 0,01 | 1 | 16 |
| Q6P6L0 | Filip1l | 9 | 20 | 20 | 0,01 | 1 | 16 |
| Q8BTM8 | Flna | 83 | 94 | 97 | 13 | 37 | 84 |
| B7FAU9 | Flna | 83 | 94 | 97 | 13 | 37 | 84 |
| Q80X90 | Flnb | 74 | 88 | 82 | 4 | 20 | 73 |
| D3YUX2 | Mpdz | 6 | 26 | 27 | 0,01 | 0,01 | 15 |
| Q80U35 | Arhgef17 | 6 | 13 | 20 | 0,01 | 2 | 14 |
| Q8VBX6 | Mpdz | 6 | 26 | 27 | 0,01 | 0,01 | 15 |
| Q91Z49 | Fyttd1 | 11 | 16 | 20 | 5 | 7 | 14 |
| Q9R1K5 | Fzr1 | 5 | 9 | 11 | 0,01 | 0,01 | 6 |
| F8VPV0 | Pcnt | 6 | 12 | 11 | 0,01 | 0,01 | 5 |
| Q4VAA7 | Snx33 | 6 | 11 | 10 | 0,01 | 0,01 | 5 |
| P26039 | Tln1 | 36 | 50 | 51 | 3 | 5 | 40 |
| A0A1W2P737 | Pcnt | 6 | 12 | 11 | 0,01 | 0,01 | 5 |
| Q8VBX6 | Mpdz | 6 | 26 | 27 | 0,01 | 0,01 | 15 |
| Q8R322 | Gle1 | 9 | 16 | 22 | 0,01 | 0,01 | 16 |
| D3YYT1 | Glyr1 | 11 | 17 | 21 | 6 | 8 | 15 |
| A2AWL7 | Mga | 5 | 12 | 21 | 0,01 | 2 | 14 |
| Q8CI11 | Gnl3 | 14 | 18 | 17 | 1 | 6 | 12 |
| G5E8T6 | Magi3 | 11 | 12 | 16 | 0,01 | 0,01 | 12 |
| F6SMY7 | Mycbp2 | 52 | 74 | 75 | 0,01 | 5 | 61 |
| A0A087WP63 | Gm42715 | 5 | 12 | 11 | 0,01 | 0,01 | 6 |
| Q80X80 | C2cd2l | 11 | 15 | 15 | 3 | 2 | 9 |
| Q99MI1 | Erc1 | 5 | 16 | 13 | 0,01 | 0,01 | 6 |
| Q6P5H2 | Nes | 24 | 34 | 40 | 0,01 | 0,01 | 34 |
| Q8K284 | Gtf3c1 | 46 | 64 | 65 | 0,01 | 10 | 49 |

| | | | | | | | |
|---|---|---|---|---|---|---|---|
| Q3TMP1 | Gtf3c3 | 20 | 25 | 25 | 6 | 6 | 21 |
| Q8BMQ2 | Gtf3c4 | 26 | 36 | 38 | 3 | 9 | 29 |
| Q8R2T8 | Gtf3c5 | 13 | 17 | 17 | 2 | 7 | 11 |
| Q2PFD7 | Psd3 | 5 | 11 | 10 | 0,01 | 1 | 3 |
| Q8R2T8 | Gtf3c5 | 13 | 17 | 17 | 2 | 7 | 11 |
| V9GXP8 | Erc1 | 5 | 16 | 13 | 0,01 | 0,01 | 6 |
| G3X922 | Dnajc13 | 35 | 51 | 51 | 0,01 | 6 | 41 |
| Q2PFD7 | Psd3 | 5 | 11 | 10 | 0,01 | 1 | 3 |
| A0A0R4J0J8 | Specc1l | 20 | 28 | 33 | 1 | 5 | 24 |
| Q2KN98 | Specc1l | 20 | 28 | 33 | 1 | 5 | 24 |
| A2AWL7 | Mga | 5 | 12 | 21 | 0,01 | 2 | 14 |
| E9QLG3 | Mga | 5 | 12 | 21 | 0,01 | 2 | 14 |
| G3X9B1 | Heatr1 | 39 | 43 | 54 | 5 | 19 | 49 |
| E9QAM5 | Helz2 | 21 | 12 | 11 | 0,01 | 0,01 | 3 |
| Q9R1Y5 | Hic1 | 7 | 14 | 19 | 0,01 | 4 | 12 |
| Z4YJP0 | Hic1 | 7 | 14 | 19 | 0,01 | 4 | 12 |
| F8VPM7 | Erc1 | 5 | 16 | 13 | 0,01 | 0,01 | 6 |
| A0A1D5RM5[illegible] | Tns1 | 26 | 37 | 39 | 0,01 | 3 | 32 |
| A0A087WQS[illegible] | Tns1 | 26 | 37 | 39 | 0,01 | 3 | 32 |
| E9Q0S6 | Tns1 | 26 | 37 | 39 | 0,01 | 3 | 32 |
| V9GXF0 | Erc1 | 5 | 15 | 13 | 0,01 | 0,01 | 0,01 |
| V9GXH3 | Erc1 | 5 | 15 | 13 | 0,01 | 0,01 | 0,01 |
| Q2PFD7 | Psd3 | 5 | 11 | 10 | 0,01 | 1 | 3 |
| Q8BJA3 | Hmbox1 | 9 | 12 | 17 | 0,01 | 7 | 11 |
| H3BKM3 | Hmbox1 | 9 | 12 | 17 | 0,01 | 7 | 11 |
| H3BKF8 | Hmbox1 | 9 | 12 | 17 | 0,01 | 7 | 11 |
| E9PUQ5 | Golga2 | 16 | 25 | 21 | 4 | 4 | 13 |
| E9Q0A3 | Arhgef11 | 11 | 20 | 22 | 0,01 | 2 | 17 |
| Q9Z204 | Hnrnpc | 23 | 26 | 28 | 12 | 14 | 24 |
| Q9Z204 | Hnrnpc | 23 | 26 | 28 | 12 | 14 | 24 |
| P15116 | Cdh2 | 9 | 12 | 12 | 3 | 2 | 8 |
| D3YYT0 | Cdh2 | 9 | 12 | 12 | 3 | 2 | 8 |
| P14873 | Map1b | 4 | 15 | 14 | 0,01 | 0,01 | 9 |
| P70333 | Hnrnph2 | 12 | 12 | 14 | 0,01 | 4 | 9 |
| E9PVP1 | Crybg1 | 15 | 20 | 22 | 0,01 | 3 | 15 |
| A0A0G2JG52 | Crybg1 | 15 | 20 | 22 | 0,01 | 3 | 15 |
| G5E924 | Hnrnpl | 12 | 16 | 22 | 2 | 2 | 13 |
| Q8VHM5 | Hnrnpr | 11 | 10 | 14 | 0,01 | 0,01 | 9 |
| Q00PI9 | Hnrnpul2 | 43 | 42 | 48 | 24 | 22 | 41 |
| Z4YKB8 | Hp1bp3 | 21 | 20 | 21 | 13 | 9 | 17 |
| Q3TEA8 | Hp1bp3 | 21 | 20 | 21 | 13 | 9 | 17 |
| P20029 | Hspa5 | 27 | 33 | 34 | 15 | 15 | 29 |
| Q9Z1M8 | Ik | 13 | 25 | 31 | 4 | 9 | 18 |
| D3YU58 | Anapc7 | 17 | 17 | 23 | 0,01 | 6 | 16 |
| Q9Z1X4 | Ilf3 | 30 | 36 | 40 | 8 | 11 | 33 |
| Q9Z1X4 | Ilf3 | 30 | 36 | 40 | 8 | 11 | 33 |
| G5E8I8 | Cherp | 14 | 18 | 20 | 0,01 | 4 | 14 |

| | | | | | | | |
|---|---|---|---|---|---|---|---|
| Q6ZPV2 | Ino80 | 23 | 31 | 44 | 2 | 11 | 30 |
| Q6ZPV2 | Ino80 | 23 | 31 | 44 | 2 | 11 | 30 |
| Q6P4S8 | Ints1 | 8 | 12 | 19 | 0,01 | 0,01 | 12 |
| Q3UQ44 | Iqgap2 | 42 | 50 | 47 | 5 | 10 | 39 |
| E9Q9B7 | Kidins220 | 4 | 12 | 11 | 0,01 | 1 | 7 |
| S4R1P5 | Dst | 4 | 14 | 19 | 0,01 | 0,01 | 7 |
| Q8BZQ7 | Anapc2 | 11 | 18 | 18 | 0,01 | 3 | 14 |
| Q9JKF1 | Iqgap1 | 20 | 27 | 28 | 0,01 | 4 | 19 |
| P11881 | Itpr1 | 40 | 44 | 47 | 7 | 8 | 30 |
| P11881 | Itpr1 | 40 | 44 | 47 | 7 | 8 | 30 |
| P11881 | Itpr1 | 40 | 44 | 47 | 7 | 8 | 30 |
| P11881 | Itpr1 | 40 | 44 | 47 | 7 | 8 | 30 |
| P11881 | Itpr1 | 40 | 44 | 47 | 7 | 8 | 30 |
| P11881 | Itpr1 | 40 | 44 | 47 | 7 | 8 | 30 |
| P11881 | Itpr1 | 40 | 44 | 47 | 7 | 8 | 30 |
| P11881 | Itpr1 | 40 | 44 | 47 | 7 | 8 | 30 |
| Q9Z329 | Itpr2 | 65 | 67 | 78 | 14 | 23 | 65 |
| P70227 | Itpr3 | 13 | 19 | 17 | 0,01 | 0,01 | 10 |
| Q8C1D8 | Iws1 | 12 | 13 | 16 | 0,01 | 4 | 12 |
| Q6NZF1 | Zc3h11a | 4 | 16 | 21 | 0,01 | 4 | 13 |
| A2AQD5 | Ssfa2 | 17 | 22 | 22 | 2 | 2 | 17 |
| A0A1Y7VMH7 | Kidins220 | 4 | 12 | 11 | 0,01 | 1 | 7 |
| A3KG93 | Kdm1a | 6 | 9 | 12 | 0,01 | 0,01 | 7 |
| Q6ZQ88 | Kdm1a | 6 | 9 | 12 | 0,01 | 0,01 | 7 |
| A0A1Y7VME9 | Kidins220 | 4 | 12 | 11 | 0,01 | 1 | 7 |
| A2AAE1 | Kiaa1109 | 21 | 37 | 37 | 0,01 | 4 | 22 |
| A2AAE1 | Kiaa1109 | 21 | 37 | 37 | 0,01 | 4 | 22 |
| A2AAE1 | Kiaa1109 | 21 | 37 | 37 | 0,01 | 4 | 22 |
| A2AAE1 | Kiaa1109 | 21 | 37 | 37 | 0,01 | 4 | 22 |
| Q6PDK2 | Kmt2d | 9 | 11 | 15 | 0,01 | 0,01 | 8 |
| Q80YR9 | Rbm12b1 | 24 | 28 | 33 | 2 | 6 | 27 |
| Q99L88 | Sntb1 | 19 | 22 | 24 | 4 | 5 | 17 |
| Q68FD5 | Cltc | 39 | 38 | 35 | 8 | 8 | 28 |
| Q5SXR6 | Cltc | 39 | 38 | 35 | 8 | 8 | 28 |
| Q6P5B5 | Fxr2 | 20 | 17 | 23 | 2 | 4 | 19 |
| E9PU96 | Urb1 | 45 | 53 | 72 | 3 | 13 | 55 |
| G5E8G6 | Myo5b | 54 | 54 | 64 | 3 | 10 | 51 |
| P21271 | Myo5b | 54 | 54 | 64 | 3 | 10 | 51 |
| Q5XJE5 | Leo1 | 12 | 16 | 22 | 3 | 7 | 13 |
| Q8BH80 | Vapb | 12 | 12 | 12 | 4 | 3 | 8 |
| Q9QY76 | Vapb | 12 | 12 | 12 | 4 | 3 | 8 |
| P48678 | Lmna | 54 | 49 | 52 | 30 | 35 | 48 |
| Q61586 | Gpam | 21 | 22 | 24 | 6 | 4 | 19 |
| O88322 | Nid2 | 25 | 33 | 35 | 3 | 8 | 28 |
| Q91ZX7 | Lrp1 | 44 | 57 | 55 | 3 | 4 | 36 |
| A0A0R4J0I9 | Lrp1 | 44 | 57 | 55 | 3 | 4 | 36 |
| Q8BWT1 | Acaa2 | 14 | 17 | 15 | 6 | 4 | 11 |

| | | | | | | | |
|---|---|---|---|---|---|---|---|
| Q6RHR9 | Magi1 | 7 | 15 | 15 | 0,01 | 0,01 | 11 |
| Q60847 | Col12a1 | 55 | 50 | 53 | 4 | 12 | 46 |
| E9PX70 | Col12a1 | 55 | 50 | 53 | 4 | 12 | 46 |
| Q6RHR9 | Magi1 | 7 | 15 | 15 | 0,01 | 0,01 | 0,01 |
| Q60847 | Col12a1 | 55 | 50 | 53 | 4 | 12 | 46 |
| Q60847 | Col12a1 | 55 | 50 | 53 | 4 | 12 | 46 |
| A0A0N4SUZ0 | Magi1 | 7 | 15 | 15 | 0,01 | 0,01 | 0,01 |
| Q9EQJ9 | Magi3 | 11 | 12 | 16 | 0,01 | 0,01 | 12 |
| Q8K310 | Matr3 | 48 | 48 | 55 | 16 | 17 | 49 |
| Q91VS8 | Farp2 | 19 | 23 | 23 | 2 | 5 | 19 |
| J3QMC5 | Mdn1 | 7 | 29 | 40 | 0,01 | 0,01 | 16 |
| A2ANY6 | Mdn1 | 7 | 29 | 40 | 0,01 | 0,01 | 16 |
| F7BGR7 | Gm21992 | 11 | 14 | 19 | 0,01 | 3 | 14 |
| C0HKD8 | Mfap1a | 11 | 16 | 18 | 1 | 5 | 12 |
| C0HKD9 | Mfap1b | 11 | 16 | 18 | 1 | 5 | 12 |
| H7BX50 | Mga | 5 | 12 | 21 | 0,01 | 2 | 14 |
| Q8BML1 | Mical2 | 7 | 14 | 18 | 0,01 | 0,01 | 13 |
| D3Z598 | Ltbp4 | 29 | 36 | 40 | 2 | 11 | 35 |
| Q8K4G1 | Ltbp4 | 29 | 36 | 40 | 2 | 11 | 35 |
| A0A087WSN( | Fn1 | 57 | 69 | 71 | 23 | 0,01 | 62 |
| B9EHT6 | Fn1 | 57 | 69 | 71 | 23 | 0,01 | 62 |
| Q810V0 | Mphosph10 | 9 | 7 | 12 | 0,01 | 0,01 | 8 |
| Q3TYA6 | Mphosph8 | 7 | 13 | 17 | 0,01 | 5 | 12 |
| P11247 | Mpo | 6 | 12 | 16 | 0,01 | 2 | 9 |
| Q9JMH9 | Myo18a | 76 | 86 | 86 | 13 | 23 | 71 |
| Q9JMH9 | Myo18a | 76 | 86 | 86 | 13 | 23 | 71 |
| P46978 | Stt3a | 16 | 12 | 15 | 5 | 4 | 8 |
| Q9R190 | Mta2 | 16 | 28 | 27 | 2 | 8 | 21 |
| Q924K8 | Mta3 | 4 | 16 | 13 | 0,01 | 4 | 9 |
| Q3UII8 | Mta3 | 4 | 16 | 13 | 0,01 | 4 | 9 |
| Q80WJ7 | Mtdh | 19 | 21 | 25 | 3 | 8 | 20 |
| A0A0U1RNK7 | Dock7 | 36 | 50 | 51 | 6 | 10 | 44 |
| F8VQB6 | Myo10 | 12 | 13 | 22 | 0,01 | 0,01 | 17 |
| Q8R1A4 | Dock7 | 36 | 50 | 51 | 6 | 10 | 44 |
| A2A9M5 | Dock7 | 36 | 50 | 51 | 6 | 10 | 44 |
| E9PX48 | Dock7 | 36 | 50 | 51 | 6 | 10 | 44 |
| Q8R1A4 | Dock7 | 36 | 50 | 51 | 6 | 10 | 44 |
| A2A9M4 | Dock7 | 36 | 50 | 51 | 6 | 10 | 44 |
| Q5SWZ5 | Mprip | 77 | 80 | 82 | 0,01 | 25 | 78 |
| F8WHL2 | Copa | 31 | 43 | 37 | 7 | 11 | 29 |
| Q8CIE6 | Copa | 31 | 43 | 37 | 7 | 11 | 29 |
| Q8BPM0 | Daam1 | 16 | 16 | 16 | 7 | 3 | 12 |
| Q8BPM0 | Daam1 | 16 | 16 | 16 | 7 | 3 | 12 |
| Q62383 | Supt6h | 46 | 54 | 61 | 4 | 15 | 47 |
| P09405 | Ncl | 17 | 22 | 21 | 4 | 11 | 17 |
| A2AL85 | Asph | 14 | 19 | 18 | 6 | 5 | 14 |
| Q8BSY0 | Asph | 14 | 19 | 18 | 6 | 5 | 14 |

| | | | | | | | |
|---|---|---|---|---|---|---|---|
| Q6PIJ4 | Nfrkb | 14 | 33 | 34 | 5 | 11 | 27 |
| Q8BY02 | Nkrf | 9 | 10 | 18 | 0,01 | 2 | 13 |
| H9KV00 | Son | 39 | 53 | 58 | 2 | 19 | 49 |
| Q6DFW4 | Nop58 | 16 | 22 | 24 | 4 | 10 | 19 |
| Q9D6T0 | Nosip | 4 | 10 | 12 | 0,01 | 0,01 | 5 |
| E9Q7G0 | Numa1 | 65 | 102 | 107 | 31 | 68 | 97 |
| F6ZQA3 | Numa1 | 18 | 32 | 36 | 0,01 | 23 | 32 |
| Q8BH74 | Nup107 | 34 | 36 | 39 | 9 | 14 | 34 |
| Q8R0G9 | Nup133 | 30 | 32 | 38 | 8 | 13 | 29 |
| E9Q3G8 | Nup153 | 13 | 22 | 26 | 3 | 10 | 19 |
| Q9Z0W3 | Nup160 | 34 | 38 | 46 | 8 | 14 | 38 |
| A0A0J9YUD5 | Nup205 | 29 | 26 | 37 | 2 | 5 | 32 |
| B9EJ54 | Nup205 | 29 | 26 | 37 | 2 | 5 | 32 |
| Q9QY81 | Nup210 | 14 | 17 | 21 | 3 | 4 | 15 |
| A0A0R4J1I6 | Nup210 | 14 | 17 | 21 | 3 | 4 | 15 |
| Q9JIH2 | Nup50 | 6 | 16 | 15 | 0,01 | 2 | 8 |
| A0A1B0GSX7 | Nup98 | 38 | 50 | 52 | 8 | 14 | 37 |
| Q8CIC2 | Nupl2 | 9 | 9 | 12 | 0,01 | 0,01 | 8 |
| E9QL43 | Nupl2 | 9 | 9 | 12 | 0,01 | 0,01 | 8 |
| Q61235 | Sntb2 | 22 | 26 | 26 | 4 | 10 | 21 |
| A0A0R4J150 | Osbpl8 | 6 | 11 | 10 | 0,01 | 0,01 | 6 |
| B9EJ86 | Osbpl8 | 6 | 12 | 10 | 0,01 | 0,01 | 6 |
| Q8K2T8 | Paf1 | 18 | 22 | 26 | 3 | 10 | 21 |
| Q63ZW7 | Patj | 28 | 31 | 41 | 2 | 5 | 28 |
| P58501 | Paxbp1 | 19 | 27 | 30 | 0,01 | 3 | 18 |
| P49586 | Pcyt1a | 7 | 9 | 9 | 0,01 | 0,01 | 5 |
| Q6NS46 | Pdcd11 | 28 | 38 | 49 | 0,01 | 11 | 39 |
| Q3TJD7 | Pdlim7 | 9 | 14 | 15 | 0,01 | 3 | 9 |
| Q4VA53 | Pds5b | 14 | 15 | 19 | 0,01 | 4 | 14 |
| Q9QY23 | Pkp3 | 6 | 14 | 14 | 0,01 | 0,01 | 10 |
| Q9QY23 | Pkp3 | 6 | 14 | 14 | 0,01 | 0,01 | 10 |
| Q68FH0 | Pkp4 | 13 | 17 | 16 | 0,01 | 0,01 | 10 |
| E9Q6H8 | Plekha5 | 20 | 29 | 25 | 3 | 2 | 19 |
| Q8VCQ8 | Cald1 | 24 | 31 | 33 | 4 | 10 | 25 |
| Q922V4 | Plrg1 | 20 | 24 | 26 | 3 | 10 | 20 |
| Q60953 | Pml | 20 | 30 | 29 | 10 | 12 | 22 |
| Q99KG5 | Lsr | 17 | 16 | 18 | 5 | 4 | 14 |
| Q3TUQ5 | Pnn | 22 | 27 | 30 | 4 | 11 | 24 |
| O35691 | Pnn | 22 | 27 | 30 | 4 | 11 | 24 |
| A2AJ88 | Pnpla7 | 7 | 14 | 13 | 0,01 | 0,01 | 7 |
| A2AJ88 | Pnpla7 | 7 | 14 | 13 | 0,01 | 0,01 | 7 |
| O35134 | Polr1a | 6 | 17 | 19 | 0,01 | 0,01 | 7 |
| P70700 | Polr1b | 5 | 12 | 10 | 0,01 | 0,01 | 3 |
| P08775 | Polr2a | 34 | 50 | 54 | 4 | 15 | 40 |
| A0A0R4J0V5 | Polr2a | 34 | 50 | 54 | 4 | 15 | 40 |
| Q8CFI7 | Polr2b | 26 | 38 | 35 | 5 | 11 | 30 |
| Q9QZH3 | Ppie | 10 | 12 | 12 | 0,01 | 3 | 6 |

| | | | | | | | |
|---|---|---|---|---|---|---|---|
| Q5FWX6 | Prkd3 | 5 | 8 | 11 | 0,01 | 0,01 | 4 |
| Q8K1Y2 | Prkd3 | 5 | 8 | 11 | 0,01 | 0,01 | 4 |
| Q99KP6 | Prpf19 | 10 | 15 | 16 | 2 | 6 | 12 |
| Q922U1 | Prpf3 | 18 | 23 | 26 | 3 | 14 | 20 |
| Q8CCF0 | Prpf31 | 11 | 12 | 15 | 2 | 4 | 11 |
| Q9DAW6 | Prpf4 | 18 | 20 | 26 | 4 | 12 | 19 |
| Q9R1C7 | Prpf40a | 20 | 23 | 34 | 3 | 6 | 24 |
| Q61136 | Prpf4b | 8 | 15 | 23 | 0,01 | 3 | 12 |
| Q91YR7 | Prpf6 | 29 | 42 | 42 | 14 | 19 | 33 |
| Q99PV0 | Prpf8 | 92 | 109 | 117 | 32 | 53 | 94 |
| Q14C51 | Ptcd3 | 9 | 9 | 9 | 0,01 | 0,01 | 5 |
| Q3UEB3 | Puf60 | 12 | 11 | 15 | 0,01 | 4 | 10 |
| Q3UEB3 | Puf60 | 12 | 11 | 15 | 0,01 | 4 | 10 |
| Q61550 | Rad21 | 12 | 20 | 20 | 1 | 4 | 14 |
| Q8C570 | Rae1 | 11 | 11 | 15 | 1 | 2 | 10 |
| Q9EP71 | Rai14 | 43 | 47 | 52 | 9 | 18 | 44 |
| Q64012 | Raly | 20 | 23 | 25 | 10 | 11 | 20 |
| Q9ERU9 | Ranbp2 | 52 | 84 | 95 | 12 | 28 | 74 |
| Q8BX09 | Rbbp5 | 7 | 13 | 12 | 1 | 2 | 5 |
| Q99KG3 | Rbm10 | 23 | 27 | 32 | 2 | 11 | 26 |
| Q8C2Q3 | Rbm14 | 21 | 27 | 32 | 10 | 16 | 26 |
| Q0VBL3 | Rbm15 | 14 | 14 | 19 | 2 | 7 | 13 |
| B2RY56 | Rbm25 | 22 | 28 | 33 | 6 | 10 | 26 |
| Q8VH51 | Rbm39 | 11 | 15 | 18 | 3 | 6 | 14 |
| Q8VH51 | Rbm39 | 11 | 15 | 18 | 3 | 6 | 14 |
| Q91YE7 | Rbm5 | 17 | 25 | 29 | 5 | 8 | 24 |
| Q91YE7 | Rbm5 | 17 | 25 | 29 | 5 | 8 | 24 |
| S4R1W5 | Rbm6 | 19 | 24 | 30 | 2 | 9 | 25 |
| Q9CQT2 | Rbm7 | 8 | 11 | 11 | 0,01 | 0,01 | 7 |
| Q9WV02 | Rbmx | 16 | 19 | 24 | 11 | 11 | 18 |
| Q91VM5 | Rbmxl1 | 18 | 19 | 26 | 11 | 10 | 19 |
| Q6PR54 | Rif1 | 9 | 20 | 27 | 0,01 | 4 | 21 |
| Q6PR54 | Rif1 | 9 | 20 | 27 | 0,01 | 4 | 21 |
| Q6PR54 | Rif1 | 9 | 20 | 27 | 0,01 | 4 | 21 |
| Q3UJU9 | Rmdn3 | 12 | 16 | 14 | 4 | 3 | 10 |
| P27659 | Rpl3 | 28 | 29 | 29 | 10 | 15 | 24 |
| Q9D8E6 | Rpl4 | 34 | 37 | 40 | 9 | 14 | 34 |
| P47911 | Rpl6 | 21 | 24 | 30 | 11 | 12 | 23 |
| P12970 | Rpl7a | 23 | 25 | 27 | 11 | 13 | 23 |
| Q6P5B0 | Rrp12 | 14 | 22 | 30 | 2 | 3 | 22 |
| Q99LF4 | Rtcb | 12 | 14 | 19 | 0,01 | 1 | 10 |
| P60122 | Ruvbl1 | 21 | 21 | 26 | 8 | 11 | 21 |
| Q9WTM5 | Ruvbl2 | 30 | 30 | 33 | 14 | 21 | 29 |
| D3YXK2 | Safb | 28 | 33 | 39 | 5 | 16 | 29 |
| Q80YR5 | Safb2 | 40 | 47 | 53 | 9 | 21 | 44 |
| Q9ER74 | Sall1 | 5 | 13 | 19 | 0,01 | 1 | 10 |
| Q6P5E3 | Sall1 | 5 | 13 | 19 | 0,01 | 1 | 10 |

| | | | | | | | |
|---|---|---|---|---|---|---|---|
| Q9Z315 | Sart1 | 40 | 48 | 51 | 11 | 21 | 44 |
| E9PZM7 | Scaf11 | 14 | 24 | 26 | 0,01 | 5 | 20 |
| Q80U72 | Scrib | 24 | 34 | 33 | 2 | 4 | 27 |
| Q9EP97 | Senp3 | 5 | 10 | 13 | 0,01 | 2 | 7 |
| E9Q5F9 | Setd2 | 6 | 21 | 33 | 0,01 | 0,01 | 15 |
| E9Q5F9 | Setd2 | 6 | 21 | 33 | 0,01 | 0,01 | 15 |
| Q9D554 | Sf3a3 | 14 | 18 | 19 | 2 | 7 | 12 |
| Q99NB9 | Sf3b1 | 48 | 55 | 58 | 17 | 30 | 53 |
| G5E866 | Sf3b1 | 48 | 55 | 58 | 17 | 30 | 53 |
| Q3UJB0 | Sf3b2 | 31 | 37 | 38 | 10 | 19 | 32 |
| Q921M3 | Sf3b3 | 36 | 42 | 47 | 15 | 24 | 40 |
| Q3USH5 | Sfswap | 5 | 19 | 22 | 0,01 | 3 | 14 |
| Q60520 | Sin3a | 12 | 27 | 22 | 0,01 | 3 | 17 |
| Q60520 | Sin3a | 12 | 27 | 22 | 0,01 | 3 | 17 |
| Q62141 | Sin3b | 6 | 10 | 10 | 0,01 | 0,01 | 0,01 |
| Q62141 | Sin3b | 6 | 10 | 10 | 0,01 | 0,01 | 6 |
| Q9CZU3 | Skiv2l2 | 29 | 33 | 36 | 2 | 9 | 26 |
| Q8CH25 | Sltm | 20 | 28 | 34 | 0,01 | 9 | 25 |
| Q8BHJ9 | Slu7 | 12 | 15 | 18 | 0,01 | 0,01 | 12 |
| Q6DIC0 | Smarca2 | 29 | 34 | 35 | 0,01 | 0,01 | 25 |
| F2Z4A9 | Smarca2 | 29 | 34 | 35 | 0,01 | 0,01 | 25 |
| A0A0R4J170 | Smarca4 | 23 | 29 | 28 | 2 | 7 | 21 |
| Q3TKT4 | Smarca4 | 23 | 29 | 28 | 2 | 7 | 21 |
| Q3TKT4 | Smarca4 | 23 | 29 | 28 | 2 | 7 | 21 |
| P97496 | Smarcc1 | 8 | 17 | 16 | 3 | 0,01 | 10 |
| P97496 | Smarcc1 | 8 | 17 | 16 | 3 | 0,01 | 10 |
| Q99JR8 | Smarcd2 | 9 | 17 | 16 | 0,01 | 0,01 | 7 |
| Q99JR8 | Smarcd2 | 9 | 17 | 16 | 0,01 | 0,01 | 7 |
| O54941 | Smarce1 | 9 | 13 | 13 | 2 | 4 | 8 |
| Q9CU62 | Smc1a | 41 | 57 | 58 | 9 | 25 | 52 |
| Q9CW03 | Smc3 | 41 | 49 | 53 | 9 | 19 | 41 |
| Q3UKJ7 | Smu1 | 11 | 16 | 19 | 2 | 6 | 12 |
| Q6P4T2 | Snrnp200 | 69 | 90 | 103 | 30 | 50 | 91 |
| Q6PE01 | Snrnp40 | 9 | 14 | 16 | 2 | 4 | 10 |
| A0A0B4J1E2 | Snw1 | 18 | 25 | 24 | 4 | 8 | 19 |
| A2ADB0 | Spen | 13 | 26 | 35 | 0,01 | 2 | 26 |
| Q62261 | Sptbn1 | 123 | 136 | 140 | 44 | 71 | 131 |
| A0A087WNL7 | Srcap | 5 | 12 | 11 | 0,01 | 0,01 | 6 |
| A0A087WQ44 | Srcap | 5 | 12 | 11 | 0,01 | 0,01 | 6 |
| A0A087WNX7 | Srcap | 5 | 12 | 11 | 0,01 | 0,01 | 6 |
| Q922B9 | Ssfa2 | 17 | 22 | 22 | 2 | 2 | 17 |
| Q9D3E6 | Stag1 | 12 | 23 | 26 | 0,01 | 4 | 9 |
| O35638 | Stag2 | 14 | 21 | 18 | 0,01 | 0,01 | 11 |
| A2AFF6 | Stag2 | 14 | 21 | 18 | 0,01 | 0,01 | 11 |
| Q9D666 | Sun1 | 6 | 10 | 10 | 1 | 2 | 5 |
| O55201 | Supt5h | 13 | 27 | 32 | 0,01 | 4 | 19 |
| Q9D198 | Syf2 | 8 | 9 | 12 | 0,01 | 1 | 6 |

| | | | | | | | |
|---|---|---|---|---|---|---|---|
| Q6ZWQ0 | Syne2 | 97 | 187 | 235 | 6 | 38 | 154 |
| Q80UV9 | Taf1 | 14 | 30 | 30 | 0,01 | 6 | 17 |
| Q80UV9 | Taf1 | 14 | 30 | 30 | 0,01 | 6 | 17 |
| E9QAP7 | Taf4 | 9 | 12 | 14 | 1 | 3 | 10 |
| F8VPY2 | Taf5 | 10 | 19 | 20 | 2 | 8 | 16 |
| Q62311 | Taf6 | 17 | 24 | 27 | 1 | 6 | 16 |
| Q9R1C0 | Taf7 | 6 | 8 | 11 | 1 | 2 | 3 |
| Q8CGF7 | Tcerg1 | 9 | 16 | 20 | 0,01 | 1 | 9 |
| Q8CGF7 | Tcerg1 | 9 | 16 | 20 | 0,01 | 1 | 9 |
| Q9ERA6 | Tfip11 | 21 | 26 | 33 | 0,01 | 2 | 28 |
| B1AZI6 | Thoc2 | 7 | 12 | 17 | 0,01 | 2 | 8 |
| Q569Z6 | Thrap3 | 20 | 30 | 31 | 3 | 12 | 27 |
| P39447 | Tjp1 | 59 | 62 | 71 | 17 | 25 | 61 |
| B9EHJ3 | Tjp1 | 58 | 61 | 70 | 17 | 26 | 60 |
| Q9Z0U1 | Tjp2 | 35 | 43 | 41 | 7 | 16 | 36 |
| P58871 | Tnks1bp1 | 29 | 47 | 53 | 2 | 8 | 40 |
| Q04750 | Top1 | 23 | 23 | 21 | 15 | 9 | 17 |
| Q01320 | Top2a | 13 | 20 | 26 | 4 | 11 | 20 |
| Q64511 | Top2b | 39 | 55 | 63 | 9 | 23 | 44 |
| Q921T2 | Tor1aip1 | 13 | 16 | 22 | 4 | 6 | 17 |
| P70399 | Tp53bp1 | 12 | 25 | 27 | 0,01 | 3 | 15 |
| P70399 | Tp53bp1 | 12 | 25 | 27 | 0,01 | 3 | 15 |
| Q7M739 | Tpr | 23 | 29 | 33 | 2 | 6 | 29 |
| F6ZDS4 | Tpr | 23 | 29 | 33 | 2 | 6 | 29 |
| Q99KW3 | Triobp | 8 | 16 | 14 | 0,01 | 0,01 | 9 |
| Q99KW3 | Triobp | 8 | 16 | 14 | 0,01 | 0,01 | 9 |
| Q99KW3 | Triobp | 8 | 16 | 14 | 0,01 | 0,01 | 9 |
| G5E870 | Trip12 | 5 | 28 | 34 | 0,01 | 3 | 22 |
| A2ASS6 | Ttn | 61 | 138 | 185 | 17 | 61 | 0,01 |
| P68372 | Tubb4b | 17 | 17 | 21 | 5 | 0,01 | 16 |
| P99024 | Tubb5 | 15 | 16 | 20 | 7 | 9 | 15 |
| E9QJS1 | Tyk2 | 9 | 12 | 14 | 0,01 | 0,01 | 8 |
| Q9R117 | Tyk2 | 9 | 12 | 14 | 0,01 | 0,01 | 8 |
| Q6NV83 | U2surp | 27 | 34 | 36 | 6 | 14 | 30 |
| Q6NV83 | U2surp | 27 | 34 | 36 | 6 | 14 | 30 |
| Q811S7 | Ubp1 | 13 | 17 | 19 | 2 | 4 | 14 |
| Q9EPU0 | Upf1 | 16 | 27 | 22 | 0,01 | 0,01 | 17 |
| Q9EPU0 | Upf1 | 16 | 27 | 22 | 0,01 | 0,01 | 17 |
| Q571H0 | Urb1 | 45 | 53 | 72 | 3 | 13 | 55 |
| E9Q7L1 | Urb2 | 17 | 28 | 34 | 6 | 14 | 29 |
| Q3TIX9 | Usp39 | 13 | 18 | 17 | 1 | 3 | 12 |
| Q8C7V3 | Utp15 | 19 | 20 | 24 | 4 | 8 | 17 |
| Q5XG71 | Utp20 | 48 | 56 | 72 | 2 | 16 | 61 |
| E9QK83 | Utp20 | 48 | 56 | 72 | 2 | 16 | 61 |
| E9Q6R7 | Utrn | 121 | 139 | 150 | 25 | 60 | 128 |
| Q01853 | Vcp | 14 | 17 | 23 | 1 | 4 | 14 |
| E9PZY8 | Virma | 9 | 19 | 26 | 0,01 | 0,01 | 17 |

| | | | | | | | |
|---|---|---|---|---|---|---|---|
| A2AIV2 | Virma | 9 | 19 | 26 | 0,01 | 0,01 | 17 |
| Q923D5 | Wbp11 | 4 | 6 | 17 | 0,01 | 2 | 6 |
| Q8K4P0 | Wdr33 | 5 | 15 | 21 | 0,01 | 4 | 13 |
| Q3TA68 | Wdr36 | 20 | 28 | 29 | 6 | 11 | 24 |
| Q3TAQ9 | Wdr36 | 20 | 28 | 29 | 6 | 11 | 24 |
| Q6ZQL4 | Wdr43 | 13 | 18 | 20 | 0,01 | 6 | 16 |
| Q3U821 | Wdr75 | 14 | 16 | 25 | 3 | 6 | 20 |
| Q9DCD2 | Xab2 | 17 | 22 | 27 | 0,01 | 7 | 22 |
| Q9DBR1 | Xrn2 | 6 | 11 | 16 | 0,01 | 0,01 | 7 |
| Q9DBR1 | Xrn2 | 6 | 11 | 16 | 0,01 | 0,01 | 7 |
| D3YWX2 | Ylpm1 | 19 | 33 | 45 | 0,01 | 6 | 27 |
| E9Q5K9 | Ythdc1 | 9 | 13 | 17 | 0,01 | 4 | 12 |
| E9Q5K9 | Ythdc1 | 9 | 13 | 17 | 0,01 | 4 | 12 |
| E9Q5K9 | Ythdc1 | 9 | 13 | 17 | 0,01 | 4 | 12 |
| Q8K0L9 | Zbtb20 | 13 | 20 | 20 | 0,01 | 3 | 15 |
| Q8K0L9 | Zbtb20 | 13 | 20 | 20 | 0,01 | 3 | 15 |
| E9Q784 | Zc3h13 | 4 | 19 | 24 | 0,01 | 0,01 | 14 |
| Q8BJ05 | Zc3h14 | 12 | 21 | 24 | 0,01 | 5 | 15 |
| Q9CYA6 | Zcchc8 | 9 | 11 | 15 | 0,01 | 2 | 8 |
| Q8C827 | Zfp62 | 14 | 23 | 23 | 0,01 | 6 | 19 |
| Q8C827 | Zfp62 | 14 | 23 | 23 | 0,01 | 6 | 19 |
| E9QML5 | Zfp638 | 18 | 45 | 55 | 0,01 | 10 | 35 |
| O88532 | Zfr | 11 | 19 | 21 | 0,01 | 0,01 | 12 |
| A2A483 | Zmynd8 | 11 | 17 | 23 | 0,01 | 3 | 16 |
| Q3UH28 | Zmynd8 | 11 | 17 | 23 | 0,01 | 3 | 16 |
| E9Q8D1 | Zmynd8 | 11 | 17 | 23 | 0,01 | 3 | 16 |
| Q3U1M7 | Zmynd8 | 11 | 17 | 23 | 0,01 | 3 | 16 |
| Q61624 | Znf148 | 4 | 12 | 18 | 0,01 | 4 | 9 |
