## Supplemental table 3 for "The Splicing Factor XAB2 interacts with ERCC1-XPF and XPG for RNA-loop processing during mammalian development"

**Table S3. A list of 255 differentially expressed genes in HEPA cells transfected with dsRNA targeting the *Xab2* transcript vs. scramble control cells**

| Gene ID | Gene symbol | FC | P value | Q value |
|---|---|---|---|---|
| XLOC_024651 | Arglu1 | 5,493774 | 5,00E-05 | 0,00581 |
| XLOC_000040 | Gsta3 | 2,790866 | 5,00E-05 | 0,00581 |
| XLOC_008926 | Kmt2d | 2,585648 | 5,00E-05 | 0,00581 |
| XLOC_002682 | Mbd6 | 2,577908 | 5,00E-05 | 0,00581 |
| XLOC_005261 | Mir8099-1 | 2,437726 | 5,00E-05 | 0,00581 |
| XLOC_024417 | Nlrc5 | 2,114796 | 5,00E-05 | 0,00581 |
| XLOC_013555 | Lrp4 | 2,055607 | 5,00E-05 | 0,00581 |
| XLOC_022045 | Lmtk3 | 1,943463 | 5,00E-05 | 0,00581 |
| XLOC_018507 | Cacna2d1 | 1,858967 | 5,00E-05 | 0,00581 |
| XLOC_026663 | Ptpn23 | 1,79707 | 5,00E-05 | 0,00581 |
| XLOC_016963 | Col27a1 | 1,755826 | 5,00E-05 | 0,00581 |
| XLOC_017857 | Szt2 | 1,673532 | 5,00E-05 | 0,00581 |
| XLOC_020480 | Kbtbd8 | 1,608034 | 5,00E-05 | 0,00581 |
| XLOC_025836 | Gsta4 | 1,604497 | 0,0002 | 0,016562 |
| XLOC_006331 | Lhfpl2 | 1,58489 | 0,0001 | 0,009797 |
| XLOC_018736 | Ugt2b35 | 1,567962 | 5,00E-05 | 0,00581 |
| XLOC_005623 | Zfyve26 | 1,553334 | 5,00E-05 | 0,00581 |
| XLOC_000021 | Xkr9 | 1,535324 | 0,0003 | 0,022372 |
| XLOC_022746 | Gltscr1 | 1,530128 | 5,00E-05 | 0,00581 |
| XLOC_019430 | Ugt2b34 | 1,491004 | 5,00E-05 | 0,00581 |
| XLOC_016830 | Mdn1 | 1,485966 | 5,00E-05 | 0,00581 |
| XLOC_008711 | BC024139,Eppk1 | 1,44005 | 5,00E-05 | 0,00581 |
| XLOC_014615 | Prr5l | 1,439003 | 5,00E-05 | 0,00581 |
| XLOC_014061 | Ss18l1 | 1,435786 | 0,00095 | 0,047465 |
| XLOC_027448 | Clcn5 | 1,43536 | 5,00E-05 | 0,00581 |
| XLOC_012449 | Lcor | 1,434183 | 5,00E-05 | 0,00581 |
| XLOC_018944 | Gm15800 | 1,432854 | 5,00E-05 | 0,00581 |
| XLOC_001101 | Lamc2 | 1,427963 | 5,00E-05 | 0,00581 |
| XLOC_007943 | Abcc4 | 1,422403 | 5,00E-05 | 0,00581 |
| XLOC_026981 | Maoa | 1,419764 | 5,00E-05 | 0,00581 |
| XLOC_017062 | Usp24 | 1,415167 | 5,00E-05 | 0,00581 |
| XLOC_011820 | Nrep | 1,408701 | 0,0005 | 0,032871 |
| XLOC_019713 | Cldn4 | 1,408596 | 5,00E-05 | 0,00581 |
| XLOC_014755 | Cep152 | 1,402274 | 5,00E-05 | 0,00581 |
| XLOC_022963 | Kmt2b | 1,400677 | 5,00E-05 | 0,00581 |
| XLOC_010712 | Igf2r | 1,398307 | 0,00015 | 0,013544 |
| XLOC_026949 | Xk | 1,396522 | 0,00055 | 0,03435 |
| XLOC_022366 | Ampd3 | 1,391766 | 5,00E-05 | 0,00581 |
| XLOC_000809 | Fam126b | 1,391417 | 5,00E-05 | 0,00581 |
| XLOC_027346 | Vsig1 | 1,386022 | 0,0001 | 0,009797 |
| XLOC_017307 | Hspg2 | 1,380663 | 5,00E-05 | 0,00581 |
| XLOC_016055 | Slc7a11 | 1,379557 | 5,00E-05 | 0,00581 |
| XLOC_025789 | Vps13c | 1,37503 | 5,00E-05 | 0,00581 |
| XLOC_003308 | Per1 | 1,370714 | 5,00E-05 | 0,00581 |

| | | | | |
|---|---|---|---|---|
| XLOC_020684 | Mgst1 | 1,36819 | 5,00E-05 | 0,00581 |
| XLOC_006435 | Gpr137b | 1,367162 | 0,0009 | 0,046358 |
| XLOC_018597 | Depdc5 | 1,364187 | 0,00035 | 0,024863 |
| XLOC_009821 | Plcxd2 | 1,359702 | 5,00E-05 | 0,00581 |
| XLOC_003187 | Fnip1 | 1,357773 | 5,00E-05 | 0,00581 |
| XLOC_011902 | Cep120 | 1,355205 | 5,00E-05 | 0,00581 |
| XLOC_009352 | Map3k13 | 1,349688 | 0,0003 | 0,022372 |
| XLOC_019058 | Gigyf1 | 1,346598 | 5,00E-05 | 0,00581 |
| XLOC_008878 | Gxylt1 | 1,343211 | 5,00E-05 | 0,00581 |
| XLOC_017329 | Ubr4 | 1,328262 | 5,00E-05 | 0,00581 |
| XLOC_024318 | Hmox1 | 1,327338 | 5,00E-05 | 0,00581 |
| XLOC_000272 | Spp2 | 1,3259 | 0,0003 | 0,022372 |
| XLOC_022076 | Herc2 | 1,323041 | 5,00E-05 | 0,00581 |
| XLOC_000432 | Dennd1b | 1,322424 | 5,00E-05 | 0,00581 |
| XLOC_021898 | Zfp568 | 1,315545 | 5,00E-05 | 0,00581 |
| XLOC_023213 | Chd2 | 1,314641 | 0,00025 | 0,019827 |
| XLOC_006116 | Serpinb9b | 1,314554 | 5,00E-05 | 0,00581 |
| XLOC_000775 | Slc40a1 | 1,312906 | 5,00E-05 | 0,00581 |
| XLOC_019427 | Tmprss11f | 1,312184 | 0,00015 | 0,013544 |
| XLOC_027257 | Atp7a | 1,309026 | 0,0001 | 0,009797 |
| XLOC_013985 | Ctsa | 1,308774 | 0,00015 | 0,013544 |
| XLOC_012748 | Vps13a | 1,307794 | 5,00E-05 | 0,00581 |
| XLOC_027336 | Trap1a | 1,303909 | 5,00E-05 | 0,00581 |
| XLOC_022423 | Tnrc6a | 1,299229 | 5,00E-05 | 0,00581 |
| XLOC_011491 | Dsg2 | 1,29913 | 5,00E-05 | 0,00581 |
| XLOC_016225 | Sprr1b | 1,297888 | 5,00E-05 | 0,00581 |
| XLOC_018109 | Vps13d | 1,297219 | 0,0004 | 0,027503 |
| XLOC_027335 | Mum1l1 | 1,296291 | 0,0001 | 0,009797 |
| XLOC_000169 | Pikfyve | 1,296196 | 0,0001 | 0,009797 |
| XLOC_010845 | Dusp1 | 1,295962 | 5,00E-05 | 0,00581 |
| XLOC_016792 | Trp53inp1 | 1,294208 | 5,00E-05 | 0,00581 |
| XLOC_026625 | Sema3b | 1,293413 | 0,00095 | 0,047465 |
| XLOC_010812 | Mapk8ip3 | 1,292875 | 0,00015 | 0,013544 |
| XLOC_003479 | Ccl2 | 1,292831 | 0,0001 | 0,009797 |
| XLOC_020275 | Gpnmb | 1,290966 | 5,00E-05 | 0,00581 |
| XLOC_009427 | Iqcb1 | 1,290603 | 0,00045 | 0,030247 |
| XLOC_023585 | Cln3 | 1,290531 | 0,0002 | 0,016562 |
| XLOC_011181 | Ankrd12 | 1,289866 | 0,0001 | 0,009797 |
| XLOC_009982 | Tmprss2 | 1,289174 | 0,0001 | 0,009797 |
| XLOC_023092 | Ftl1 | 1,287422 | 0,0001 | 0,009797 |
| XLOC_018614 | Htt | 1,285363 | 0,0002 | 0,016562 |
| XLOC_006341 | Gcnt4 | 1,284224 | 0,00055 | 0,03435 |
| XLOC_002173 | Ankrd52 | 1,284211 | 0,0002 | 0,016562 |
| XLOC_016368 | Csf1 | 1,278774 | 5,00E-05 | 0,00581 |
| XLOC_013130 | Arl5b | 1,278176 | 0,00035 | 0,024863 |
| XLOC_010248 | Clcn7 | 1,27778 | 0,00025 | 0,019827 |
| XLOC_015063 | Lama5 | 1,277136 | 0,0001 | 0,009797 |

| | | | | |
|---|---|---|---|---|
| XLOC_001185 | Uhmk1 | 1,275072 | 5,00E-05 | 0,00581 |
| XLOC_006866 | Zswim6 | 1,273725 | 0,0001 | 0,009797 |
| XLOC_024867 | Ifi30 | 1,273377 | 0,00035 | 0,024863 |
| XLOC_006015 | Elmo1 | 1,27002 | 0,0005 | 0,032871 |
| XLOC_009003 | Itga5 | 1,270012 | 0,00025 | 0,019827 |
| XLOC_010980 | Vars2 | 1,269703 | 0,0003 | 0,022372 |
| XLOC_024169 | Ppp1r3b | 1,267551 | 0,0005 | 0,032871 |
| XLOC_001729 | Rev3l | 1,266697 | 0,0007 | 0,039297 |
| XLOC_004995 | Dnmt3a | 1,265839 | 0,00045 | 0,030247 |
| XLOC_020984 | Atoh8 | 1,262073 | 0,00045 | 0,030247 |
| XLOC_016954 | Snx30 | 1,260936 | 0,0002 | 0,016562 |
| XLOC_008560 | Trio | 1,260702 | 0,0001 | 0,009797 |
| XLOC_001742 | Ostm1 | 1,259171 | 0,00035 | 0,024863 |
| XLOC_017753 | Jun | 1,258725 | 0,0006 | 0,036118 |
| XLOC_014303 | Zbtb26 | 1,256416 | 0,0004 | 0,027503 |
| XLOC_013154 | Acbd5 | 1,256192 | 0,0006 | 0,036118 |
| XLOC_018757 | Slc4a4 | 1,256003 | 0,0002 | 0,016562 |
| XLOC_009984 | Prdm15 | 1,25379 | 0,00055 | 0,03435 |
| XLOC_003372 | Zzef1 | 1,252404 | 0,0009 | 0,046358 |
| XLOC_005130 | Fam179b | 1,250555 | 0,0009 | 0,046358 |
| XLOC_025888 | Xrn1 | 1,249861 | 0,00075 | 0,04133 |
| XLOC_020991 | Tgoln1 | 1,248898 | 0,00075 | 0,04133 |
| XLOC_003067 | Cpeb4 | 1,248767 | 0,0008 | 0,043604 |
| XLOC_015620 | Gba | 1,24312 | 0,0003 | 0,022372 |
| XLOC_000831 | Fzd5 | 1,240482 | 0,00075 | 0,04133 |
| XLOC_014835 | Flrt3 | 1,239705 | 0,00075 | 0,04133 |
| XLOC_008671 | Ago2 | 1,237956 | 0,0003 | 0,022372 |
| XLOC_007203 | Ercc6 | 1,237761 | 0,0008 | 0,043604 |
| XLOC_002169 | Stat2 | 1,236765 | 0,00085 | 0,045665 |
| XLOC_000527 | Olfml2b | 1,235798 | 0,0006 | 0,036118 |
| XLOC_000152 | Nbeal1 | 1,232953 | 0,00055 | 0,03435 |
| XLOC_007140 | Zswim8 | 1,226879 | 0,0009 | 0,046358 |
| XLOC_019125 | Trrap | 1,222326 | 0,00055 | 0,03435 |
| XLOC_019654 | Camkk2 | -1,20376 | 0,00055 | 0,03435 |
| XLOC_016880 | Ccdc107 | -1,21111 | 0,00085 | 0,045665 |
| XLOC_022093 | Tarsl2 | -1,22344 | 0,00065 | 0,037763 |
| XLOC_027434 | Arhgap6 | -1,22856 | 0,00055 | 0,03435 |
| XLOC_023705 | Ifitm3 | -1,22894 | 0,0007 | 0,039297 |
| XLOC_010419 | H2-Q1 | -1,23057 | 0,00035 | 0,024863 |
| XLOC_023256 | Btbd1 | -1,23403 | 0,00065 | 0,037763 |
| XLOC_019480 | Antxr2 | -1,23838 | 0,0007 | 0,039297 |
| XLOC_012402 | Papss2 | -1,23967 | 0,00095 | 0,047465 |
| XLOC_026398 | Lingo1 | -1,24033 | 0,00045 | 0,030247 |
| XLOC_018489 | Abcb1b | -1,24221 | 0,0006 | 0,036118 |
| XLOC_014354 | Grb14 | -1,24339 | 0,0002 | 0,016562 |
| XLOC_014881 | Sox12 | -1,24567 | 0,00085 | 0,045665 |
| XLOC_003620 | Cisd3 | -1,24802 | 0,00055 | 0,03435 |

| | | | | |
|---|---|---|---|---|
| XLOC_013229 | Rapgef1 | -1,2529 | 0,00015 | 0,013544 |
| XLOC_019453 | Btc | -1,25334 | 0,0004 | 0,027503 |
| XLOC_010867 | Fkbp5 | -1,25482 | 0,0003 | 0,022372 |
| XLOC_004745 | Faap100 | -1,25511 | 0,00015 | 0,013544 |
| XLOC_012669 | Uqcc3 | -1,25594 | 0,0006 | 0,036118 |
| XLOC_024291 | Slc27a1 | -1,25636 | 0,0009 | 0,046358 |
| XLOC_023518 | Galnt18 | -1,25668 | 0,00015 | 0,013544 |
| XLOC_017681 | Rnf183 | -1,25714 | 0,00035 | 0,024863 |
| XLOC_004564 | Aarsd1 | -1,25749 | 0,0003 | 0,022372 |
| XLOC_013063 | Itih5 | -1,25799 | 0,0001 | 0,009797 |
| XLOC_024334 | Inpp4b | -1,258 | 0,0001 | 0,009797 |
| XLOC_020814 | Podxl | -1,25823 | 0,0007 | 0,039297 |
| XLOC_000904 | Pid1 | -1,25986 | 0,0005 | 0,032871 |
| XLOC_026554 | 1190002N15Rik | -1,26041 | 0,00035 | 0,024863 |
| XLOC_012394 | Il33 | -1,26332 | 0,00055 | 0,03435 |
| XLOC_010924 | Angptl4 | -1,26643 | 0,00075 | 0,04133 |
| XLOC_019782 | Psmg3 | -1,26695 | 0,0009 | 0,046358 |
| XLOC_005094 | Foxg1 | -1,26852 | 0,00065 | 0,037763 |
| XLOC_009911 | Adamts1 | -1,26854 | 0,0009 | 0,046358 |
| XLOC_019338 | Qdpr | -1,27211 | 5,00E-05 | 0,00581 |
| XLOC_015417 | Bhlhe22 | -1,27223 | 0,0004 | 0,027503 |
| XLOC_013275 | Ptges2 | -1,27357 | 0,0006 | 0,036118 |
| XLOC_005752 | Ccdc85c | -1,27378 | 0,00055 | 0,03435 |
| XLOC_025617 | Pvrl1 | -1,27421 | 0,00065 | 0,037763 |
| XLOC_024879 | Nr2f6 | -1,27526 | 0,00025 | 0,019827 |
| XLOC_024568 | Irf8 | -1,27637 | 0,0004 | 0,027503 |
| XLOC_004759 | Pycr1 | -1,27802 | 0,0002 | 0,016562 |
| XLOC_016443 | Enpep | -1,2848 | 5,00E-05 | 0,00581 |
| XLOC_003648 | Igfbp4 | -1,29022 | 0,0001 | 0,009797 |
| XLOC_024938 | Nfix | -1,29184 | 5,00E-05 | 0,00581 |
| XLOC_002545 | Cry1 | -1,29276 | 0,00065 | 0,037763 |
| XLOC_019327 | Hs3st1 | -1,29297 | 0,0001 | 0,009797 |
| XLOC_009707 | Etv5 | -1,29736 | 0,0002 | 0,016562 |
| XLOC_003691 | Tmem106a | -1,30111 | 0,00095 | 0,047465 |
| XLOC_005480 | 1700030C10Rik | -1,30183 | 0,00085 | 0,045665 |
| XLOC_010829 | Metrn | -1,30319 | 0,0007 | 0,039297 |
| XLOC_004406 | Stxbp4 | -1,30426 | 0,00065 | 0,037763 |
| XLOC_001099 | Rgl1 | -1,30833 | 0,0004 | 0,027503 |
| XLOC_012917 | Shtn1 | -1,30889 | 5,00E-05 | 0,00581 |
| XLOC_022953 | Sdhaf1 | -1,3094 | 0,0001 | 0,009797 |
| XLOC_011977 | Ctif | -1,31107 | 0,0002 | 0,016562 |
| XLOC_012460 | Marveld1 | -1,31284 | 5,00E-05 | 0,00581 |
| XLOC_014200 | Agpat2 | -1,31857 | 5,00E-05 | 0,00581 |
| XLOC_003740 | Tcam1 | -1,32276 | 5,00E-05 | 0,00581 |
| XLOC_023626 | Bcl7c | -1,32373 | 0,00015 | 0,013544 |
| XLOC_017266 | Map3k6 | -1,32992 | 0,0006 | 0,036118 |
| XLOC_008755 | Il2rb | -1,33001 | 5,00E-05 | 0,00581 |

| | | | | |
|---|---|---|---|---|
| XLOC_004666 | Cd300lb | -1,33065 | 5,00E-05 | 0,00581 |
| XLOC_001318 | G0s2 | -1,33151 | 0,0001 | 0,009797 |
| XLOC_009804 | Tigit | -1,33194 | 5,00E-05 | 0,00581 |
| XLOC_022220 | Lipt2 | -1,33285 | 5,00E-05 | 0,00581 |
| XLOC_022452 | Zfp771 | -1,3348 | 0,00015 | 0,013544 |
| XLOC_009611 | Dexi | -1,33516 | 0,0003 | 0,022372 |
| XLOC_022437 | Ccdc101 | -1,33528 | 0,0002 | 0,016562 |
| XLOC_004246 | Spns2 | -1,33602 | 0,0003 | 0,022372 |
| XLOC_016969 | 6330416G13Rik | -1,33835 | 5,00E-05 | 0,00581 |
| XLOC_022750 | Inafm1 | -1,34141 | 0,0009 | 0,046358 |
| XLOC_019482 | Prkg2 | -1,34323 | 5,00E-05 | 0,00581 |
| XLOC_021212 | Grcc10 | -1,34412 | 5,00E-05 | 0,00581 |
| XLOC_003051 | Fancl | -1,34449 | 0,00035 | 0,024863 |
| XLOC_003919 | Igfbp3 | -1,349 | 5,00E-05 | 0,00581 |
| XLOC_006591 | Mcur1 | -1,35569 | 5,00E-05 | 0,00581 |
| XLOC_018828 | Lrrc8b | -1,35616 | 5,00E-05 | 0,00581 |
| XLOC_009650 | Sdf2l1 | -1,35769 | 0,00015 | 0,013544 |
| XLOC_021829 | Tgfb1 | -1,35833 | 5,00E-05 | 0,00581 |
| XLOC_006828 | Tmem171 | -1,36459 | 5,00E-05 | 0,00581 |
| XLOC_019492 | Plac8 | -1,36711 | 5,00E-05 | 0,00581 |
| XLOC_000292 | Twist2 | -1,37114 | 5,00E-05 | 0,00581 |
| XLOC_009385 | Cep19 | -1,38222 | 0,0002 | 0,016562 |
| XLOC_023008 | Kctd15 | -1,38716 | 0,0001 | 0,009797 |
| XLOC_024578 | Zfpm1 | -1,38907 | 5,00E-05 | 0,00581 |
| XLOC_020448 | Txnrd3 | -1,39076 | 5,00E-05 | 0,00581 |
| XLOC_004677 | Fdxr | -1,39135 | 0,0004 | 0,027503 |
| XLOC_008876 | Abcd2 | -1,39297 | 5,00E-05 | 0,00581 |
| XLOC_013163 | Il1rn | -1,39331 | 5,00E-05 | 0,00581 |
| XLOC_007920 | Tbc1d4 | -1,39678 | 0,00025 | 0,019827 |
| XLOC_026302 | Oaf | -1,40583 | 5,00E-05 | 0,00581 |
| XLOC_009259 | Rbfox1 | -1,41625 | 0,0006 | 0,036118 |
| XLOC_021223 | Ptms | -1,42134 | 5,00E-05 | 0,00581 |
| XLOC_021068 | H1fx | -1,42328 | 5,00E-05 | 0,00581 |
| XLOC_010899 | Notch3 | -1,42502 | 0,00035 | 0,024863 |
| XLOC_014925 | Scand1 | -1,44548 | 5,00E-05 | 0,00581 |
| XLOC_014373 | Mettl8 | -1,45232 | 5,00E-05 | 0,00581 |
| XLOC_020504 | Lmcd1 | -1,45715 | 5,00E-05 | 0,00581 |
| XLOC_013579 | Lrrc4c | -1,4692 | 0,0009 | 0,046358 |
| XLOC_000478 | Tnr | -1,46958 | 5,00E-05 | 0,00581 |
| XLOC_022758 | Calm3 | -1,47282 | 5,00E-05 | 0,00581 |
| XLOC_004539 | Hap1 | -1,47942 | 5,00E-05 | 0,00581 |
| XLOC_025624 | H2afx | -1,48535 | 5,00E-05 | 0,00581 |
| XLOC_005401 | Tnfaip2 | -1,49665 | 5,00E-05 | 0,00581 |
| XLOC_000117 | Osgepl1 | -1,50134 | 5,00E-05 | 0,00581 |
| XLOC_014910 | Ggt7 | -1,50398 | 0,0001 | 0,009797 |
| XLOC_025406 | Mmp10 | -1,50667 | 5,00E-05 | 0,00581 |
| XLOC_002587 | Socs2 | -1,52788 | 5,00E-05 | 0,00581 |

| | | | | |
|---|---|---|---|---|
| XLOC_016216 | Npr1 | -1,53653 | 0,001 | 0,049632 |
| XLOC_007173 | Wnt5a | -1,54269 | 5,00E-05 | 0,00581 |
| XLOC_009695 | Clcn2 | -1,55145 | 0,0003 | 0,022372 |
| XLOC_021720 | Pglyrp1 | -1,56003 | 5,00E-05 | 0,00581 |
| XLOC_019322 | Slc2a9 | -1,56081 | 0,0009 | 0,046358 |
| XLOC_005593 | Gpr135,L3hypdh | -1,57489 | 0,0001 | 0,009797 |
| XLOC_008330 | H1f0 | -1,58099 | 5,00E-05 | 0,00581 |
| XLOC_023537 | Syt17 | -1,58359 | 0,00025 | 0,019827 |
| XLOC_012242 | Rcor2 | -1,59586 | 5,00E-05 | 0,00581 |
| XLOC_017708 | Ptprd | -1,60272 | 5,00E-05 | 0,00581 |
| XLOC_000339 | Serpinb2 | -1,63868 | 5,00E-05 | 0,00581 |
| XLOC_020096 | Ppp1r9a | -1,66783 | 5,00E-05 | 0,00581 |
| XLOC_016370 | Gstm7 | -1,68108 | 5,00E-05 | 0,00581 |
| XLOC_004761 | Notum | -1,69305 | 5,00E-05 | 0,00581 |
| XLOC_014775 | Mal | -1,69854 | 5,00E-05 | 0,00581 |
| XLOC_026190 | Epor | -1,71145 | 0,0009 | 0,046358 |
| XLOC_001061 | Pkp1 | -1,74101 | 5,00E-05 | 0,00581 |
| XLOC_010531 | Trerf1 | -1,74822 | 5,00E-05 | 0,00581 |
| XLOC_004525 | Krt33b | -1,86845 | 0,00095 | 0,047465 |
| XLOC_016058 | Ndufc1 | -2,16564 | 5,00E-05 | 0,00581 |
| XLOC_024633 | Xab2 | -2,51335 | 5,00E-05 | 0,00581 |
| XLOC_017267 | Slc9a1 | -6,86589 | 5,00E-05 | 0,00581 |
| XLOC_012228 | Mir192,Mir194-2 | -76,5398 | 5,00E-05 | 0,00581 |
