## Supplemental table 4 for "The Splicing Factor XAB2 interacts with ERCC1-XPF and XPG for RNA-loop processing during mammalian development"

**Table S4. A list of 333 differentially expressed genes in mES cells transfected with dsRNA targeting the *Xab2* transcript vs. scramble control cells**

| Gene ID | Gene symbol | FC | P value | Q value |
|---|---|---|---|---|
| XLOC_039582 | Gm21978 | ND in Scrbl cells | 0,00445 | 0,039699 |
| XLOC_024521 | Gm9959 | ND in Scrbl cells | 0,00345 | 0,031132 |
| XLOC_021960 | Gm23332 | ND in Scrbl cells | 0,00305 | 0,027701 |
| XLOC_012490 | Ighv1-60 | ND in Scrbl cells | 0,0022 | 0,020359 |
| XLOC_027409 | Scarna17 | ND in Scrbl cells | 0,00215 | 0,019965 |
| XLOC_046725 | 1700065J11Rik | ND in Scrbl cells | 0,00175 | 0,018436 |
| XLOC_031461 | Gm16355 | ND in Scrbl cells | 0,0017 | 0,018436 |
| XLOC_062989 | Gm15378 | ND in Scrbl cells | 0,0015 | 0,01775 |
| XLOC_014722 | Gm9742 | ND in Scrbl cells | 0,00145 | 0,01775 |
| XLOC_064030 | Gm14614 | ND in Scrbl cells | 0,001 | 0,01775 |
| XLOC_018145 | Ube2e2 | 13805,92565 | 5,00E-05 | 0,004846 |
| XLOC_031542 | Uckl1os | 28,79786594 | 0,00135 | 0,01775 |
| XLOC_050684 | Rnf225 | 16,70073856 | 5,00E-05 | 0,004846 |
| XLOC_034754 | Gm27403,Gm27611 | 6,900782679 | 0,0045 | 0,040089 |
| XLOC_010057 | Cep131 | 6,692382944 | 5,00E-05 | 0,004846 |
| XLOC_037000 | 4933431E20Rik | 6,437542269 | 0,00055 | 0,01775 |
| XLOC_032450 | Cd82 | 5,290595065 | 0,00055 | 0,01775 |
| XLOC_047412 | Gata2 | 3,902260257 | 0,0001 | 0,009173 |
| XLOC_048031 | Arntl2 | 3,720973201 | 0,00285 | 0,025922 |
| XLOC_011292 | AC099934.1 | 3,641745236 | 5,00E-05 | 0,004846 |
| XLOC_020851 | Kansl2 | 3,144803774 | 5,00E-05 | 0,004846 |
| XLOC_031779 | Snhg7,Snora17 | 2,924500022 | 0,001 | 0,01775 |
| XLOC_047707 | A2m | 2,796476099 | 0,0002 | 0,0177 |
| XLOC_052192 | RP23-370I17.3 | 2,401357168 | 0,00175 | 0,018436 |
| XLOC_031529 | Birc7 | 2,388508482 | 0,0027 | 0,024623 |
| XLOC_035717 | Pitx2 | 2,168563968 | 0,00275 | 0,025055 |
| XLOC_030395 | Platr26 | 2,046204022 | 0,0015 | 0,01775 |
| XLOC_040624 | Ldha-ps2 | 1,925259613 | 0,005 | 0,044365 |
| XLOC_020671 | Card10 | 1,827123307 | 0,00335 | 0,030266 |
| XLOC_054102 | Cemip | 1,82286788 | 0,00265 | 0,024191 |
| XLOC_032409 | Ptprj | 1,778874831 | 0,00265 | 0,024191 |
| XLOC_012282 | Ahnak2 | 1,763738896 | 0,0027 | 0,024623 |
| XLOC_007498 | Adam19 | 1,763103295 | 0,0005 | 0,01775 |
| XLOC_024886 | Notch3 | 1,694334825 | 0,00365 | 0,032772 |
| XLOC_041150 | Klhl17,Plekhn1 | 1,670464125 | 0,0044 | 0,039272 |
| XLOC_036591 | Kirrel | 1,651607643 | 0,001 | 0,01775 |
| XLOC_058850 | Ldlr | 1,566986771 | 0,0033 | 0,029864 |
| XLOC_035347 | Rps10-ps1 (retro) | -1,48949884 | 0,00375 | 0,033614 |
| XLOC_042975 | Gm5869 | -1,503349034 | 0,0043 | 0,038407 |
| XLOC_001865 | Gm7658 | -1,503737766 | 0,004 | 0,035803 |
| XLOC_006017 | Rpsa-ps2 | -1,506622521 | 0,0047 | 0,041762 |
| XLOC_052303 | RP23-136A14.1 | -1,508114536 | 0,00335 | 0,030266 |
| XLOC_024624 | Pabpc6 | -1,514249568 | 0,00515 | 0,045631 |
| XLOC_012147 | Gm6863 | -1,514338787 | 0,0045 | 0,040089 |

| | | | | |
|---|---|---|---|---|
| XLOC_035742 | Gm43524 | -1,517747714 | 0,00435 | 0,038835 |
| XLOC_059966 | Rpl15-ps3 | -1,51931709 | 0,0041 | 0,036681 |
| XLOC_040956 | Gm13050 | -1,520677264 | 0,0055 | 0,048561 |
| XLOC_029031 | Rpl9-ps6 | -1,523728643 | 0,00465 | 0,041337 |
| XLOC_063032 | Rps24-ps3 | -1,525971481 | 0,00355 | 0,031912 |
| XLOC_026674 | Gm10036 | -1,527824678 | 0,004 | 0,035803 |
| XLOC_002196 | Sumo1 | -1,528014252 | 0,0037 | 0,033189 |
| XLOC_057302 | Gm7901 | -1,530228396 | 0,00545 | 0,048153 |
| XLOC_044891 | Gm15484 | -1,531851025 | 0,00565 | 0,04985 |
| XLOC_001348 | Rpsa-ps1 | -1,533866581 | 0,0028 | 0,02548 |
| XLOC_056561 | 2310036O22Rik | -1,540718184 | 0,00525 | 0,046441 |
| XLOC_015341 | Gm10260 | -1,542733645 | 0,00325 | 0,029419 |
| XLOC_028250 | Fau | -1,54305662 | 0,0043 | 0,038407 |
| XLOC_057161 | Gm8623 | -1,544910219 | 0,0026 | 0,023769 |
| XLOC_045158 | Rps16-ps2 | -1,548252741 | 0,00325 | 0,029419 |
| XLOC_027132 | Gm10269 | -1,550020182 | 0,0052 | 0,046031 |
| XLOC_062956 | Rpl3-ps2 | -1,552164001 | 0,00195 | 0,018855 |
| XLOC_059156 | Rpp25 | -1,553327461 | 0,00205 | 0,01925 |
| XLOC_054828 | Ifitm2 | -1,55532923 | 0,0031 | 0,028108 |
| XLOC_026625 | Rpl27-ps3 | -1,560841454 | 0,0017 | 0,018436 |
| XLOC_020476 | Eif3h | -1,563210432 | 0,00205 | 0,01925 |
| XLOC_038868 | Gm11263 | -1,566829287 | 0,00275 | 0,025055 |
| XLOC_014531 | Rps23 | -1,569189949 | 0,0014 | 0,01775 |
| XLOC_058756 | Gm10709 | -1,575761291 | 0,0036 | 0,032338 |
| XLOC_037023 | Sars | -1,577929775 | 0,0045 | 0,040089 |
| XLOC_011096 | Gm4294 | -1,579168371 | 0,0017 | 0,018436 |
| XLOC_027343 | Rps2-ps10 | -1,583365147 | 0,00105 | 0,01775 |
| XLOC_034793 | Rps23-ps1 | -1,584780463 | 0,0032 | 0,02898 |
| XLOC_032949 | Rps15a-ps7 | -1,585269365 | 0,00185 | 0,01851 |
| XLOC_059387 | Rps27a-ps2 | -1,585957378 | 0,00225 | 0,020761 |
| XLOC_008657 | Uqcr10 | -1,593111232 | 0,00515 | 0,045631 |
| XLOC_002692 | Gm6170 | -1,595784679 | 0,0018 | 0,018436 |
| XLOC_002446 | Gm6136 | -1,595988217 | 0,0022 | 0,020359 |
| XLOC_053487 | Rps11 | -1,599411316 | 0,00095 | 0,01775 |
| XLOC_047229 | Rpl34-ps1 | -1,601615676 | 0,00085 | 0,01775 |
| XLOC_029179 | Rpl13a-ps1 | -1,602002057 | 0,0021 | 0,019685 |
| XLOC_004906 | Rps15 | -1,603276219 | 0,00115 | 0,01775 |
| XLOC_036830 | Rpl21-ps11 | -1,605803096 | 0,0042 | 0,03754 |
| XLOC_045388 | Atp5j2 | -1,605839827 | 0,0031 | 0,028108 |
| XLOC_048287 | Ybx1-ps2 | -1,606334112 | 0,0011 | 0,01775 |
| XLOC_018406 | Gm6055 | -1,607422298 | 0,00265 | 0,024191 |
| XLOC_043605 | Gm10051 | -1,60791707 | 0,00115 | 0,01775 |
| XLOC_017841 | Gm10233 | -1,611192642 | 0,0026 | 0,023769 |
| XLOC_017576 | Rps19-ps1 | -1,61660588 | 0,0051 | 0,04522 |
| XLOC_056848 | Rps23-ps2 | -1,621816955 | 0,0016 | 0,018404 |
| XLOC_060483 | Gm5619 | -1,622153113 | 0,00215 | 0,019965 |
| XLOC_008214 | Gm11539 | -1,624715372 | 0,00185 | 0,01851 |

| | | | | |
|---|---|---|---|---|
| XLOC_007801 | Rpl26 | -1,626567846 | 0,00095 | 0,01775 |
| XLOC_009544 | Gm11478 | -1,628568045 | 0,0036 | 0,032338 |
| XLOC_037363 | Rpsa-ps10 | -1,628650452 | 0,0031 | 0,028108 |
| XLOC_020029 | Rpl8 | -1,63132587 | 0,0052 | 0,046031 |
| XLOC_034678 | Gm7536 | -1,632151525 | 0,00185 | 0,01851 |
| XLOC_011266 | Gm5786 | -1,63233481 | 0,0006 | 0,01775 |
| XLOC_020387 | Basp1 | -1,632669753 | 0,0017 | 0,018436 |
| XLOC_012020 | Gm7862 | -1,632999105 | 0,00345 | 0,031132 |
| XLOC_006011 | Rps26 | -1,634771503 | 0,00105 | 0,01775 |
| XLOC_022313 | Sod1 | -1,635994616 | 0,00125 | 0,01775 |
| XLOC_059624 | Rpl29 | -1,639569434 | 0,0004 | 0,01775 |
| XLOC_014438 | Gm9625 | -1,63973764 | 0,00095 | 0,01775 |
| XLOC_052888 | RP24-490A22.9 | -1,641137366 | 0,00125 | 0,01775 |
| XLOC_026823 | Pabpc2 | -1,641747206 | 0,0055 | 0,048561 |
| XLOC_049488 | Rpl38-ps2 | -1,644376888 | 0,00055 | 0,01775 |
| XLOC_034941 | Rpl13-ps6 | -1,644925221 | 0,0012 | 0,01775 |
| XLOC_024814 | Gm8186 | -1,644985651 | 0,00125 | 0,01775 |
| XLOC_034705 | Rpl22l1 | -1,646551915 | 0,005 | 0,044365 |
| XLOC_039746 | Aurkaip1 | -1,648768674 | 0,0029 | 0,026371 |
| XLOC_019921 | Gm5045 | -1,649766675 | 0,00145 | 0,01775 |
| XLOC_012082 | Gm5436 | -1,650434633 | 0,0005 | 0,01775 |
| XLOC_043832 | Rpl21 | -1,651470272 | 0,0031 | 0,028108 |
| XLOC_051031 | Blvrb | -1,652420656 | 0,00445 | 0,039699 |
| XLOC_017541 | Rps19-ps2 | -1,652819293 | 0,00235 | 0,021609 |
| XLOC_011479 | Rpl30-ps8 | -1,653221464 | 0,00345 | 0,031132 |
| XLOC_036601 | Apoa1bp | -1,657145179 | 0,00535 | 0,047303 |
| XLOC_020478 | Gm10020 | -1,657786248 | 0,00065 | 0,01775 |
| XLOC_064183 | Gm14681 | -1,657936785 | 0,0007 | 0,01775 |
| XLOC_030034 | Rpsa-ps9 | -1,659934138 | 0,0006 | 0,01775 |
| XLOC_064838 | Gm6472,Rpl7a-ps12 | -1,661159956 | 0,0022 | 0,020359 |
| XLOC_011965 | Rpl36al | -1,661439777 | 0,00065 | 0,01775 |
| XLOC_003139 | Gm10171 | -1,662952545 | 0,00295 | 0,026812 |
| XLOC_039448 | Id3 | -1,664053711 | 0,00475 | 0,042196 |
| XLOC_049113 | Rpl32 | -1,667791569 | 0,0012 | 0,01775 |
| XLOC_024835 | Rps10 | -1,66859173 | 0,001 | 0,01775 |
| XLOC_050844 | Zfp296 | -1,669071779 | 0,00455 | 0,040505 |
| XLOC_060211 | Rps6-ps3 | -1,670290453 | 0,00125 | 0,01775 |
| XLOC_046718 | Gm13864 | -1,676160151 | 0,0042 | 0,03754 |
| XLOC_030100 | Gm13611 | -1,676640054 | 0,0008 | 0,01775 |
| XLOC_063314 | Gm16373 | -1,677318892 | 0,00315 | 0,028548 |
| XLOC_039969 | Bag1 | -1,677944503 | 0,0017 | 0,018436 |
| XLOC_048567 | Gm9794 | -1,679533998 | 0,0014 | 0,01775 |
| XLOC_001822 | Rpl7 | -1,681298634 | 0,0007 | 0,01775 |
| XLOC_039148 | Gm12848 | -1,68456839 | 0,0025 | 0,02291 |
| XLOC_062660 | Gm6274 | -1,689373119 | 0,00025 | 0,01775 |
| XLOC_063984 | Rpl39 | -1,694520394 | 0,0001 | 0,009173 |
| XLOC_047051 | Rps15-ps2 | -1,697429905 | 0,00045 | 0,01775 |

| | | | | |
|---|---|---|---|---|
| XLOC_018725 | Gm16409 | -1,699155632 | 0,00165 | 0,018404 |
| XLOC_051433 | Rpl14-ps1 | -1,705344251 | 0,0005 | 0,01775 |
| XLOC_001308 | Mrps14 | -1,707267356 | 0,00085 | 0,01775 |
| XLOC_058936 | Gm10177 | -1,711874572 | 0,0023 | 0,021206 |
| XLOC_036359 | Gm6394 | -1,71311856 | 0,0004 | 0,01775 |
| XLOC_038398 | Gm11824 | -1,713506898 | 0,0003 | 0,01775 |
| XLOC_025265 | Rpl7a-ps5 | -1,717807053 | 0,0011 | 0,01775 |
| XLOC_014150 | Gm11353 | -1,724626556 | 0,0003 | 0,01775 |
| XLOC_064384 | Gm6977 | -1,726542682 | 0,00025 | 0,01775 |
| XLOC_005610 | Gm6419 | -1,727095669 | 0,00235 | 0,021609 |
| XLOC_006007 | Rpl41 | -1,728000937 | 0,00025 | 0,01775 |
| XLOC_009086 | Gm12254 | -1,728741311 | 0,0001 | 0,009173 |
| XLOC_020726 | Ndufa6 | -1,729790117 | 0,00335 | 0,030266 |
| XLOC_059794 | Rpsa | -1,730470083 | 0,00065 | 0,01775 |
| XLOC_027217 | Pfdn1 | -1,731621958 | 0,0001 | 0,009173 |
| XLOC_008261 | Rpl19 | -1,732641286 | 0,0001 | 0,009173 |
| XLOC_040696 | Rpl28-ps3 | -1,733267107 | 0,0007 | 0,01775 |
| XLOC_049378 | Ybx3 | -1,736239537 | 0,0036 | 0,032338 |
| XLOC_022853 | Gm9843 | -1,740541243 | 0,0003 | 0,01775 |
| XLOC_059805 | Rpl14 | -1,74333642 | 0,0001 | 0,009173 |
| XLOC_038363 | Gm12918 | -1,744440028 | 0,001 | 0,01775 |
| XLOC_007395 | Rpsa-ps5 | -1,747751411 | 0,0003 | 0,01775 |
| XLOC_048715 | Mrpl35 | -1,747954947 | 0,00145 | 0,01775 |
| XLOC_040275 | Gm10154 | -1,752718127 | 0,00055 | 0,01775 |
| XLOC_033201 | Gm14303 | -1,756226596 | 0,0012 | 0,01775 |
| XLOC_051528 | Rps12l1 | -1,757822014 | 0,0006 | 0,01775 |
| XLOC_032132 | Mrpl23-ps1 | -1,757962139 | 0,0022 | 0,020359 |
| XLOC_064783 | Gm15066 | -1,759574989 | 5,00E-05 | 0,004846 |
| XLOC_057536 | Rps18-ps3 | -1,764439545 | 5,00E-05 | 0,004846 |
| XLOC_051915 | Ndufc2 | -1,766826039 | 0,004 | 0,035803 |
| XLOC_017967 | Gm10076 | -1,7705321 | 5,00E-05 | 0,004846 |
| XLOC_001278 | Gm2000 | -1,770794748 | 0,0001 | 0,009173 |
| XLOC_001880 | Tubb4b-ps2 | -1,775614014 | 0,00245 | 0,022479 |
| XLOC_058788 | Gm5611 | -1,777027491 | 0,0001 | 0,009173 |
| XLOC_049450 | Dynlt1-ps1 | -1,78264694 | 0,00015 | 0,013338 |
| XLOC_009054 | Anxa6 | -1,782735908 | 0,00555 | 0,048991 |
| XLOC_034781 | Gm12583 | -1,783829837 | 0,00015 | 0,013338 |
| XLOC_063677 | Gm7331 | -1,784596604 | 5,00E-05 | 0,004846 |
| XLOC_031032 | Gm14036 | -1,788275483 | 0,0013 | 0,01775 |
| XLOC_042648 | Gm4754 | -1,793767616 | 0,00095 | 0,01775 |
| XLOC_041051 | Rpsa-ps12 | -1,804441035 | 0,00065 | 0,01775 |
| XLOC_014482 | Gm4149 | -1,811866927 | 5,00E-05 | 0,004846 |
| XLOC_018771 | Rps3a2 | -1,819617229 | 0,00025 | 0,01775 |
| XLOC_060367 | Gm7866 | -1,825513073 | 0,0001 | 0,009173 |
| XLOC_026932 | Rps14 | -1,825937014 | 0,00015 | 0,013338 |
| XLOC_052806 | Rps8-ps4 | -1,828742566 | 0,00375 | 0,033614 |
| XLOC_064277 | Gm15361 | -1,829777211 | 0,00045 | 0,01775 |

| | | | | |
|---|---|---|---|---|
| XLOC_004908 | Reep6 | -1,833519996 | 0,0042 | 0,03754 |
| XLOC_011651 | Rps19-ps6 | -1,834308123 | 0,00015 | 0,013338 |
| XLOC_064100 | Gm14586 | -1,840293661 | 5,00E-05 | 0,004846 |
| XLOC_020011 | Rps6-ps1 | -1,844663926 | 0,0008 | 0,01775 |
| XLOC_054197 | Gm15501 | -1,847016814 | 0,0008 | 0,01775 |
| XLOC_054653 | Nupr1 | -1,848218084 | 0,00125 | 0,01775 |
| XLOC_004508 | Gm10335 | -1,854394253 | 0,00065 | 0,01775 |
| XLOC_017784 | Rpl13-ps3 | -1,855514146 | 0,0002 | 0,0177 |
| XLOC_018023 | Gm2904 | -1,858290435 | 0,001 | 0,01775 |
| XLOC_057234 | Ndufa13 | -1,861957315 | 5,00E-05 | 0,004846 |
| XLOC_051455 | Rpl18 | -1,862864836 | 5,00E-05 | 0,004846 |
| XLOC_011829 | Rps7 | -1,863754712 | 0,0005 | 0,01775 |
| XLOC_057539 | Rps26-ps1 | -1,868718162 | 0,00335 | 0,030266 |
| XLOC_055991 | RP24-74L7.3 | -1,870146125 | 0,0005 | 0,01775 |
| XLOC_054283 | Gm7027 | -1,873944207 | 5,00E-05 | 0,004846 |
| XLOC_044441 | Rpl9 | -1,876663517 | 0,00465 | 0,041337 |
| XLOC_062914 | Rpl30-ps10 | -1,879788045 | 5,00E-05 | 0,004846 |
| XLOC_000026 | Gm37108 | -1,880536098 | 5,00E-05 | 0,004846 |
| XLOC_063168 | Tpt1-ps6 | -1,889017412 | 5,00E-05 | 0,004846 |
| XLOC_014912 | Acot13 | -1,889821535 | 0,00225 | 0,020761 |
| XLOC_001009 | Gm8392 | -1,895806868 | 5,00E-05 | 0,004846 |
| XLOC_012038 | Vti1b | -1,896674355 | 0,0039 | 0,034933 |
| XLOC_020802 | Rpl31-ps8 | -1,905421272 | 5,00E-05 | 0,004846 |
| XLOC_005520 | Rps8-ps1 | -1,913776029 | 0,0001 | 0,009173 |
| XLOC_014341 | Gm6344 | -1,918830132 | 0,0009 | 0,01775 |
| XLOC_005716 | Timm13 | -1,925052778 | 0,00065 | 0,01775 |
| XLOC_033240 | Rps8-ps2 | -1,929375565 | 5,00E-05 | 0,004846 |
| XLOC_001121 | Gm15454 | -1,931195206 | 0,0016 | 0,018404 |
| XLOC_031489 | Gm14414 | -1,932382912 | 5,00E-05 | 0,004846 |
| XLOC_040228 | Rps18-ps1 | -1,93616647 | 0,0037 | 0,033189 |
| XLOC_039776 | Gm11808 | -1,937268606 | 5,00E-05 | 0,004846 |
| XLOC_020052 | Nol12 | -1,945528346 | 0,00265 | 0,024191 |
| XLOC_017160 | Gm17048 | -1,946306607 | 0,0028 | 0,02548 |
| XLOC_049011 | Gm6681 | -1,946461757 | 0,00285 | 0,025922 |
| XLOC_036687 | Gm5851 | -1,94976331 | 5,00E-05 | 0,004846 |
| XLOC_018745 | Gm10132 | -1,955530449 | 0,0006 | 0,01775 |
| XLOC_036685 | Gm5850 | -1,958574491 | 0,00015 | 0,013338 |
| XLOC_064525 | Rpl30-ps9 | -1,962501831 | 5,00E-05 | 0,004846 |
| XLOC_025344 | Gm6548 | -1,97366892 | 5,00E-05 | 0,004846 |
| XLOC_009518 | Aatf | -1,978864734 | 0,0009 | 0,01775 |
| XLOC_031462 | Gm14438 | -1,979799045 | 5,00E-05 | 0,004846 |
| XLOC_044990 | Gm13841 | -1,985659154 | 5,00E-05 | 0,004846 |
| XLOC_039204 | Gm12857 | -1,98598813 | 5,00E-05 | 0,004846 |
| XLOC_039772 | Rps20 | -1,99189003 | 5,00E-05 | 0,004846 |
| XLOC_059912 | Gm7808 | -2,003302104 | 5,00E-05 | 0,004846 |
| XLOC_036999 | Gm5075 | -2,009198356 | 0,0032 | 0,02898 |
| XLOC_017158 | Gm17046 | -2,012724903 | 0,003 | 0,027254 |

| | | | | |
|---|---|---|---|---|
| XLOC_036435 | Gm37009 | -2,016229689 | 5,00E-05 | 0,004846 |
| XLOC_019775 | Capsl | -2,035721211 | 0,00335 | 0,030266 |
| XLOC_031950 | Rpl35 | -2,041840255 | 0,00075 | 0,01775 |
| XLOC_037415 | Gm6520 | -2,043652637 | 0,0034 | 0,030703 |
| XLOC_018394 | Bmp4 | -2,049014225 | 0,00095 | 0,01775 |
| XLOC_000314 | Gm8210 | -2,057410966 | 5,00E-05 | 0,004846 |
| XLOC_045008 | Gm3786 | -2,058466543 | 0,0007 | 0,01775 |
| XLOC_003119 | Rpl27-ps1 | -2,059365636 | 0,0001 | 0,009173 |
| XLOC_033083 | Gtsf1l | -2,064181743 | 5,00E-05 | 0,004846 |
| XLOC_008978 | Gm12191 | -2,065555751 | 0,0055 | 0,048561 |
| XLOC_043683 | Gm16089 | -2,067833463 | 0,001 | 0,01775 |
| XLOC_064304 | Gm8692 | -2,074120913 | 0,00165 | 0,018404 |
| XLOC_030834 | Gm14016 | -2,077963049 | 0,00125 | 0,01775 |
| XLOC_018749 | Dnajc15 | -2,086608587 | 0,0045 | 0,040089 |
| XLOC_020899 | Bin2 | -2,090502835 | 0,00505 | 0,044787 |
| XLOC_058944 | Tpt1-ps5 | -2,094607604 | 0,0052 | 0,046031 |
| XLOC_026983 | Gm17669 | -2,101471659 | 5,00E-05 | 0,004846 |
| XLOC_020438 | Gm3362 | -2,115443274 | 5,00E-05 | 0,004846 |
| XLOC_054352 | Dnajc19-ps | -2,118421986 | 0,00035 | 0,01775 |
| XLOC_053489 | Rpl13a | -2,130762802 | 5,00E-05 | 0,004846 |
| XLOC_030136 | Gm13408 | -2,141260024 | 5,00E-05 | 0,004846 |
| XLOC_036764 | Rpl31-ps11 | -2,146371792 | 5,00E-05 | 0,004846 |
| XLOC_003368 | Gm19777 | -2,153509966 | 5,00E-05 | 0,004846 |
| XLOC_050978 | Rps19 | -2,156153665 | 0,0007 | 0,01775 |
| XLOC_032778 | Gm14044 | -2,159354337 | 0,00015 | 0,013338 |
| XLOC_035723 | Rpl7a-ps7 | -2,176168095 | 0,00035 | 0,01775 |
| XLOC_009165 | Rps13-ps5 | -2,178039321 | 5,00E-05 | 0,004846 |
| XLOC_052242 | E130201H02Rik | -2,182452115 | 0,00025 | 0,01775 |
| XLOC_036577 | Rps3a1 | -2,184162204 | 5,00E-05 | 0,004846 |
| XLOC_017150 | Gm3348 | -2,184434731 | 0,0037 | 0,033189 |
| XLOC_028431 | Prune2 | -2,184798154 | 0,0006 | 0,01775 |
| XLOC_020933 | Itgb7 | -2,193690507 | 0,00565 | 0,04985 |
| XLOC_014603 | Rpl31-ps13 | -2,195668113 | 5,00E-05 | 0,004846 |
| XLOC_063124 | Rpl17-ps8 | -2,205292468 | 5,00E-05 | 0,004846 |
| XLOC_024543 | Rpl31-ps16 | -2,224575403 | 0,00165 | 0,018404 |
| XLOC_017339 | Rpl23a-ps3 | -2,225392791 | 5,00E-05 | 0,004846 |
| XLOC_022879 | Rpl31-ps4 | -2,239970916 | 0,00195 | 0,018855 |
| XLOC_053112 | Gm4613 | -2,269209801 | 0,0022 | 0,020359 |
| XLOC_053968 | Rpl17-ps10 | -2,284423293 | 5,00E-05 | 0,004846 |
| XLOC_022482 | Rpl31-ps12 | -2,285120113 | 5,00E-05 | 0,004846 |
| XLOC_020887 | Gm9763 | -2,299308426 | 0,00155 | 0,01812 |
| XLOC_035524 | Gm4540 | -2,301508866 | 5,00E-05 | 0,004846 |
| XLOC_051250 | Rpl17-ps9 | -2,309691232 | 5,00E-05 | 0,004846 |
| XLOC_054162 | Rps13-ps2 | -2,343703391 | 5,00E-05 | 0,004846 |
| XLOC_048012 | Rps25-ps1 | -2,358010267 | 5,00E-05 | 0,004846 |
| XLOC_060808 | Rps27rt | -2,358369873 | 5,00E-05 | 0,004846 |
| XLOC_050570 | Rps9 | -2,3690358 | 5,00E-05 | 0,004846 |

| | | | | |
|---|---|---|---|---|
| XLOC_011642 | Dio3 | -2,375712077 | 0,00245 | 0,022479 |
| XLOC_054218 | Aamdc | -2,377853775 | 0,0034 | 0,030703 |
| XLOC_008572 | Ccdc40 | -2,390562293 | 0,0037 | 0,033189 |
| XLOC_002271 | Rpl31-ps14 | -2,392518367 | 5,00E-05 | 0,004846 |
| XLOC_009436 | Rpl23a | -2,398279828 | 5,00E-05 | 0,004846 |
| XLOC_002678 | Gm8451 | -2,41119779 | 0,00025 | 0,01775 |
| XLOC_002845 | Gm7266 | -2,413940305 | 5,00E-05 | 0,004846 |
| XLOC_051460 | Ccdc114 | -2,414425587 | 0,00075 | 0,01775 |
| XLOC_014037 | Hist1h2ai | -2,463774128 | 0,0024 | 0,022042 |
| XLOC_014126 | Hist1h2ab | -2,466696126 | 0,0011 | 0,01775 |
| XLOC_064432 | Zc4h2 | -2,486092542 | 0,00135 | 0,01775 |
| XLOC_038702 | Gm12482 | -2,487230133 | 5,00E-05 | 0,004846 |
| XLOC_030898 | Rps12-ps10 | -2,509152227 | 0,00135 | 0,01775 |
| XLOC_008026 | Gm11204 | -2,548220287 | 0,00275 | 0,025055 |
| XLOC_023997 | Gm8225 | -2,592924143 | 5,00E-05 | 0,004846 |
| XLOC_056590 | Rps13-ps1 | -2,61204545 | 5,00E-05 | 0,004846 |
| XLOC_053209 | RP24-471H15.4 | -2,648581679 | 5,00E-05 | 0,004846 |
| XLOC_014824 | Hist1h2ah | -2,697849744 | 0,00485 | 0,043074 |
| XLOC_017272 | Gm7565 | -2,880341757 | 5,00E-05 | 0,004846 |
| XLOC_014878 | Hist1h2ae | -3,058533335 | 0,00165 | 0,018404 |
| XLOC_057615 | Rps13-ps4 | -3,099939372 | 0,0023 | 0,021206 |
| XLOC_019856 | Rps19-ps5 | -3,147573353 | 5,00E-05 | 0,004846 |
| XLOC_014884 | Hist1h2ac | -3,233955888 | 0,001 | 0,01775 |
| XLOC_022752 | Dubr | -3,316450516 | 0,0054 | 0,047734 |
| XLOC_014826 | Hist1h2ag | -3,338823893 | 0,0003 | 0,01775 |
| XLOC_001415 | Gm7299 | -3,363166024 | 0,0012 | 0,01775 |
| XLOC_036291 | Ndufc1 | -3,459203769 | 0,0053 | 0,046872 |
| XLOC_027008 | Myo5b | -3,526336133 | 0,00095 | 0,01775 |
| XLOC_014115 | Hist1h2ad | -3,830014887 | 5,00E-05 | 0,004846 |
| XLOC_031601 | 9230102O04Rik,Gata3 | -3,894019207 | 0,0004 | 0,01775 |
| XLOC_014815 | Hist1h2ap | -4,043262863 | 5,00E-05 | 0,004846 |
| XLOC_056849 | Xab2 | -4,173882536 | 5,00E-05 | 0,004846 |
| XLOC_002874 | Pla2g4a | -4,211838155 | 0,0013 | 0,01775 |
| XLOC_014047 | Hist1h2ao | -4,229684341 | 5,00E-05 | 0,004846 |
| XLOC_056788 | Vat1l | -4,31230733 | 0,00265 | 0,024191 |
| XLOC_040846 | Man1c1 | -4,454983227 | 0,0001 | 0,009173 |
| XLOC_038474 | Gm11918 | -4,531127227 | 0,00055 | 0,01775 |
| XLOC_001830 | Jph1 | -5,117999477 | 0,00505 | 0,044787 |
| XLOC_006071 | Gm20746,Gm26445 | -5,781925421 | 0,0001 | 0,009173 |
| XLOC_064058 | Smarca1 | -7,355584564 | 0,00015 | 0,013338 |
| XLOC_014340 | Unc5a | -8,241060486 | 0,0031 | 0,028108 |
| XLOC_014498 | A830082K12Rik | -9,031026631 | 0,0024 | 0,022042 |
| XLOC_063433 | Tceal3 | -9,597375898 | 0,00105 | 0,01775 |
| XLOC_036993 | Slc6a17 | -11,07212323 | 0,0001 | 0,009173 |
| XLOC_052247 | Vwa3a | -12,10918614 | 0,00145 | 0,01775 |
| XLOC_031015 | AU019990 | -12,41532394 | 0,0014 | 0,01775 |
| XLOC_059042 | C1qtnf5,Mfrp | -28,79207779 | 5,00E-05 | 0,004846 |

| | | | | |
|---|---|---|---|---|
| XLOC_042512 | Kcnk3 | -30,82715829 | 0,0047 | 0,041762 |
| XLOC_009585 | Hlf | -31,60235135 | 0,0018 | 0,018436 |
| XLOC_018791 | Gm25130 | ND in si*Xab2* cells | 0,0013 | 0,01775 |
| XLOC_052323 | RP23-421P23.5 | ND in si*Xab2* cells | 0,0013 | 0,01775 |
| XLOC_004025 | | ND in si*Xab2* cells | 0,00105 | 0,01775 |
| XLOC_060788 | Rp31-ps19 | ND in si*Xab2* cells | 0,001 | 0,01775 |
| XLOC_053450 | RP23-361A18.17 | ND in si*Xab2* cells | 5,00E-05 | 0,004846 |
