## Supplemental table 5 for "The Splicing Factor XAB2 interacts with ERCC1-XPF and XPG for RNA-loop processing during mammalian development"

### Table S5. *Xab2* -induced differential splicing events in HEPA cells

| GENE | EVENT | COORD | Complex | β_siRNA | β_Ctrl | β_siRNA_Ctrl | P_xzero |
|---|---|---|---|---|---|---|---|
| Atp6v1a | MmuALTD0001887-1/3 | chr16:44139075-44139093 | Alt5 | 0,1090087 | 0,7996605 | -0,690651737 | 0,922 |
| Cyp4f16 | MmuALTA0004959-1/2 | chr17:32673943-32674141 | Alt3 | 0,35937 | 0,9639825 | -0,604612448 | 0,838 |
| Ncor1 | MmuEX0031185 | chr11:62147165-62147658 | C1 | 0,5356821 | 0,9895236 | -0,453841488 | 0,8233333 |
| Atf2 | MmuEX0006584 | chr2:73684254-73684432 | C2 | 0,6880473 | 0,9750753 | -0,287027975 | 0,8053333 |
| Serpinb6a | MmuALTD0012645-1/5 | chr13:34027636-34027663 | Alt5 | 0,0785913 | 0,3655519 | -0,28696058 | 0,9293333 |
| Mr1 | MmuALTD0008714-1/2 | chr1:156984829-156984904 | Alt5 | 0,0804305 | 0,3341185 | -0,253687984 | 0,9653333 |
| Tmem161a | MmuEX0047784 | chr8:72698202-72698305 | S | 0,5361885 | 0,7826644 | -0,24647598 | 0,8593333 |
| Git2 | MmuALTA0007752-1/2 | chr5:115202288-115202336 | Alt3 | 0,4584876 | 0,6987804 | -0,240292837 | 0,804 |
| Fmr1 | MmuALTA0007321-1/2 | chrX:65965442-65965480 | Alt3 | 0,1628548 | 0,395801 | -0,232946172 | 0,91 |
| Mier2 | MmuALTA0010968-1/2 | chr10:79012381-79012424 | Alt3 | 0,6986348 | 0,917274 | -0,218639214 | 0,8653333 |
| Nfya | MmuALTD0009355-1/2 | chr17:48538310-48538363 | Alt5 | 0,1562663 | 0,3699905 | -0,213724207 | 0,9033333 |
| Rbm14 | MmuALTD0011633-1/2 | chr19:4793816-4793849 | Alt5 | 0,565482 | 0,7791952 | -0,213713145 | 0,8313333 |
| Alkbh7 | MmuEX0004696 | chr17:57137812-57137985 | S | 0,7540418 | 0,9539332 | -0,199891474 | 0,856 |
| D19Wsu162e | MmuEX0013522 | chr19:46681202-46681249 | S | 0,6695706 | 0,8598263 | -0,190255669 | 0,942 |
| Arhgef12 | MmuALTA0001960-1/2 | chr9:42816099-42816191 | Alt3 | 0,3999025 | 0,5895391 | -0,189636567 | 0,8406667 |
| Mtif3 | MmuALTD0008893-1/3 | chr5:147775299-147775319 | Alt5 | 0,4193444 | 0,6079851 | -0,188640649 | 0,8473333 |
| Fanca | MmuINT0062263 | chr8:125792738-125792842 | IR-C | 0,3356849 | 0,5219056 | -0,186220661 | 0,9126667 |
| Tpcn2 | MmuEX0048557 | chr7:152441356-152441438 | S | 0,7927872 | 0,9789566 | -0,186169376 | 0,8446667 |
| Tmem38b | MmuEX0047954 | chr4:53853499-53853655 | S | 0,7359678 | 0,9215461 | -0,185578221 | 0,9393333 |
| Zfp740 | MmuALTA0020740-1/2 | chr15:102038703-102038773 | Alt3 | 0,3971194 | 0,5807793 | -0,18365998 | 0,9173333 |
| Ubr4 | MmuEX0050365 | chr4:138997107-138997211 | S | 0,6823619 | 0,8633006 | -0,180938727 | 0,88 |
| BC017643 | MmuALTD0002029-1/2 | chr11:121090373-121090409 | Alt5 | 0,2559084 | 0,4348297 | -0,178921364 | 0,8013333 |
| Cdk12 | MmuALTA0003745-1/2 | chr11:98111024-98111414 | Alt3 | 0,6222564 | 0,8000865 | -0,177830112 | 0,862 |
| Ankzf1 | MmuEX0005222 | chr1:75190753-75190859 | S | 0,8046241 | 0,9769959 | -0,172371803 | 0,98 |
| Atp6v1e1 | MmuALTD0001895-1/3 | chr6:120772610- | Alt5 | 0,1445453 | 0,3162145 | -0,171669252 | 0,96 |
| Actr5 | MmuALTD0000932-1/2 | chr2:158457365-158457416 | Alt5 | 0,7795977 | 0,9511359 | -0,171538247 | 0,892 |
| Eed | MmuALTD0004652-1/2 | chr7:97104854-97104923 | Alt5 | 0,1313608 | 0,3024625 | -0,171101681 | 0,926 |
| Smek1 | MmuALTD0013244-1/2 | chr12:102291678-102291794 | Alt5 | 0,6204392 | 0,7910117 | -0,170572583 | 0,8033333 |
| 1110057K04Rik | MmuALTD0000068-1/3 | chr12:-8214998 | Alt5 | 0,2262214 | 0,3938678 | -0,1676464 | 0,808 |
| Clasrp | MmuALTD0003224-1/2 | chr7:20170133-20170172 | Alt5 | 0,6606028 | 0,8255283 | -0,164925491 | 0,8286667 |
| Tubgcp2 | MmuALTD0015006-1/5 | chr7:147222246- | Alt5 | 0,2273246 | 0,3920411 | -0,164716461 | 0,8826667 |
| Trrap | MmuALTA0018970-1/2 | chr5:145552143-145552237 | Alt3 | 0,2511486 | 0,4152459 | -0,164097284 | 0,858 |
| Bmp4 | MmuINT0025192 | chr14:47005770-47006830 | IR-S | 0,072019 | 0,2340737 | -0,16205473 | 0,958 |
| Fam48a | MmuALTA0006920-1/3 | chr3:54526479-54526562 | Alt3 | 0,1799672 | 0,3417749 | -0,161807718 | 0,8166667 |
| Acot8 | MmuEX0003517 | chr2:164628497-164628633 | S | 0,6458656 | 0,8053368 | -0,159471221 | 0,8526667 |
| Ddx6 | MmuALTD0004130-1/6 | chr9:-44413010 | Alt5 | 0,2108359 | 0,3700148 | -0,159178893 | 0,8586667 |
| Mef2d | MmuEX0028565 | chr3:87971960-87972266 | S | 0,6627814 | 0,8216785 | -0,158897152 | 0,858 |
| Mef2a | MmuEX0028561 | chr7:74412953-74413090 | C3 | 0,3623816 | 0,5202061 | -0,15782453 | 0,8893333 |
| Uqcc | MmuEX0050723 | chr2:155727034-155727091 | C3 | 0,8071934 | 0,9633238 | -0,156130393 | 0,8566667 |
| Senp6 | MmuALTA0016126-1/2 | chr9:79946700-79946713 | Alt3 | 0,0288114 | 0,1836152 | -0,154803855 | 0,8113333 |
| Ift52 | MmuALTD0006914-1/2 | chr2:-162843269 | Alt5 | 0,7147012 | 0,867003 | -0,152301798 | 0,8493333 |
| Dlg4 | MmuEX0014814 | chr11:69857160-69857269 | S | 0,4193698 | 0,5711075 | -0,15173777 | 0,87 |
| Atf2 | MmuEX0006585 | chr2:73683496-73683610 | C1 | 0,6891787 | 0,8379289 | -0,148750189 | 0,808 |
| Cep55 | MmuALTD0003039-1/2 | chr19:-38129571 | Alt5 | 0,5879371 | 0,7358439 | -0,147906807 | 0,8133333 |
| Mtpap | MmuALTA0011446-1/2 | chr18:4383187-4383411 | Alt3 | 0,7592401 | 0,9056124 | -0,146372269 | 0,8386667 |
| Thap7 | MmuEX0047170 | chr16:17530323-17530478 | C2 | 0,5922371 | 0,738304 | -0,146066897 | 0,866 |
| Banp | MmuEX0007671 | chr8:124513855-124513971 | S | 0,6597408 | 0,8038098 | -0,144069008 | 0,856 |
| Setd5 | MmuEX0041847 | chr6:113055911-113056135 | C1 | 0,550027 | 0,6936253 | -0,143598346 | 0,902 |
| Nr2f2 | MmuALTD0009542-1/2 | chr7:77504774-77505479 | Alt5 | 0,730448 | 0,8720211 | -0,141573183 | 0,8713333 |
| Pcyt2 | MmuEX0034099 | chr11:120474344-120474397 | S | 0,4387014 | 0,5794984 | -0,14079701 | 0,904 |
| Odf2 | MmuALTD0009758-1/2 | chr2:29748964-29749032 | Alt5 | 0,4045722 | 0,5439274 | -0,139355163 | 0,876 |
| Fam53a | MmuEX0018352 | chr5:33971502-33971626 | S | 0,8441551 | 0,9826355 | -0,138480319 | 0,9733333 |
| Acbd5 | MmuEX0003421 | chr2:22931010-22931130 | S | 0,8325246 | 0,9709464 | -0,138421792 | 0,828 |
| Senp6 | MmuEX0041679 | chr9:79946693-79946713 | S | 0,6422916 | 0,7805176 | -0,13822596 | 0,8866667 |
| Prpf40b | MmuEX0037345 | chr15:99135597-99135624 | S | 0,7811273 | 0,918409 | -0,137281671 | 0,8426667 |
| Hgfac | MmuEX0022853 | chr5:35389138-35389277 | S | 0,7169644 | 0,8533042 | -0,136339872 | 0,828 |
| 1810031K17Rik | MmuINT0001713 | chr1:75138078-75138631 | IR-C | 0,4049344 | 0,5403648 | -0,135430444 | 0,874 |
| Hsd17b7 | MmuEX0023306 | chr1:171889554-171889610 | S | 0,8118881 | 0,9469799 | -0,135091849 | 0,952 |
| Stim1 | MmuEX0045370 | chr7:109577288-109577380 | S | 0,3353827 | 0,4696332 | -0,134250429 | 0,818 |
| Rif1 | MmuEX0039819 | chr2:51972188-51972265 | S | 0,3597079 | 0,4935274 | -0,133819493 | 0,8393333 |

| | | | | | | | |
|---|---|---|---|---|---|---|---|
| Pus10 | MmuALTA0014590-1/3 | chr11:23628975-23629071 | Alt3 | 0,7909149 | 0,92387 | -0,132955151 | 0,8126667 |
| D930015E06Rik | MmuALTA0005042-1/3 | chr3:83704249-83704337 | Alt3 | 0,106469 | 0,2387084 | -0,132239347 | 0,8426667 |
| Ubr3 | MmuALTA0019357-1/2 | chr2:69789404-69789465 | Alt3 | 0,1049374 | 0,2357565 | -0,130819101 | 0,8433333 |
| Fbxw5 | MmuEX0018961 | chr2:25357921-25358078 | S | 0,6564712 | 0,7871617 | -0,130690505 | 0,8246667 |
| Cacnb3 | MmuALTA0003082-1/2 | chr15:98471878-98471889 | Alt3 | 0,0162489 | 0,1462584 | -0,130009467 | 0,8013333 |
| Ttc19 | MmuEX0049456 | chr11:62097633-62097671 | S | 0,7870955 | 0,9164614 | -0,129365881 | 0,896 |
| Rpl23 | MmuALTA0015503-1/2 | chr11:97642718-97642744 | Alt3 | 0,6148007 | 0,7436096 | -0,128808889 | 0,8606667 |
| Wbp1 | MmuEX0051721 | chr6:83070766-83070867 | S | 0,2637117 | 0,3921128 | -0,128401096 | 0,928 |
| Kbtbd4 | MmuALTD0007324-1/2 | chr2:-90744969 | Alt5 | 0,0728694 | 0,200331 | -0,12746162 | 0,8393333 |
| Luc7l | MmuEX0027311 | chr17:26390919-26390989 | S | 0,2907271 | 0,4173451 | -0,126618061 | 0,8953333 |
| Cln6 | MmuEX0011748 | chr9:62688292-62688398 | S | 0,0896504 | 0,2160062 | -0,12635584 | 0,8246667 |
| Tmem161a | MmuEX0047786 | chr8:72701140-72701237 | C3 | 0,661919 | 0,7879595 | -0,126040528 | 0,8573333 |
| Pan3 | MmuALTA0012718-1/2 | chr5:148348038-148348116 | Alt3 | 0,0720857 | 0,1974971 | -0,125411449 | 0,8013333 |
| Plekha6 | MmuEX0035729 | chr1:135195497-135195620 | C1 | 0,8082256 | 0,9323877 | -0,124162094 | 0,916 |
| Tsc2 | MmuEX0049266 | chr17:24743169-24743297 | S | 0,5000818 | 0,6241061 | -0,12402428 | 0,8026667 |
| Sept6 | MmuALTD0012613-1/2 | chrX:34482077-34482211 | Alt5 | 0,0800657 | 0,2040751 | -0,124009373 | 0,846 |
| Dsn1 | MmuALTD0004496-1/2 | chr2:156832711- | Alt5 | 0,7990031 | 0,9224673 | -0,123464172 | 0,802 |
| Abcc1 | MmuEX0003090 | chr16:14413314-14413475 | S | 0,8501773 | 0,9732794 | -0,123102125 | 0,9286667 |
| Rap1gap | MmuEX0038750 | chr4:137279661-137279738 | S | 0,1544303 | 0,2772729 | -0,122842639 | 0,8893333 |
| Senp1 | MmuEX0041653 | chr15:97892381-97892458 | S | 0,2325932 | 0,3539959 | -0,12140271 | 0,8526667 |
| Zfp568 | MmuINT0179471 | chr7:30774127-30782768 | IR-S | 0,1157406 | 0,2365569 | -0,120816282 | 0,846 |
| Sbf1 | MmuEX0041054 | chr15:89126383-89126460 | S | 0,518793 | 0,6389842 | -0,120191155 | 0,8713333 |
| Tmem131 | MmuALTA0018335-1/2 | chr1:36850889-36851040 | Alt3 | 0,201343 | 0,3214679 | -0,120124887 | 0,8066667 |
| Palm | MmuEX0033522 | chr10:79279537-79279668 | S | 0,4573629 | 0,5774638 | -0,120100887 | 0,8353333 |
| Stk25 | MmuEX0045410 | chr1:95521611-95521725 | S | 0,7834155 | 0,9030233 | -0,119607809 | 0,8266667 |
| Nol11 | MmuALTA0012107-1/3 | chr11:107034524-107034715 | Alt3 | 0,3342629 | 0,4522152 | -0,11795233 | 0,8193333 |
| Myo9b | MmuALTA0011588-1/2 | chr8:73883563- | Alt3 | 0,0964731 | 0,2120939 | -0,115620823 | 0,824 |
| Rock2 | MmuEX0040276 | chr12:16965434-16965680 | S | 0,8760814 | 0,9916898 | -0,115608465 | 0,824 |
| Cnot1 | MmuALTA0004327-1/2 | chr8:98278885-98279008 | Alt3 | 0,2163931 | 0,3317658 | -0,115372705 | 0,8066667 |
| 0610037P05Rik | MmuEX0000067 | chr16:14311140-14311224 | C2 | 0,7928753 | 0,9060622 | -0,113186937 | 0,9006667 |
| Ccdc80 | MmuINT0031273 | chr16:45118363-45122959 | IR-S | 0,1911747 | 0,3035195 | -0,112344827 | 0,872 |
| Arntl | MmuEX0006229 | chr7:120422971-120423096 | C2 | 0,8389886 | 0,951268 | -0,112279349 | 0,85 |
| Arfgap1 | MmuALTA0001881-1/2 | chr2:180702307-180702519 | Alt3 | 0,5865742 | 0,6985279 | -0,111953688 | 0,8493333 |
| Cdc16 | MmuEX0010339 | chr8:13758998-13759052 | S | 0,806557 | 0,9182388 | -0,111681717 | 0,8526667 |
| Pstk | MmuEX0037613 | chr7:138517050-138517344 | S | 0,6811754 | 0,7926626 | -0,11148717 | 0,8473333 |
| Prrc2c | MmuALTA0014278-1/2 | chr1:164653241-164653412 | Alt3 | 0,2437823 | 0,3551295 | -0,111347172 | 0,8593333 |
| Tmem39a | MmuEX0047961 | chr16:38575856-38575939 | S | 0,8366273 | 0,9477744 | -0,111147156 | 0,9333333 |
| Mtmr11 | MmuEX0030063 | chr3:95968356-95968491 | S | 0,8471985 | 0,9582376 | -0,111039059 | 0,914 |
| Eif2ak4 | MmuEX0016515 | chr2:118230783-118230885 | S | 0,8684732 | 0,9792872 | -0,110813963 | 0,8293333 |
| Zfml | MmuEX0053002 | chr6:83931678-83931779 | C1 | 0,2949814 | 0,4055253 | -0,110543931 | 0,8253333 |
| Cgnl1 | MmuALTD0003083-1/2 | chr9:71619316- | Alt5 | 0,7599611 | 0,8679934 | -0,108032235 | 0,8366667 |
| Plb1 | MmuEX0035556 | chr5:32627766-32627809 | S | 0,8209311 | 0,9284829 | -0,107551786 | 0,8413333 |
| Igdcc4 | MmuEX0023770 | chr9:64972419-64972583 | S | 0,8320373 | 0,9395162 | -0,107478857 | 0,9213333 |
| Arfgap3 | MmuALTA0001885-1/2 | chr15:83140680-83140811 | Alt3 | 0,1483152 | 0,2554752 | -0,107160007 | 0,8073333 |
| Ncor2 | MmuALTD0009193-1/2 | chr5:125507172-125507294 | Alt5 | 0,5100857 | 0,6168141 | -0,106728338 | 0,8006667 |
| Plb1 | MmuEX0035557 | chr5:32628472-32628580 | S | 0,800764 | 0,9066867 | -0,105922718 | 0,8213333 |
| Nelf | MmuEX0031485 | chr2:24913485-24913490 | MIC | 0,065704 | 0,1713461 | -0,105642009 | 0,8353333 |
| Kif4 | MmuALTA0009695-1/2 | chrX:97909220-97909346 | Alt3 | 0,0648301 | 0,1700661 | -0,105236014 | 0,8086667 |
| Cda | MmuEX0010292 | chr4:137907099-137907210 | S | 0,8257685 | 0,9299397 | -0,104171155 | 0,91 |
| 5730469M10Rik | MmuALTA0000599-1/2 | chr14:41817330-41817492 | Alt3 | 0,6603025 | 0,7638922 | -0,103589723 | 0,9013333 |
| Plcg1 | MmuINT0121525 | chr2:160579748-160580020 | IR-S | 0,1149734 | 0,2182585 | -0,103285028 | 0,882 |
| Phf6 | MmuEX0034779 | chrX:50270343-50270444 | C2 | 0,8504822 | 0,9533676 | -0,102885395 | 0,836 |
| Ssh3 | MmuEX0045049 | chr19:4267747-4267978 | C3 | 0,795896 | 0,8984539 | -0,102557869 | 0,8333333 |
| Plekhj1 | MmuEX0035798 | chr10:80260655-80260722 | S | 0,672185 | 0,7738169 | -0,101631882 | 0,888 |
| Ikbkap | MmuEX0023862 | chr4:56787889-56787962 | S | 0,8565629 | 0,9580132 | -0,101450315 | 0,88 |
| Strada | MmuEX0045526 | chr11:106032268-106032391 | S | 0,8681843 | 0,9686617 | -0,100477445 | 0,892 |
| Sgsm3 | MmuALTD0012747-1/2 | chr15:80833790-80833875 | Alt5 | 0,8585732 | 0,9565324 | -0,097959155 | 0,8413333 |
| Tnpo3 | MmuALTA0018602-1/2 | chr6:29504883-29504962 | Alt3 | 0,8661573 | 0,9638323 | -0,097675057 | 0,9906667 |
| Map4k3 | MmuEX0027789 | chr17:81028626-81028688 | S | 0,6408269 | 0,737983 | -0,097156033 | 0,8313333 |
| Nhej1 | MmuEX0031694 | chr1:75014743-75014849 | S | 0,8041598 | 0,9006952 | -0,096535403 | 0,8453333 |
| Ypel3 | MmuEX0052574 | chr7:133921568-133921611 | S | 0,8358928 | 0,9318137 | -0,095920955 | 0,926 |
| Aldoc | MmuEX0004623 | chr11:78139437-78139636 | S | 0,8649017 | 0,9606447 | -0,095743036 | 0,9233333 |
| Crtc2 | MmuEX0012798 | chr3:90063076-90063144 | S | 0,6226972 | 0,718181 | -0,095483787 | 0,8213333 |

| | | | | | | | |
|---|---|---|---|---|---|---|---|
| Dgka | MmuEX0014397 | chr10:128168380-128168468 | S | 0,8314131 | 0,9268635 | -0,09545043 | 0,8226667 |
| Pkmyt1 | MmuINT0120788 | chr17:23869792-23870789 | IR-S | 0,0721441 | 0,1673 | -0,095155873 | 0,8546667 |
| Hcfc1 | MmuALTA0008418-1/2 | chrX:71192534-71192853 | Alt3 | 0,6523508 | 0,7471562 | -0,094805424 | 0,8146667 |
| Ankrd11 | MmuEX0005001 | chr8:125424502-125424598 | S | 0,1575116 | 0,2520934 | -0,094581864 | 0,842 |
| Tpx2 | MmuALTD0014706-1/2 | chr2:-152673845 | Alt5 | 0,5687491 | 0,6627698 | -0,094020696 | 0,8546667 |
| Supv3l1 | MmuALTD0013858-1/3 | chr10:61906143-61906311 | Alt5 | 0,8693623 | 0,9626975 | -0,093335273 | 0,8073333 |
| Gm98 | MmuEX0021244 | chr19:10303143-10303406 | S | 0,7326759 | 0,8257307 | -0,093054758 | 0,814 |
| Ccnt1 | MmuEX0010085 | chr15:98377176-98377339 | S | 0,8565989 | 0,9496098 | -0,093010995 | 0,8446667 |
| Ercc1 | MmuINT0058725 | chr7:19940522-19940641 | IR-S | 0,1079107 | 0,2004439 | -0,092533172 | 0,812 |
| Nae1 | MmuEX0030804 | chr8:107051278-107051349 | S | 0,850793 | 0,9427523 | -0,091959279 | 0,856 |
| Tmem161a | MmuEX0047785 | chr8:72700744-72700824 | C3 | 0,779516 | 0,870924 | -0,091407944 | 0,844 |
| Ubp1 | MmuEX0050295 | chr9:113868494-113868601 | S | 0,3630995 | 0,4544094 | -0,091309857 | 0,8173333 |
| Tle6 | MmuEX0047448 | chr10:81062780-81062860 | C1 | 0,873347 | 0,9643958 | -0,09104888 | 0,8973333 |
| Tank | MmuINT0156890 | chr2:61488346-61491435 | IR-C | 0,1363223 | 0,2271398 | -0,090817461 | 0,8673333 |
| Gmpr2 | MmuINT0071744 | chr14:56294570-56295591 | IR-S | 0,0609904 | 0,151763 | -0,090772581 | 0,8726667 |
| Eml2 | MmuEX0016785 | chr7:19787179-19787276 | S | 0,5593494 | 0,6499512 | -0,090601843 | 0,8246667 |
| Nup214 | MmuEX0032768 | chr2:31845755-31845881 | S | 0,8869195 | 0,9772394 | -0,090319981 | 0,9106667 |
| Phf20l1 | MmuEX0034753 | chr15:66462429-66462528 | S | 0,8921779 | 0,9815791 | -0,089401182 | 0,8006667 |
| Rbm5 | MmuEX0039182 | chr9:107662107-107662180 | S | 0,8966366 | 0,9859963 | -0,089359659 | 0,9373333 |
| Tspan17 | MmuEX0049335 | chr13:54897327-54897443 | S | 0,8586109 | 0,9477367 | -0,08912578 | 0,8753333 |
| Tpcn2 | MmuEX0048563 | chr7:152442419-152442501 | C2 | 0,8929207 | 0,981913 | -0,088992317 | 0,8173333 |
| 0610037P05Rik | MmuEX0000066 | chr16:14313909-14314106 | C2 | 0,8302135 | 0,918782 | -0,08856846 | 0,9093333 |
| Crtc2 | MmuEX0012795 | chr3:90061039-90061140 | C2 | 0,682401 | 0,7707774 | -0,088376408 | 0,8073333 |
| Zfp334 | MmuALTA0020582-1/2 | chr2:-165207383 | Alt3 | 0,0383216 | 0,1262951 | -0,087973586 | 0,8326667 |
| Sema3f | MmuALTD0012567-1/2 | chr9:107592017-107592110 | Alt5 | 0,8850566 | 0,97219 | -0,087133431 | 0,8746667 |
| Slc39a10 | MmuEX0043270 | chr1:46874908-46874988 | S | 0,8896325 | 0,9762224 | -0,086589877 | 0,9166667 |
| Mgat5 | MmuALTA0010915-1/2 | chr1:129203214-129203593 | Alt3 | 0,1594591 | 0,2453861 | -0,085926956 | 0,8133333 |
| 2310037I24Rik | MmuEX0000923 | chr15:98361754-98361868 | C3 | 0,0799454 | 0,1654669 | -0,085521544 | 0,8213333 |
| Tnrc6a | MmuEX0048382 | chr7:130318940-130318972 | S | 0,8829847 | 0,9684413 | -0,085456613 | 0,8693333 |
| Rer1 | MmuALTD0011754-1/2 | chr4:154457241-154457345 | Alt5 | 0,0717396 | 0,1570057 | -0,08526614 | 0,886 |
| Mdm2 | MmuEX0028331 | chr10:117146756-117146840 | S | 0,8355692 | 0,9202132 | -0,084644036 | 0,9026667 |
| Baiap2 | MmuEX0007649 | chr11:119858803-119859004 | S | 0,8813238 | 0,9657221 | -0,084398334 | 0,8426667 |
| Rin1 | MmuALTA0015265-1/2 | chr19:5051774-5051832 | Alt3 | 0,0633696 | 0,1476444 | -0,084274778 | 0,8173333 |
| Srek1 | MmuEX0044884 | chr13:104554376-104554499 | S | 0,1142729 | 0,1983654 | -0,084092409 | 0,8146667 |
| Tcf4 | MmuEX0046677 | chr18:69506933-69507024 | C2 | 0,8884951 | 0,9725733 | -0,084078171 | 0,9073333 |
| Trp53bp1 | MmuEX0049084 | chr2:121095955-121096045 | S | 0,8710434 | 0,9548362 | -0,083792717 | 0,832 |
| Fbxo9 | MmuEX0018925 | chr9:77942952-77943066 | S | 0,8855864 | 0,9691353 | -0,083548917 | 0,8606667 |
| Nt5c | MmuEX0032475 | chr11:115352452-115352512 | S | 0,8442576 | 0,9274049 | -0,083147248 | 0,9486667 |
| Taz | MmuEX0046361 | chrX:71534444-71534521 | C2 | 0,8914998 | 0,9743221 | -0,082822279 | 0,8673333 |
| Zfp740 | MmuEX0053594 | chr15:102039184-102039307 | S | 0,79732 | 0,8800224 | -0,082702417 | 0,874 |
| Lipa | MmuALTA0010065-1/2 | chr19:34599228-34599332 | Alt3 | 0,0691366 | 0,1517978 | -0,082661255 | 0,8286667 |
| Crtc2 | MmuEX0012796 | chr3:90061217-90061333 | C3 | 0,7017613 | 0,7839038 | -0,082142475 | 0,8386667 |
| Mfsd11 | MmuALTD0008478-1/2 | chr11:116722837-116722981 | Alt5 | 0,8915589 | 0,9731993 | -0,081640444 | 0,864 |
| Ablim3 | MmuINT0009131 | chr18:61968616-61970964 | IR-S | 0,0518255 | 0,1332581 | -0,081432638 | 0,8146667 |
| Naga | MmuEX0030814 | chr15:82168668-82168744 | S | 0,898523 | 0,9799032 | -0,081380242 | 0,9213333 |
| Med22 | MmuINT0097426 | chr2:26761514-26763612 | IR-S | 0,8866078 | 0,9678484 | -0,081240667 | 0,854 |
| Bag6 | MmuEX0007606 | chr17:35283702-35283848 | S | 0,5108394 | 0,5917009 | -0,080861455 | 0,8613333 |
| Crtc2 | MmuEX0012797 | chr3:90062387-90062448 | C3 | 0,6969783 | 0,7776384 | -0,080660051 | 0,822 |
| Pcyox1l | MmuEX0034095 | chr18:61863007-61863048 | S | 0,1317144 | 0,2120427 | -0,080328371 | 0,8366667 |
| Chd2 | MmuALTD0003110-1/2 | chr7:80586655-80586685 | Alt5 | 0,0286812 | 0,1089589 | -0,080277684 | 0,8113333 |
| Mtss1 | MmuEX0030166 | chr15:58778881-58779075 | C3 | 0,8298963 | 0,9096934 | -0,079797066 | 0,8746667 |
| Zfml | MmuEX0053001 | chr6:83928988-83929903 | C1 | 0,8762065 | 0,9559297 | -0,079723108 | 0,8806667 |
| D6Wsu116e | MmuEX0013632 | chr6:116204431-116204511 | S | 0,8614894 | 0,9408365 | -0,079347104 | 0,9013333 |
| E130309D02Rik | MmuALTD0004586-1/2 | chr5:144076582- | Alt5 | 0,8417398 | 0,9209925 | -0,079252694 | 0,812 |
| Rrp36 | MmuEX0040605 | chr17:46809655-46809796 | C2 | 0,7772997 | 0,8563711 | -0,079071396 | 0,8386667 |
| Tbcel | MmuEX0046565 | chr9:42252490-42252671 | S | 0,815544 | 0,8944391 | -0,078895142 | 0,8753333 |
| Sun1 | MmuALTA0017589-1/2 | chr5:139714785-139714893 | Alt3 | 0,2646829 | 0,3434744 | -0,078791503 | 0,8306667 |
| 2700029M09Rik | MmuEX0001175 | chr8:63375535-63375633 | C3 | 0,8654866 | 0,9440122 | -0,078525639 | 0,8113333 |
| Hsdl2 | MmuEX0023325 | chr4:59606109-59606207 | S | 0,8456623 | 0,9235314 | -0,07786905 | 0,8253333 |
| Lyn | MmuEX0027367 | chr4:3669589-3669634 | C1 | 0,8983439 | 0,9760818 | -0,077737948 | 0,82 |
| Nptn | MmuALTD0009528-1/2 | chr9:58491339-58491491 | Alt5 | 0,1213367 | 0,1982069 | -0,076870145 | 0,8266667 |
| Prmt5 | MmuEX0037247 | chr14:55135334-55135452 | S | 0,8355651 | 0,9124284 | -0,076863394 | 0,866 |
| Kirrel | MmuEX0025646 | chr3:86901664-86901828 | S | 0,8994507 | 0,9760516 | -0,07660085 | 0,8873333 |

| | | | | | | | |
|---|---|---|---|---|---|---|---|
| Ankrd13b | MmuINT0015420 | chr11:77285381-77285515 | IR-S | 0,8828828 | 0,9592844 | -0,076401565 | 0,88 |
| Clcn7 | MmuEX0011608 | chr17:25293746-25293797 | S | 0,8565431 | 0,9328127 | -0,076269541 | 0,8473333 |
| Ttf2 | MmuALTA0019095-1/2 | chr3:100758114-100758242 | Alt3 | 0,0327104 | 0,1089258 | -0,076215353 | 0,8206667 |
| Fam114a2 | MmuALTD0005173-1/2 | chr11:57327597-57327790 | Alt5 | 0,8902454 | 0,9664043 | -0,076158926 | 0,8726667 |
| Ppip5k2 | MmuALTA0013876-1/2 | chr1:99637401-99637519 | Alt3 | 0,1084772 | 0,1846021 | -0,076124886 | 0,8213333 |
| Tada2a | MmuEX0046146 | chr11:83931750-83931809 | S | 0,8936759 | 0,9696512 | -0,075975295 | 0,8633333 |
| Napg | MmuINT0105631 | chr18:63151534-63153965 | IR-S | 0,0255005 | 0,1014431 | -0,075942627 | 0,8786667 |
| Ipo13 | MmuALTA0009268-1/2 | chr4:117576403-117576509 | Alt3 | 0,0316015 | 0,1074845 | -0,07588298 | 0,846 |
| Pcif1 | MmuINT0116133 | chr2:164704990-164708274 | IR-S | 0,0846829 | 0,1605082 | -0,075825334 | 0,8086667 |
| Gale | MmuINT0067228 | chr4:135522032-135522284 | IR-S | 0,060263 | 0,1358746 | -0,075611593 | 0,8533333 |
| Ide | MmuEX0023575 | chr19:37364534-37364678 | C3 | 0,0393236 | 0,1144393 | -0,075115737 | 0,8993333 |
| 0610007P22Rik | MmuEX0000030 | chr17:25379104-25379167 | S | 0,8209081 | 0,8958068 | -0,074898676 | 0,9206667 |
| Asl | MmuINT0019804 | chr5:130499967-130500163 | IR-S | 0,1980609 | 0,2727898 | -0,074728942 | 0,8113333 |
| Mtmr14 | MmuEX0030080 | chr6:113219495-113219553 | S | 0,8812172 | 0,9559015 | -0,074684357 | 0,842 |
| Cnot1 | MmuALTA0004325-1/2 | chr8:98285223-98285336 | Alt3 | 0,056235 | 0,1308714 | -0,074636417 | 0,8793333 |
| Ift80 | MmuEX0023730 | chr3:68739769-68739843 | S | 0,9020276 | 0,9766171 | -0,074589425 | 0,8826667 |
| Adck5 | MmuINT0011724 | chr15:76425697-76425766 | IR-S | 0,0467649 | 0,1208047 | -0,074039806 | 0,8226667 |
| Snx14 | MmuEX0044238 | chr9:88295561-88295587 | S | 0,0885921 | 0,1626251 | -0,074033037 | 0,818 |
| Echdc2 | MmuEX0016195 | chr4:107842320-107842387 | S | 0,8927867 | 0,9667281 | -0,07394143 | 0,8953333 |
| Hsdl2 | MmuEX0023328 | chr4:59610129-59610227 | C2 | 0,9071617 | 0,9808165 | -0,073654845 | 0,8726667 |
| 4933427D14Rik | MmuEX0001873 | chr11:72015914-72016052 | S | 0,8975653 | 0,9711757 | -0,073610486 | 0,8586667 |
| Picalm | MmuALTA0013222-1/2 | chr7:97330750-97330878 | Alt3 | 0,102073 | 0,1754313 | -0,073358363 | 0,844 |
| Trp53inp1 | MmuEX0049097 | chr4:11095044-11095081 | S | 0,5699951 | 0,643016 | -0,07302088 | 0,8186667 |
| Mef2a | MmuEX0028559 | chr7:74389407-74389430 | S | 0,0398313 | 0,1124604 | -0,072629028 | 0,9146667 |
| Stxbp5 | MmuEX0045641 | chr10:9497819-9497869 | C1 | 0,9057084 | 0,9780467 | -0,072338317 | 0,9053333 |
| Atg16l2 | MmuEX0006630 | chr7:108447520-108447622 | S | 0,8868871 | 0,9589537 | -0,072066611 | 0,8373333 |
| Ap3d1 | MmuEX0005453 | chr10:80179219-80179364 | C2 | 0,8967651 | 0,9685243 | -0,071759211 | 0,824 |
| Nthl1 | MmuEX0032515 | chr17:24771670-24771840 | S | 0,9084683 | 0,9801912 | -0,071722898 | 0,9166667 |
| Pnpla6 | MmuEX0036053 | chr8:3537969-3538085 | C3 | 0,895969 | 0,9675259 | -0,071556899 | 0,8553333 |
| Irak1 | MmuEX0024482 | chrX:71268405-71268536 | S | 0,8865139 | 0,9577614 | -0,07124752 | 0,898 |
| Tdrd7 | MmuEX0046837 | chr4:46031402-46031488 | S | 0,9107171 | 0,9818913 | -0,071174215 | 0,908 |
| Ubxn8 | MmuEX0050426 | chr8:34743974-34744099 | S | 0,9127998 | 0,9838115 | -0,071011615 | 0,9293333 |
| Cdk2ap2 | MmuALTD0002919-1/2 | chr19:4097817-4097907 | Alt5 | 0,1330213 | 0,2039503 | -0,070929071 | 0,876 |
| Anapc16 | MmuEX0004851 | chr10:59465166-59465215 | C1 | 0,2331475 | 0,3040312 | -0,070883762 | 0,8213333 |
| Hsf1 | MmuEX0023331 | chr15:76330561-76330626 | S | 0,7951784 | 0,8660618 | -0,070883479 | 0,8626667 |
| ORF61 | MmuALTA0012469-1/3 | chr10:79441141-79441169 | Alt3 | 0,0798433 | 0,1500755 | -0,070232196 | 0,864 |
| Med14 | MmuALTA0010782-1/3 | chrX:12324975-12325103 | Alt3 | 0,9076454 | 0,9776828 | -0,070037412 | 0,864 |
| Zmynd8 | MmuEX0053991 | chr2:165709679-165709739 | S | 0,0709462 | 0,140737 | -0,06979072 | 0,8133333 |
| Acer3 | MmuEX0003470 | chr7:105366566-105366611 | S | 0,9109092 | 0,9804519 | -0,069542682 | 0,8946667 |
| Fanca | MmuEX0018526 | chr8:125796220-125796332 | S | 0,9098238 | 0,9784116 | -0,068587743 | 0,8946667 |
| Chka | MmuEX0011300 | chr19:3871003-3871062 | C2 | 0,1504018 | 0,2186319 | -0,068230097 | 0,8533333 |
| Stx5a | MmuALTD0013798-1/2 | chr19:8817843-8817913 | Alt5 | 0,890502 | 0,9587237 | -0,068221672 | 0,8566667 |
| Rbm39 | MmuEX0039147 | chr2:156004616-156004688 | S | 0,2218962 | 0,2900972 | -0,068200945 | 0,8826667 |
| Impact | MmuEX0024095 | chr18:13134488-13134540 | S | 0,8981259 | 0,9662241 | -0,068098213 | 0,914 |
| Fam176b | MmuEX0018106 | chr4:125826128-125826289 | S | 0,7928569 | 0,8609372 | -0,068080265 | 0,9233333 |
| Prpf40a | MmuALTA0014214-1/2 | chr2:53004858-53004949 | Alt3 | 0,755817 | 0,8238891 | -0,068072156 | 0,8073333 |
| Irak1 | MmuEX0024486 | chrX:71267524-71267588 | C3 | 0,9015151 | 0,9693237 | -0,067808512 | 0,8326667 |
| Megf8 | MmuEX0028608 | chr7:26119811-26119974 | S | 0,9144313 | 0,9817313 | -0,067300061 | 0,842 |
| Stx16 | MmuEX0045570 | chr2:173919430-173919573 | S | 0,9151827 | 0,9818932 | -0,066710509 | 0,9173333 |
| Rhebl1 | MmuEX0039725 | chr15:98709439-98709486 | S | 0,8059649 | 0,8725565 | -0,066591589 | 0,8266667 |
| Klhdc5 | MmuEX0025728 | chr6:147050088-147050281 | S | 0,9115647 | 0,9779717 | -0,06640696 | 0,8573333 |
| Map4k4 | MmuEX0027796 | chr1:40068495-40068686 | S | 0,6736001 | 0,7400021 | -0,066401988 | 0,816 |
| Zyx | MmuEX0054111 | chr6:42301269-42301361 | S | 0,7424483 | 0,8088151 | -0,066366845 | 0,8106667 |
| Picalm | MmuEX0035018 | chr7:97342783-97342806 | S | 0,3407291 | 0,4069238 | -0,066194758 | 0,832 |
| Sgta | MmuALTD0012748-1/3 | chr10:80522873- | Alt5 | 0,8329168 | 0,8990918 | -0,066175067 | 0,8866667 |
| Top1mt | MmuINT0163850 | chr15:75487645-75488113 | IR-S | 0,0339897 | 0,0998191 | -0,06582939 | 0,814 |
| Vegfb | MmuEX0051323 | chr19:7060780-7060976 | S | 0,9113588 | 0,9771387 | -0,065779895 | 0,9066667 |
| Fblim1 | MmuEX0018705 | chr4:141157898-141158005 | C3 | 0,095138 | 0,1608837 | -0,065745716 | 0,8846667 |
| Exo1 | MmuALTA0006647-1/2 | chr1:177823104-177823192 | Alt3 | 0,040973 | 0,1066379 | -0,065664901 | 0,8093333 |
| Polr3a | MmuEX0036306 | chr14:25295235-25295338 | S | 0,9062238 | 0,9718768 | -0,065653019 | 0,8473333 |
| Dph1 | MmuALTD0004436-1/4 | chr11:75003901-75004021 | Alt5 | 0,8870615 | 0,9526318 | -0,065570282 | 0,8086667 |
| Ccnt1 | MmuINT0031772 | chr15:98377026-98377175 | IR-S | 0,0405382 | 0,1059673 | -0,065429112 | 0,826 |
| Bnip2 | MmuEX0008099 | chr9:69852113-69852148 | S | 0,1870393 | 0,2524149 | -0,065375575 | 0,842 |

| | | | | | | | |
|---|---|---|---|---|---|---|---|
| Myef2 | MmuEX0030334 | chr2:124923299-124923370 | S | 0,3917399 | 0,4569664 | -0,065226542 | 0,852 |
| Kntc1 | MmuEX0025859 | chr5:124205683-124205803 | S | 0,9036859 | 0,9686462 | -0,06496024 | 0,8033333 |
| Gtf3c1 | MmuALTA0008284-1/2 | chr7:132802571-132802725 | Alt3 | 0,0301008 | 0,0949765 | -0,064875781 | 0,9113333 |
| Ctdp1 | MmuEX0013022 | chr18:80630905-80631067 | S | 0,9191807 | 0,9839871 | -0,064806366 | 0,9033333 |
| Dapk3 | MmuEX0013762 | chr10:80653787-80653813 | S | 0,8439674 | 0,9082761 | -0,064308773 | 0,8206667 |
| Susd1 | MmuEX0045801 | chr4:59382403-59382509 | C3 | 0,8312409 | 0,8953062 | -0,064065297 | 0,806 |
| Trrap | MmuINT0166368 | chr5:145598193-145600269 | IR-S | 0,0462986 | 0,1102209 | -0,06392229 | 0,8686667 |
| Dgka | MmuEX0014395 | chr10:128171144-128171193 | S | 0,8978061 | 0,9616759 | -0,063869797 | 0,8293333 |
| Myst2 | MmuEX0030642 | chr11:95152834-95152896 | S | 0,7020155 | 0,7652955 | -0,063280076 | 0,814 |
| Hsdl2 | MmuEX0023327 | chr4:59609735-59609838 | C3 | 0,9175584 | 0,980716 | -0,063157538 | 0,8573333 |
| Sec24b | MmuEX0041495 | chr3:129723456-129723560 | S | 0,5625496 | 0,6254742 | -0,062924574 | 0,806 |
| Sf1 | MmuEX0041898 | chr19:6375658-6375862 | S | 0,334966 | 0,3976079 | -0,062641841 | 0,8466667 |
| Ppox | MmuEX0036654 | chr1:173210000-173210132 | S | 0,9257774 | 0,9883304 | -0,062553016 | 0,9333333 |
| Yipf1 | MmuEX0052538 | chr4:107009042-107009129 | S | 0,9245796 | 0,9862429 | -0,061663333 | 0,928 |
| Plekha5 | MmuEX0035717 | chr6:140521386-140521487 | S | 0,8971594 | 0,9583842 | -0,061224779 | 0,8566667 |
| Pigx | MmuEX0035099 | chr16:32087380-32087593 | S | 0,8798613 | 0,941061 | -0,061199713 | 0,8393333 |
| Trim24 | MmuALTD0014777-1/2 | chr6:37895521-37895687 | Alt5 | 0,7167443 | 0,7778556 | -0,061111319 | 0,8206667 |
| Zfp516 | MmuEX0053402 | chr18:83162462-83162581 | S | 0,921829 | 0,9827088 | -0,060879772 | 0,9026667 |
| D6Wsu116e | MmuEX0013631 | chr6:116177364-116177552 | S | 0,8805798 | 0,9414338 | -0,060854023 | 0,8373333 |
| Tcof1 | MmuEX0046741 | chr18:60991820-60991963 | S | 0,8629627 | 0,9233767 | -0,060413978 | 0,9193333 |
| Sun1 | MmuEX0045737 | chr5:139706727-139706792 | C3 | 0,9309874 | 0,9911346 | -0,060147129 | 0,898 |
| Map2k7 | MmuEX0027697 | chr8:4244861-4245003 | S | 0,9180676 | 0,9781436 | -0,060076 | 0,9373333 |
| Prune | MmuINT0127594 | chr3:95072246-95085506 | IR-S | 0,0565985 | 0,1166417 | -0,060043172 | 0,83 |
| Dido1 | MmuEX0014686 | chr2:180425227-180425338 | S | 0,0467828 | 0,1066938 | -0,059910976 | 0,804 |
| Clcn7 | MmuEX0011609 | chr17:25294434-25294519 | S | 0,9271042 | 0,9869035 | -0,059799266 | 0,932 |
| Fcho2 | MmuALTA0007148-1/2 | chr13:99513366-99513490 | Alt3 | 0,021065 | 0,0806856 | -0,059620617 | 0,844 |
| Tmem50b | MmuEX0047998 | chr16:91583521-91583633 | S | 0,9241752 | 0,9833821 | -0,059206874 | 0,8286667 |
| Drap1 | MmuALTA0005846-1/2 | chr19:5423607-5423653 | Alt3 | 0,8715319 | 0,9303197 | -0,058787822 | 0,9366667 |
| Sipa1 | MmuINT0143848 | chr19:5652874-5653998 | IR-C | 0,0383053 | 0,0970273 | -0,058722013 | 0,812 |
| Htra2 | MmuEX0023442 | chr6:83002692-83002797 | S | 0,8781887 | 0,9366253 | -0,058436537 | 0,898 |
| Lmo7 | MmuEX0026624 | chr14:102317157-102317261 | S | 0,9082141 | 0,9664919 | -0,058277746 | 0,8393333 |
| Sigirr | MmuALTD0012812-1/2 | chr7:148282519-148282535 | Alt5 | 0,8730359 | 0,9309578 | -0,057921869 | 0,8773333 |
| Gtpbp2 | MmuEX0022162 | chr17:46302727-46302946 | S | 0,9249384 | 0,9827025 | -0,057764067 | 0,8933333 |
| Rock2 | MmuEX0040284 | chr12:16935740-16935840 | S | 0,9178161 | 0,9750796 | -0,057263549 | 0,85 |
| Snapc1 | MmuEX0044080 | chr12:75083465-75083566 | S | 0,8988441 | 0,9560499 | -0,0572058 | 0,828 |
| Fastkd2 | MmuEX0018663 | chr1:63782419-63782542 | S | 0,8817543 | 0,9389018 | -0,057147483 | 0,8026667 |
| Gnl3 | MmuALTD0006118-1/2 | chr14:31830013-31830093 | Alt5 | 0,8805983 | 0,9374256 | -0,056827229 | 0,846 |
| Tnrc6a | MmuEX0048379 | chr7:130316747-130316923 | S | 0,9191078 | 0,9757779 | -0,056670114 | 0,86 |
| Prpsap1 | MmuALTA0014241-1/2 | chr11:116340966-116341138 | Alt3 | 0,9220242 | 0,9786462 | -0,056621982 | 0,872 |
| Clk3 | MmuEX0011722 | chr9:57606434-57606530 | S | 0,8876653 | 0,9442267 | -0,056561364 | 0,92 |
| Atf7 | MmuEX0006593 | chr15:102393745-102393841 | S | 0,9306796 | 0,9870104 | -0,056330871 | 0,8213333 |
| Tmem20 | MmuEX0047880 | chr19:38474924-38475104 | S | 0,9090442 | 0,9651301 | -0,056085948 | 0,8226667 |
| P2rx4 | MmuEX0033383 | chr5:123174603-123174683 | S | 0,8217096 | 0,8777302 | -0,056020608 | 0,844 |
| Otud4 | MmuALTA0012620-1/2 | chr8:82190615-82190738 | Alt3 | 0,9258557 | 0,9817891 | -0,055933383 | 0,862 |
| Unc5b | MmuEX0050631 | chr10:60240138-60240302 | C1 | 0,8792162 | 0,9348975 | -0,05568135 | 0,9533333 |
| Metap1d | MmuEX0028695 | chr2:71360590-71360687 | S | 0,9287198 | 0,9843217 | -0,055601927 | 0,8913333 |
| Atg9a | MmuEX0006681 | chr1:75181030-75181147 | S | 0,9022147 | 0,9575551 | -0,055340391 | 0,8446667 |
| Psen1 | MmuEX0037504 | chr12:85040305-85040371 | S | 0,923603 | 0,9788145 | -0,055211492 | 0,896 |
| Cntrob | MmuEX0012034 | chr11:69116354-69116468 | S | 0,9217556 | 0,9769495 | -0,055193931 | 0,8546667 |
| Lrrc14 | MmuEX0026925 | chr15:76543831-76544415 | S | 0,91037 | 0,9652892 | -0,054919257 | 0,8113333 |
| Isyna1 | MmuEX0024549 | chr8:73118519-73118647 | S | 0,9045155 | 0,9594028 | -0,054887319 | 0,9173333 |
| Tnk2 | MmuALTD0014613-1/2 | chr16:32670039-32670302 | Alt5 | 0,0350653 | 0,0898962 | -0,05483084 | 0,8293333 |
| Inf2 | MmuEX0024138 | chr12:113841923-113842064 | S | 0,9318404 | 0,9866197 | -0,054779354 | 0,938 |
| Sgsm3 | MmuEX0042101 | chr15:80837877-80838020 | S | 0,9314712 | 0,9860522 | -0,054580939 | 0,8033333 |
| Vegfb | MmuEX0051324 | chr19:7060511-7060584 | S | 0,9238792 | 0,9783441 | -0,054464921 | 0,9393333 |
| Fbf1 | MmuEX0018697 | chr11:116012103-116012161 | S | 0,9323616 | 0,9866009 | -0,054239296 | 0,808 |
| Papola | MmuEX0033600 | chr12:107071310-107071372 | S | 0,889698 | 0,9435887 | -0,053890676 | 0,8033333 |
| Terf2 | MmuEX0046964 | chr8:109618722-109618852 | S | 0,9239508 | 0,9777862 | -0,053835428 | 0,82 |
| Aatf | MmuEX0002922 | chr11:84324985-84325080 | S | 0,921951 | 0,9754157 | -0,053464745 | 0,9006667 |
| Med20 | MmuEX0028520 | chr17:47755758-47756011 | C1 | 0,911816 | 0,9648627 | -0,053046732 | 0,81 |
| Blm | MmuEX0008028 | chr7:87618915-87619110 | S | 0,9208838 | 0,9736118 | -0,05272796 | 0,8233333 |
| Plaa | MmuEX0035534 | chr4:94256572-94256765 | S | 0,9053482 | 0,9579092 | -0,052560964 | 0,836 |
| Lztr1 | MmuEX0027436 | chr16:17524401-17524550 | S | 0,901482 | 0,9538453 | -0,052363298 | 0,8233333 |

| | | | | | | | |
|---|---|---|---|---|---|---|---|
| Npr2 | MmuEX0032213 | chr4:43656207-43656281 | S | 0,9341341 | 0,9857696 | -0,051635497 | 0,8926667 |
| AA408296 | MmuALTD0000665-1/3 | chr1:194956238- | Alt5 | 0,923652 | 0,9752804 | -0,051628384 | 0,8146667 |
| Cpt1c | MmuEX0012628 | chr7:52220454-52220636 | S | 0,933221 | 0,9847986 | -0,051577695 | 0,8646667 |
| Azin1 | MmuEX0007181 | chr15:38436942-38437101 | S | 0,1780855 | 0,2292625 | -0,051177063 | 0,8173333 |
| Flot1 | MmuEX0019350 | chr17:35961224-35961314 | S | 0,8765688 | 0,9276 | -0,051031245 | 0,886 |
| Lepre1 | MmuALTA0009994-1/3 | chr4:118914288-118914409 | Alt3 | 0,933772 | 0,9847588 | -0,050986772 | 0,8853333 |
| Rnf34 | MmuEX0040181 | chr5:123311698-123311916 | S | 0,9336067 | 0,9841651 | -0,050558435 | 0,866 |
| Srrm1 | MmuEX0044966 | chr4:134902231-134902360 | S | 0,0201509 | 0,0705394 | -0,050388584 | 0,8553333 |
| Mllt6 | MmuEX0029243 | chr11:97537453-97537519 | S | 0,9277897 | 0,977953 | -0,050163313 | 0,84 |
| 5730469M10Rik | MmuALTD0000562-1/2 | chr14:41826987- | Alt5 | 0,9070016 | 0,8567972 | 0,050204366 | 0,8213333 |
| Mtmr11 | MmuINT0101894 | chr3:95974570-95975008 | IR-S | 0,092044 | 0,0414674 | 0,050576607 | 0,8026667 |
| Stk25 | MmuINT0154086 | chr1:95522184-95522357 | IR-S | 0,1648711 | 0,1141689 | 0,050702197 | 0,8053333 |
| Sh2b1 | MmuINT0142937 | chr7:133612843-133614635 | IR-S | 0,0948951 | 0,0440791 | 0,050816028 | 0,82 |
| Thop1 | MmuALTA0018165-1/2 | chr10:80540661-80540760 | Alt3 | 0,0724702 | 0,0214363 | 0,051033816 | 0,8386667 |
| Tnpo3 | MmuINT0163475 | chr6:29504963-29505152 | IR-S | 0,0918527 | 0,0407077 | 0,051144951 | 0,866 |
| Xrcc1 | MmuINT0176964 | chr7:25352946-25355234 | IR-S | 0,1106431 | 0,0594393 | 0,051203739 | 0,8053333 |
| Lrrc42 | MmuINT0093029 | chr4:106906590-106908404 | IR-S | 0,074224 | 0,0230065 | 0,051217547 | 0,8693333 |
| Pkn1 | MmuINT0120799 | chr8:86196232-86197200 | IR-S | 0,0737856 | 0,0224177 | 0,051367904 | 0,802 |
| Mfge8 | MmuEX0028788 | chr7:86288164-86288274 | S | 0,2707689 | 0,2191984 | 0,051570559 | 0,93 |
| Rexo4 | MmuEX0039487 | chr2:26813999-26814087 | C2 | 0,9787914 | 0,9271728 | 0,051618663 | 0,85 |
| Nfat5 | MmuEX0031580 | chr8:109871567-109871757 | C3 | 0,9626324 | 0,9109333 | 0,05169913 | 0,804 |
| Arhgef5 | MmuINT0018639 | chr6:43215743-43222303 | IR-S | 0,084003 | 0,0320916 | 0,05191135 | 0,8073333 |
| Bpnt1 | MmuEX0008140 | chr1:187161985-187162111 | S | 0,9162525 | 0,8642913 | 0,051961235 | 0,8633333 |
| Pan3 | MmuEX0033551 | chr5:148331081-148331328 | C3 | 0,9579407 | 0,9059168 | 0,052023924 | 0,8333333 |
| Tbrg4 | MmuINT0157805 | chr11:6520147-6520744 | IR-S | 0,2091838 | 0,1569318 | 0,052251967 | 0,812 |
| Aurkb | MmuINT0022300 | chr11:68863979-68864401 | IR-S | 0,0974391 | 0,0451841 | 0,052254957 | 0,8926667 |
| Marveld1 | MmuINT0095940 | chr19:42224050-42225287 | IR-S | 0,1100797 | 0,057675 | 0,052404644 | 0,8206667 |
| Clasrp | MmuINT0036794 | chr7:20166508-20166729 | IR-S | 0,0772263 | 0,0246872 | 0,052539133 | 0,872 |
| Hsf1 | MmuINT0079205 | chr15:76330767-76330846 | IR-S | 0,1249457 | 0,0723902 | 0,052555545 | 0,8246667 |
| Trmt2a | MmuINT0165646 | chr16:18252342-18252494 | IR-S | 0,118647 | 0,065908 | 0,052738981 | 0,814 |
| Hjurp | MmuINT0077956 | chr1:90166875-90169303 | IR-C | 0,1343469 | 0,081531 | 0,052815865 | 0,8946667 |
| Tuft1 | MmuINT0168279 | chr3:94426068-94426609 | IR-S | 0,0977568 | 0,0447678 | 0,05298897 | 0,84 |
| 2500003M10Rik | MmuINT0002587 | chr3:90304030-90306037 | IR-S | 0,1023592 | 0,0493571 | 0,05300215 | 0,8346667 |
| Acadvl | MmuINT0009473 | chr11:69826586-69826673 | IR-S | 0,0812593 | 0,0282269 | 0,053032475 | 0,8326667 |
| Traf7 | MmuEX0048730 | chr17:24653450-24653541 | C1 | 0,8549857 | 0,8017595 | 0,053226185 | 0,8413333 |
| Ppp1r16a | MmuEX0036712 | chr15:76523869-76523944 | S | 0,9608965 | 0,9076181 | 0,053278412 | 0,808 |
| Traf7 | MmuINT0164563 | chr17:24647317-24647400 | IR-S | 0,0785362 | 0,0251755 | 0,05336067 | 0,9346667 |
| Cacnb3 | MmuINT0027924 | chr15:98471182-98471389 | IR-S | 0,0820453 | 0,0286218 | 0,053423503 | 0,8626667 |
| Gga1 | MmuINT0068629 | chr15:78716375-78718540 | IR-S | 0,0973421 | 0,0438131 | 0,053528942 | 0,84 |
| Fam195a | MmuINT0061458 | chr17:26001298-26001538 | IR-S | 0,080721 | 0,0271197 | 0,05360138 | 0,8293333 |
| Hsf1 | MmuEX0023336 | chr15:76328584-76328720 | S | 0,8971121 | 0,8434704 | 0,053641683 | 0,82 |
| Zfp498 | MmuEX0053380 | chr5:146047177-146047378 | S | 0,9675719 | 0,9139247 | 0,053647163 | 0,8386667 |
| Pcif1 | MmuEX0033977 | chr2:164714886-164714972 | C3 | 0,9568075 | 0,9030128 | 0,053794695 | 0,8273333 |
| Atxn7l3 | MmuINT0022253 | chr11:102153522-102153755 | IR-S | 0,0864964 | 0,0326163 | 0,053880079 | 0,8233333 |
| Cdk11b | MmuEX0010536 | chr4:155016594-155016684 | S | 0,9850335 | 0,9309614 | 0,054072191 | 0,8806667 |
| Grk6 | MmuINT0074221 | chr13:55553205-55553495 | IR-S | 0,085863 | 0,0317038 | 0,054159231 | 0,906 |
| Zranb1 | MmuEX0054026 | chr7:140162869-140163022 | C2 | 0,989543 | 0,9352383 | 0,054304642 | 0,8453333 |
| Phrf1 | MmuALTA0013192-1/2 | chr7:148443243-148443397 | Alt3 | 0,9718447 | 0,9175295 | 0,05431518 | 0,8006667 |
| 1810032O08Rik | MmuINT0001724 | chr11:116534053-116535251 | IR-S | 0,1044883 | 0,0500333 | 0,054454999 | 0,802 |
| Ercc1 | MmuINT0058724 | chr7:19938055-19939674 | IR-S | 0,0837965 | 0,029308 | 0,054488476 | 0,8066667 |
| Oxa1l | MmuINT0114193 | chr14:54983180-54986392 | IR-C | 0,0887778 | 0,0339921 | 0,054785696 | 0,8693333 |
| Adrbk1 | MmuINT0012180 | chr19:4291331-4291589 | IR-S | 0,089615 | 0,0345387 | 0,055076315 | 0,8573333 |
| Usp21 | MmuINT0171436 | chr1:173213873-173214092 | IR-C | 0,1129805 | 0,0578902 | 0,055090308 | 0,8126667 |
| Cinp | MmuEX0011439 | chr12:112115051-112115180 | S | 0,9691636 | 0,9140597 | 0,055103817 | 0,8393333 |
| Csnk1e | MmuINT0043069 | chr15:79251441-79255269 | IR-C | 0,1203181 | 0,0650453 | 0,055272763 | 0,8306667 |
| Fbrs | MmuINT0062998 | chr7:134628757-134628866 | IR-C | 0,0804732 | 0,0251673 | 0,05530589 | 0,832 |
| 1810014F10Rik | MmuINT0001673 | chr7:147285606-147285777 | IR-S | 0,1401394 | 0,0847458 | 0,055393629 | 0,822 |
| Agxt2l2 | MmuEX0004351 | chr11:51412153-51412242 | S | 0,9809492 | 0,9254912 | 0,05545802 | 0,8626667 |
| Cep55 | MmuALTD0003042-1/2 | chr19:38132279-38132473 | Alt5 | 0,9490901 | 0,8935969 | 0,055493173 | 0,812 |
| Ppan | MmuINT0124382 | chr9:20695457-20695526 | IR-S | 0,1170112 | 0,0612728 | 0,055738457 | 0,8446667 |
| Lhpp | MmuEX0026395 | chr7:139822165-139822352 | S | 0,9724688 | 0,9166234 | 0,055845413 | 0,828 |
| Fbrs | MmuEX0018747 | chr7:134631388-134631468 | S | 0,9894576 | 0,9336113 | 0,05584627 | 0,94 |
| Ephx1 | MmuALTA0006488-1/2 | chr1:182924010-182924176 | Alt3 | 0,0694498 | 0,0134603 | 0,0559895 | 0,9206667 |

| | | | | | | | |
|---|---|---|---|---|---|---|---|
| Alg8 | MmuINT0014149 | chr7:104532317-104535311 | IR-S | 0,1204432 | 0,064415 | 0,056028194 | 0,8866667 |
| Arhgap27 | MmuEX0005853 | chr11:103200339-103200422 | S | 0,9720185 | 0,915975 | 0,056043483 | 0,8893333 |
| Crim1 | MmuEX0012728 | chr17:78680586-78680706 | C2 | 0,9525291 | 0,8964131 | 0,056116015 | 0,8053333 |
| Hira | MmuEX0022943 | chr16:18947237-18947335 | S | 0,0857214 | 0,0294787 | 0,056242698 | 0,84 |
| Mrps2 | MmuEX0029731 | chr2:28324741-28324848 | S | 0,0713263 | 0,0150221 | 0,056304148 | 0,932 |
| Pcif1 | MmuEX0033976 | chr2:164714594-164714766 | C2 | 0,9568668 | 0,9001551 | 0,056711688 | 0,8333333 |
| Rxrb | MmuINT0138112 | chr17:34173910-34174111 | IR-S | 0,0878396 | 0,0311211 | 0,056718478 | 0,8126667 |
| Ipo4 | MmuINT0082786 | chr14:56247825-56247903 | IR-S | 0,1192524 | 0,0623795 | 0,056872902 | 0,8086667 |
| Nol7 | MmuINT0109458 | chr13:43494165-43494879 | IR-S | 0,095118 | 0,0382316 | 0,056886382 | 0,8413333 |
| Pvt1 | MmuINT0129846 | chr15:62007104-62073728 | IR-S | 0,0960882 | 0,0391585 | 0,056929646 | 0,8193333 |
| Spag9 | MmuINT0150947 | chr11:93973461-93975486 | IR-S | 0,118452 | 0,0615078 | 0,056944178 | 0,818 |
| Dusp6 | MmuINT0054000 | chr10:98726726-98727177 | IR-S | 0,0716378 | 0,0146591 | 0,056978707 | 0,9953333 |
| Commd10 | MmuEX0012374 | chr18:47123314-47123424 | C3 | 0,9862819 | 0,9292659 | 0,057016031 | 0,928 |
| Tatdn1 | MmuEX0046340 | chr15:58755460-58755523 | S | 0,9496299 | 0,8925251 | 0,057104774 | 0,832 |
| Ska1 | MmuEX0042478 | chr18:74362244-74362341 | S | 0,9712165 | 0,914041 | 0,05717554 | 0,826 |
| Ranbp3 | MmuINT0131512 | chr17:56846662-56847273 | IR-S | 0,0939629 | 0,0367296 | 0,057233265 | 0,8646667 |
| Ercc1 | MmuEX0017237 | chr7:19937982-19938054 | S | 0,9355209 | 0,8782526 | 0,057268342 | 0,812 |
| Spsb3 | MmuEX0044809 | chr17:25027513-25027700 | S | 0,9816605 | 0,9242004 | 0,057460091 | 0,8886667 |
| Sigirr | MmuINT0143637 | chr7:148277401-148277701 | IR-S | 0,3132091 | 0,2556435 | 0,057565602 | 0,866 |
| Kat5 | MmuINT0085267 | chr19:5605684-5607550 | IR-S | 0,0819885 | 0,0242191 | 0,057769323 | 0,858 |
| Rin1 | MmuINT0135051 | chr19:5053276-5053598 | IR-S | 0,0931067 | 0,035275 | 0,057831736 | 0,822 |
| Git1 | MmuINT0068966 | chr11:77314617-77316330 | IR-S | 0,0917318 | 0,033782 | 0,057949819 | 0,8393333 |
| Zfyve27 | MmuEX0053824 | chr19:42263687-42263733 | S | 0,9747899 | 0,9166441 | 0,058145832 | 0,8613333 |
| Dgka | MmuEX0014392 | chr10:128165312-128165397 | S | 0,9825221 | 0,9243554 | 0,058166618 | 0,904 |
| Parp3 | MmuINT0115442 | chr9:106375538-106375763 | IR-S | 0,1161416 | 0,0579083 | 0,058233382 | 0,8013333 |
| Gstm2 | MmuINT0074614 | chr3:107788601-107788951 | IR-S | 0,0713821 | 0,0130099 | 0,058372233 | 0,9226667 |
| Rab6 | MmuEX0038387 | chr7:107778352-107778457 | C3 | 0,8427674 | 0,7842701 | 0,058497362 | 0,834 |
| Psrc1 | MmuINT0128262 | chr3:108187204-108187291 | IR-S | 0,2728287 | 0,2142918 | 0,058536855 | 0,8266667 |
| Fam119a | MmuINT0060622 | chr1:64661784-64662846 | IR-S | 0,0812377 | 0,0226777 | 0,058559966 | 0,826 |
| Ncln | MmuINT0106543 | chr10:80956285-80958824 | IR-S | 0,0859421 | 0,0273661 | 0,058576002 | 0,9273333 |
| Nudt1 | MmuEX0032597 | chr5:140808377-140808427 | S | 0,9048131 | 0,8461495 | 0,058663529 | 0,812 |
| Cxxc1 | MmuINT0044416 | chr18:74378310-74378385 | IR-S | 0,0844981 | 0,025788 | 0,058710152 | 0,9113333 |
| Polrmt | MmuINT0124031 | chr10:79200255-79200344 | IR-S | 0,1025777 | 0,043844 | 0,058733752 | 0,8113333 |
| Rnpepl1 | MmuINT0136062 | chr1:94813049-94813287 | IR-S | 0,0773048 | 0,0184385 | 0,058866318 | 0,878 |
| Haus6 | MmuINT0076035 | chr4:86250194-86251540 | IR-S | 0,0881754 | 0,0292811 | 0,058894289 | 0,808 |
| Gbf1 | MmuEX0020159 | chr19:46344748-46344841 | S | 0,9812119 | 0,9222916 | 0,058920313 | 0,8793333 |
| Psip1 | MmuINT0127791 | chr4:83107146-83109498 | IR-C | 0,1235875 | 0,0646021 | 0,058985427 | 0,8313333 |
| Nubp2 | MmuINT0111394 | chr17:25022563-25022663 | IR-S | 0,1007979 | 0,0416678 | 0,059130113 | 0,8326667 |
| Otud5 | MmuINT0114115 | chrX:7451080-7451358 | IR-S | 0,0817627 | 0,0226014 | 0,059161219 | 0,852 |
| Ide | MmuEX0023574 | chr19:37365160-37365304 | S | 0,976757 | 0,917286 | 0,059470936 | 0,9193333 |
| Micall1 | MmuINT0098687 | chr15:78953417-78954930 | IR-S | 0,0897975 | 0,0302548 | 0,059542654 | 0,8026667 |
| 2700007P21Rik | MmuEX0001172 | chr2:106812925-106813319 | C2 | 0,1033494 | 0,043802 | 0,059547374 | 0,8153333 |
| Ddx17 | MmuALTD0004079-1/3 | chr15:79362108-79362298 | Alt5 | 0,7583282 | 0,6986605 | 0,059667671 | 0,8433333 |
| Ampd3 | MmuEX0004816 | chr7:117934702-117934861 | C3 | 0,9788842 | 0,9190907 | 0,059793564 | 0,8106667 |
| Inf2 | MmuINT0082086 | chr12:113843074-113843583 | IR-S | 0,089777 | 0,0299727 | 0,059804311 | 0,8826667 |
| Csnk1g2 | MmuINT0043093 | chr10:80101979-80102324 | IR-S | 0,1118794 | 0,0518776 | 0,060001819 | 0,9113333 |
| Id1 | MmuINT0080143 | chr2:152562482-152562707 | IR-S | 0,0922599 | 0,0322343 | 0,06002561 | 0,9993333 |
| Scly | MmuEX0041234 | chr1:93206374-93206480 | S | 0,9622991 | 0,9021627 | 0,060136369 | 0,8606667 |
| Apba3 | MmuINT0016991 | chr10:80732330-80733712 | IR-S | 0,0883143 | 0,0279831 | 0,060331156 | 0,854 |
| Tmem134 | MmuINT0161475 | chr19:4127640-4127759 | IR-S | 0,1189764 | 0,0585952 | 0,060381157 | 0,854 |
| R3hdm2 | MmuEX0038219 | chr10:126899262-126899339 | S | 0,9512997 | 0,8908595 | 0,060440196 | 0,8466667 |
| Eml3 | MmuINT0057207 | chr19:9011978-9012052 | IR-S | 0,1204689 | 0,0600277 | 0,060441219 | 0,8146667 |
| Zmynd8 | MmuINT0180616 | chr2:165646189-165653707 | IR-C | 0,1139522 | 0,0529135 | 0,061038747 | 0,836 |
| Hgs | MmuINT0077598 | chr11:120344416-120344702 | IR-S | 0,1521313 | 0,0910582 | 0,061073086 | 0,806 |
| Alg9 | MmuEX0004668 | chr9:50591363-50591433 | S | 0,9542773 | 0,893094 | 0,061183314 | 0,8246667 |
| Snhg1 | MmuINT0149687 | chr19:8799537-8799711 | IR-S | 0,2299587 | 0,1687266 | 0,061232075 | 0,8506667 |
| Mff | MmuEX0028787 | chr1:82743658-82743717 | C2 | 0,5469946 | 0,4857098 | 0,061284796 | 0,826 |
| Llgl1 | MmuINT0091354 | chr11:60522396-60522936 | IR-S | 0,1064659 | 0,0451699 | 0,061296033 | 0,8086667 |
| Eif4g2 | MmuINT0056593 | chr7:118219227-118220209 | IR-C | 0,1153149 | 0,0539695 | 0,06134537 | 0,8786667 |
| Pcsk7 | MmuEX0034084 | chr9:45734817-45734911 | C1 | 0,9865739 | 0,9251535 | 0,061420478 | 0,8433333 |
| Aph1a | MmuINT0017118 | chr3:95700284-95700616 | IR-S | 0,9202598 | 0,8587794 | 0,061480383 | 0,95 |
| Rpp21 | MmuINT0136861 | chr17:36394652-36394727 | IR-S | 0,1769545 | 0,1152755 | 0,061678951 | 0,8046667 |
| Acin1 | MmuEX0003479 | chr14:55272051-55272115 | S | 0,2053971 | 0,143678 | 0,061719109 | 0,8193333 |

| | | | | | | | |
|---|---|---|---|---|---|---|---|
| Ranbp10 | MmuEX0038697 | chr8:108298187-108298335 | S | 0,9823311 | 0,920438 | 0,061893147 | 0,828 |
| Ftsjd2 | MmuINT0066431 | chr17:29835675-29836164 | IR-C | 0,0984449 | 0,0365352 | 0,061909703 | 0,8786667 |
| Dtx3 | MmuINT0053774 | chr10:126631691-126632749 | IR-S | 0,0782592 | 0,0163058 | 0,06195341 | 0,9206667 |
| Cd97 | MmuINT0032583 | chr8:86253425-86257312 | IR-C | 0,1005087 | 0,0385017 | 0,062007045 | 0,81 |
| Fam49b | MmuEX0018344 | chr15:63802485-63802523 | C2 | 0,86899 | 0,8069792 | 0,062010764 | 0,8326667 |
| Cast | MmuEX0009297 | chr13:74917951-74918022 | S | 0,9796 | 0,9171789 | 0,062421066 | 0,8153333 |
| Bet1l | MmuINT0024795 | chr7:148040721-148040972 | IR-S | 0,1455856 | 0,0831392 | 0,062446401 | 0,8413333 |
| Spns1 | MmuINT0151896 | chr7:133517588-133518580 | IR-S | 0,1702246 | 0,107772 | 0,062452646 | 0,8226667 |
| Cx3cl1 | MmuEX0013248 | chr8:97301928-97302048 | S | 0,9807558 | 0,9182741 | 0,062481774 | 0,868 |
| Tia1 | MmuINT0160065 | chr6:86375065-86375443 | IR-S | 0,1107139 | 0,0480267 | 0,062687206 | 0,842 |
| Clk3 | MmuINT0037424 | chr9:57599635-57599921 | IR-S | 0,0925189 | 0,0297779 | 0,062740939 | 0,8666667 |
| Adrbk1 | MmuINT0012165 | chr19:4290006-4290073 | IR-S | 0,0833389 | 0,0203444 | 0,06299447 | 0,9486667 |
| Sri | MmuEX0044919 | chr5:8059353-8059422 | S | 0,9549121 | 0,8917781 | 0,063134007 | 0,8826667 |
| Gltscr2 | MmuINT0069537 | chr7:16523239-16523522 | IR-S | 0,1227324 | 0,059388 | 0,063344376 | 0,8673333 |
| Cxxc1 | MmuINT0044419 | chr18:74379222-74379491 | IR-S | 0,1345044 | 0,0708536 | 0,063650851 | 0,848 |
| Znf512b | MmuALTA0020891-1/3 | chr2:181323784-181323949 | Alt3 | 0,0886273 | 0,0249604 | 0,063666887 | 0,844 |
| Egr1 | MmuINT0055886 | chr18:35021441-35022118 | IR-S | 0,1013937 | 0,0373911 | 0,06400258 | 0,8573333 |
| Gtse1 | MmuINT0075106 | chr15:85702012-85704069 | IR-S | 0,0951432 | 0,0310676 | 0,064075595 | 0,8573333 |
| Hnrnpd | MmuEX0023101 | chr5:100405553-100405609 | S | 0,5671751 | 0,5029533 | 0,064221806 | 0,8606667 |
| Hgsnat | MmuALTA0008585-1/2 | chr8:27083314-27083423 | Alt3 | 0,0876475 | 0,023303 | 0,064344473 | 0,83 |
| Nxf1 | MmuINT0112178 | chr19:8839522-8841286 | IR-S | 0,3454367 | 0,2810845 | 0,064352194 | 0,8233333 |
| Top2a | MmuINT0163887 | chr11:98854728-98855077 | IR-S | 0,3283833 | 0,2638778 | 0,064505505 | 0,84 |
| Lysmd2 | MmuINT0094108 | chr9:75483525-75485018 | IR-S | 0,1001622 | 0,0356472 | 0,064514984 | 0,8013333 |
| Foxm1 | MmuINT0065643 | chr6:128322622-128322794 | IR-S | 0,1765812 | 0,1117783 | 0,064802871 | 0,808 |
| Grpel2 | MmuINT0074315 | chr18:61879510-61885879 | IR-S | 0,1045877 | 0,0397554 | 0,06483228 | 0,8073333 |
| Lypla2 | MmuINT0094085 | chr4:135525445-135525515 | IR-S | 0,122587 | 0,057534 | 0,065052947 | 0,8306667 |
| Rbbp5 | MmuINT0132414 | chr1:134386306-134386999 | IR-S | 0,0962656 | 0,0310969 | 0,065168687 | 0,8386667 |
| Map4k4 | MmuEX0027798 | chr1:40060608-40060832 | C2 | 0,5342253 | 0,4690072 | 0,065218063 | 0,846 |
| Actn1 | MmuEX0003624 | chr12:81271170-81271235 | C3 | 0,4643682 | 0,3991421 | 0,065226088 | 0,8353333 |
| Rnpep | MmuEX0040242 | chr1:137168939-137169055 | S | 0,8612515 | 0,7959294 | 0,06532211 | 0,816 |
| Thop1 | MmuINT0159914 | chr10:80539796-80540657 | IR-S | 0,1024555 | 0,0369024 | 0,065553014 | 0,8453333 |
| Usp21 | MmuINT0171433 | chr1:173215170-173215385 | IR-C | 0,1048465 | 0,0390078 | 0,06583871 | 0,8666667 |
| 4933424B01Rik | MmuINT0004974 | chr6:146498508-146498608 | IR-S | 0,1204428 | 0,0542566 | 0,06618618 | 0,8153333 |
| Cxxc1 | MmuINT0044414 | chr18:74377651-74377739 | IR-S | 0,128468 | 0,0620623 | 0,066405703 | 0,8173333 |
| Syvn1 | MmuINT0156325 | chr19:6050686-6050940 | IR-C | 0,1092955 | 0,0428799 | 0,066415632 | 0,838 |
| Pan2 | MmuINT0114946 | chr10:127756095-127757303 | IR-S | 0,0921731 | 0,0257382 | 0,066434878 | 0,8533333 |
| Jub | MmuINT0085080 | chr14:55190671-55192228 | IR-C | 0,1248484 | 0,058366 | 0,066482325 | 0,814 |
| 2310021P13Rik | MmuINT0002177 | chr14:21537215-21537648 | IR-S | 0,1020802 | 0,0354234 | 0,066656878 | 0,914 |
| Plxnb1 | MmuINT0122681 | chr9:108998160-109001784 | IR-S | 0,1058321 | 0,0391609 | 0,066671201 | 0,806 |
| Magt1 | MmuINT0094619 | chrX:103166366-103166774 | IR-S | 0,9296865 | 0,8629742 | 0,066712324 | 0,8646667 |
| Mbnl1 | MmuEX0028097 | chr3:60418605-60418658 | S | 0,291869 | 0,2251207 | 0,066748315 | 0,808 |
| Hspa13 | MmuEX0023366 | chr16:75758877-75758972 | S | 0,1627082 | 0,0959474 | 0,066760814 | 0,8253333 |
| Rrp1b | MmuALTD0012250-1/2 | chr17:32182874-32182956 | Alt5 | 0,9231743 | 0,8563959 | 0,066778469 | 0,816 |
| Slc9a8 | MmuINT0148011 | chr2:167249798-167260281 | IR-C | 0,1197066 | 0,0529281 | 0,066778495 | 0,8006667 |
| Trim47 | MmuINT0165258 | chr11:115969323-115969579 | IR-C | 0,109404 | 0,0425805 | 0,066823533 | 0,8366667 |
| 2610301G19Rik | MmuEX0001138 | chr14:70542732-70542895 | S | 0,984782 | 0,9173447 | 0,067437328 | 0,9206667 |
| Rbm5 | MmuALTA0014985-1/2 | chr9:107652039-107652127 | Alt3 | 0,1021621 | 0,0343395 | 0,067822645 | 0,8086667 |
| Itfg3 | MmuINT0083542 | chr17:26351673-26352129 | IR-S | 0,0993948 | 0,031454 | 0,067940767 | 0,8366667 |
| Srsf6 | MmuINT0152721 | chr2:162757879-162759161 | IR-S | 0,1679852 | 0,0999854 | 0,06799986 | 0,9826667 |
| Mapkap1 | MmuEX0027862 | chr2:34274628-34274785 | C2 | 0,0985336 | 0,0304419 | 0,068091691 | 0,828 |
| Hdac6 | MmuINT0076412 | chrX:7516659-7516731 | IR-C | 0,1514744 | 0,0832811 | 0,068193313 | 0,8506667 |
| Sec24c | MmuEX0041502 | chr14:21494953-21495081 | S | 0,1172782 | 0,0488434 | 0,068434781 | 0,8933333 |
| Dido1 | MmuINT0049507 | chr2:180419882-180422180 | IR-S | 0,1082657 | 0,0396917 | 0,068574001 | 0,856 |
| Lmnb2 | MmuINT0091567 | chr10:80367519-80367630 | IR-S | 0,1251825 | 0,0562315 | 0,068951026 | 0,822 |
| Racgap1 | MmuALTA0014746-1/2 | chr15:99473314-99473398 | Alt3 | 0,1631816 | 0,0938957 | 0,069285872 | 0,848 |
| Mff | MmuALTA0010879-1/2 | chr1:82731994-82732082 | Alt3 | 0,5987742 | 0,5292961 | 0,069478127 | 0,8653333 |
| Gcn1l1 | MmuINT0068236 | chr5:116065609-116066359 | IR-S | 0,1115988 | 0,0421083 | 0,069490549 | 0,8526667 |
| Commd4 | MmuINT0040778 | chr9:57004363-57004581 | IR-S | 0,1145417 | 0,0445284 | 0,070013391 | 0,852 |
| Hdac7 | MmuINT0076453 | chr15:97640471-97641283 | IR-S | 0,1308466 | 0,0606727 | 0,070173921 | 0,8766667 |
| Slc25a44 | MmuEX0042934 | chr3:88219910-88220037 | S | 0,9418334 | 0,8715297 | 0,070303773 | 0,8153333 |
| Ehmt1 | MmuEX0016475 | chr2:24708185-24708322 | S | 0,9786136 | 0,9081446 | 0,070468947 | 0,8226667 |
| Med24 | MmuINT0097479 | chr11:98577840-98579223 | IR-S | 0,1443417 | 0,0736573 | 0,070684323 | 0,8273333 |
| Sltm | MmuEX0043731 | chr9:70431739-70431919 | S | 0,9899901 | 0,9192874 | 0,0707027 | 0,846 |

| | | | | | | | |
|---|---|---|---|---|---|---|---|
| Mark3 | MmuEX0027961 | chr12:112856529-112856594 | S | 0,9849615 | 0,9142356 | 0,07072595 | 0,9486667 |
| Pnma5 | MmuINT0123081 | chrX:70279667-70280502 | IR-S | 0,4836766 | 0,4127696 | 0,070907009 | 0,88 |
| Nfat5 | MmuEX0031586 | chr8:109890079-109890227 | C3 | 0,9644071 | 0,8932618 | 0,071145223 | 0,8446667 |
| Slc12a4 | MmuINT0144551 | chr8:108471852-108473037 | IR-S | 0,1493486 | 0,0778233 | 0,071525297 | 0,9033333 |
| Igsf9 | MmuEX0023844 | chr1:174420792-174420903 | C3 | 0,967502 | 0,8958207 | 0,071681259 | 0,9086667 |
| Atxn7l2 | MmuEX0007104 | chr3:108007419-108007540 | S | 0,9765835 | 0,9048378 | 0,071745706 | 0,8066667 |
| Dtx3 | MmuINT0053776 | chr10:126630414-126630785 | IR-C | 0,1761181 | 0,1038548 | 0,0722633 | 0,8453333 |
| Csf1 | MmuEX0012866 | chr3:107559907-107560046 | S | 0,1309285 | 0,058641 | 0,072287476 | 0,8273333 |
| Isg20l2 | MmuINT0083411 | chr3:87734534-87735252 | IR-S | 0,1367704 | 0,0644521 | 0,072318305 | 0,8566667 |
| Orc4 | MmuINT0113508 | chr2:48789143-48792291 | IR-S | 0,0954598 | 0,0230498 | 0,07240999 | 0,898 |
| Dvl1 | MmuINT0054028 | chr4:155227868-155228091 | IR-S | 0,169412 | 0,0967597 | 0,072652369 | 0,876 |
| Rbm28 | MmuINT0132749 | chr6:29105190-29107067 | IR-S | 0,1509651 | 0,0780669 | 0,07289823 | 0,8733333 |
| Smtn | MmuINT0149363 | chr11:3417831-3420754 | IR-S | 0,1912504 | 0,1182994 | 0,072951014 | 0,856 |
| Wdr67 | MmuEX0052001 | chr15:57751486-57751601 | S | 0,8537658 | 0,7807858 | 0,07298 | 0,8173333 |
| Gon4l | MmuEX0021473 | chr3:88662309-88662428 | S | 0,9658858 | 0,8927874 | 0,073098428 | 0,81 |
| Hmha1 | MmuINT0078331 | chr10:79487597-79488153 | IR-S | 0,1823968 | 0,1091975 | 0,073199265 | 0,894 |
| Pold1 | MmuINT0123476 | chr7:51789168-51789254 | IR-S | 0,1352978 | 0,0620475 | 0,073250314 | 0,824 |
| Gigyf2 | MmuEX0020409 | chr1:89230845-89230909 | S | 0,9786783 | 0,9053678 | 0,07331043 | 0,8593333 |
| Mark3 | MmuEX0027960 | chr12:112853607-112853655 | C1 | 0,985441 | 0,9119569 | 0,07348418 | 0,8433333 |
| Mpnd | MmuINT0100349 | chr17:56151767-56151838 | IR-S | 0,1296785 | 0,0561361 | 0,073542436 | 0,8346667 |
| Wdr11 | MmuINT0174948 | chr7:136744125-136746512 | IR-S | 0,1168472 | 0,0430546 | 0,073792607 | 0,8146667 |
| Mtx1 | MmuINT0102245 | chr3:89014106-89014187 | IR-S | 0,1430152 | 0,0688602 | 0,074154986 | 0,908 |
| Cherp | MmuALTA0004076-1/4 | chr8:74985900-74985944 | Alt3 | 0,2148929 | 0,1402158 | 0,074677078 | 0,82 |
| Tsen54 | MmuINT0166510 | chr11:115676394-115676488 | IR-S | 0,1648376 | 0,0899974 | 0,074840221 | 0,8006667 |
| Xab2 | MmuINT0176563 | chr8:3613710-3613798 | IR-S | 0,1373014 | 0,0619824 | 0,07531903 | 0,82 |
| Glipr1 | MmuINT0069262 | chr10:111422739-111422833 | IR-C | 0,3313286 | 0,2558261 | 0,075502517 | 0,9293333 |
| Cpt1c | MmuINT0041893 | chr7:52215307-52215419 | IR-S | 0,2042594 | 0,1287331 | 0,07552628 | 0,806 |
| Tm9sf1 | MmuINT0160864 | chr14:56256970-56259221 | IR-C | 0,163959 | 0,0883164 | 0,075642611 | 0,8526667 |
| Gas8 | MmuALTA0007571-1/2 | chr8:126054733-126054936 | Alt3 | 0,1432626 | 0,0673603 | 0,075902293 | 0,8353333 |
| Tlcd1 | MmuINT0160470 | chr11:77993125-77993448 | IR-S | 0,1644801 | 0,0885224 | 0,075957765 | 0,8006667 |
| Ilkap | MmuEX0024066 | chr1:93287709-93287774 | S | 0,9426429 | 0,8665802 | 0,076062728 | 0,8553333 |
| Snhg12 | MmuINT0149697 | chr4:131864823-131865752 | IR-S | 0,1738396 | 0,0976594 | 0,076180202 | 0,8246667 |
| Tcfap4 | MmuALTD0014106-1/2 | chr16:4559487- | Alt5 | 0,1022286 | 0,026007 | 0,076221637 | 0,8186667 |
| Sbf1 | MmuINT0139006 | chr15:89125744-89125825 | IR-S | 0,0969699 | 0,0207233 | 0,07624654 | 0,956 |
| Ncoa3 | MmuALTA0011807-1/2 | chr2:165874009-165874055 | Alt3 | 0,1456405 | 0,0693283 | 0,076312129 | 0,806 |
| Cyth4 | MmuINT0045563 | chr15:78440305-78441878 | IR-C | 0,1673002 | 0,090626 | 0,076674126 | 0,804 |
| Zfr | MmuEX0053757 | chr15:12047994-12048093 | S | 0,9254986 | 0,8488075 | 0,076691076 | 0,922 |
| Mcm3ap | MmuALTD0008318-1/2 | chr10:75952080-75952321 | Alt5 | 0,9548696 | 0,8781562 | 0,076713379 | 0,8226667 |
| H2-T10 | MmuINT0075658 | chr17:36257232-36257242 | IR-C | 0,0927356 | 0,0159476 | 0,076788032 | 0,926 |
| Smyd5 | MmuINT0149486 | chr6:85390739-85391553 | IR-S | 0,1640009 | 0,0871004 | 0,07690045 | 0,8773333 |
| Naa35 | MmuEX0030749 | chr13:59710122-59710205 | S | 0,9682817 | 0,8912073 | 0,077074369 | 0,8193333 |
| Lrrc68 | MmuEX0027071 | chr7:20122719-20122816 | S | 0,9447315 | 0,8672789 | 0,077452671 | 0,878 |
| Prpf38b | MmuEX0037322 | chr3:108713261-108713313 | S | 0,1606365 | 0,0831268 | 0,077509742 | 0,8693333 |
| Ppan | MmuINT0124378 | chr9:20693884-20694077 | IR-S | 0,2171025 | 0,1393778 | 0,077724713 | 0,836 |
| N4bp2l2 | MmuEX0030701 | chr5:151449576-151449652 | S | 0,9288168 | 0,8508835 | 0,077933362 | 0,8993333 |
| Cd47 | MmuEX0010235 | chr16:49908167-49908191 | C1 | 0,1857464 | 0,1075285 | 0,078217889 | 0,8513333 |
| Polrmt | MmuINT0124032 | chr10:79200108-79200201 | IR-S | 0,129461 | 0,0512115 | 0,078249473 | 0,8293333 |
| Igsf9 | MmuEX0023846 | chr1:174421996-174422141 | C3 | 0,9640827 | 0,8855529 | 0,078529779 | 0,9046667 |
| Kdelc2 | MmuEX0025246 | chr9:53202041-53202237 | S | 0,9803448 | 0,9016892 | 0,078655512 | 0,8173333 |
| Mre11a | MmuEX0029633 | chr9:14614248-14614328 | S | 0,8899892 | 0,8111424 | 0,078846809 | 0,8413333 |
| March7 | MmuINT0095819 | chr2:60083318-60085946 | IR-S | 0,1372753 | 0,058267 | 0,079008342 | 0,8113333 |
| Sdccag3 | MmuINT0140196 | chr2:26241130-26241509 | IR-C | 0,1934352 | 0,1141435 | 0,079291695 | 0,8106667 |
| Tuft1 | MmuEX0049920 | chr3:94443317-94443391 | C1 | 0,8068491 | 0,7275503 | 0,079298861 | 0,84 |
| Nedd4l | MmuEX0031402 | chr18:65327168-65327227 | S | 0,2075422 | 0,1278083 | 0,079733902 | 0,9033333 |
| Ikbkb | MmuINT0081086 | chr8:23770932-23771873 | IR-S | 0,110594 | 0,0300561 | 0,080537888 | 0,876 |
| Gpaa1 | MmuINT0072449 | chr15:76163530-76163606 | IR-S | 0,1828326 | 0,1021767 | 0,080655975 | 0,89 |
| Tnr | MmuINT0163502 | chr1:161780677-161782087 | IR-S | 0,1226352 | 0,041864 | 0,080771225 | 0,8346667 |
| Crim1 | MmuEX0012727 | chr17:78679313-78679555 | S | 0,9612274 | 0,8804003 | 0,080827096 | 0,8833333 |
| Ncor1 | MmuEX0031180 | chr11:62203193-62203294 | C1 | 0,9828088 | 0,901602 | 0,081206803 | 0,8166667 |
| Tlk2 | MmuEX0047474 | chr11:105101675-105101770 | S | 0,9070461 | 0,8257524 | 0,081293604 | 0,8713333 |
| Tjap1 | MmuINT0160350 | chr17:46396433-46396885 | IR-S | 0,2347667 | 0,1534205 | 0,081346212 | 0,8073333 |
| Slc25a37 | MmuINT0145664 | chr14:69863412-69865637 | IR-S | 0,1490192 | 0,0676147 | 0,081404473 | 0,9266667 |
| Mrs2 | MmuINT0101017 | chr13:25085058-25085538 | IR-S | 0,1304588 | 0,0486111 | 0,08184771 | 0,8026667 |

| | | | | | | | |
|---|---|---|---|---|---|---|---|
| Gtf3a | MmuINT0074921 | chr5:147766760-147766943 | IR-C | 0,478461 | 0,3965043 | 0,081956686 | 0,9113333 |
| Senp1 | MmuEX0041655 | chr15:97907343-97907473 | S | 0,1191087 | 0,0371502 | 0,081958519 | 0,902 |
| Eif4g3 | MmuINT0056627 | chr4:137733926-137736461 | IR-C | 0,2335836 | 0,1512667 | 0,082316955 | 0,814 |
| Abhd6 | MmuEX0003237 | chr14:8860751-8860903 | C1 | 0,9660437 | 0,8833261 | 0,082717649 | 0,844 |
| Mospd2 | MmuINT0100134 | chrX:161385287-161385998 | IR-S | 0,1191455 | 0,0362872 | 0,082858271 | 0,844 |
| Llgl1 | MmuALTA0010072-1/2 | chr11:60519943-60520075 | Alt3 | 0,1444517 | 0,0611898 | 0,083261869 | 0,8446667 |
| Gm2382 | MmuEX0021047 | chr9:88598764-88598974 | C1 | 0,1384934 | 0,0550913 | 0,083402118 | 0,862 |
| Nde1 | MmuINT0106827 | chr16:14190326-14191810 | IR-C | 0,2626074 | 0,1787619 | 0,083845526 | 0,8713333 |
| Fbf1 | MmuEX0018696 | chr11:116007750-116007935 | S | 0,9130781 | 0,8289527 | 0,084125391 | 0,8933333 |
| Stx5a | MmuINT0154631 | chr19:8823068-8823144 | IR-S | 0,1635214 | 0,0793842 | 0,084137201 | 0,826 |
| Abi1 | MmuEX0003242 | chr2:22818734-22818748 | MIC | 0,1436753 | 0,0595042 | 0,084171096 | 0,8466667 |
| Taz | MmuINT0157185 | chrX:71529033-71529270 | IR-C | 0,1500669 | 0,0657287 | 0,084338178 | 0,8546667 |
| 5730455O13Rik | MmuINT0005364 | chr19:38296404-38298451 | IR-S | 0,1584526 | 0,0740909 | 0,084361621 | 0,808 |
| Mbnl1 | MmuEX0028095 | chr3:60400641-60400751 | S | 0,1794726 | 0,0950729 | 0,084399755 | 0,8126667 |
| Rpe | MmuALTD0012056-1/2 | chr1:-66747591 | Alt5 | 0,1370095 | 0,052504 | 0,084505566 | 0,8613333 |
| Igsf9 | MmuEX0023845 | chr1:174421746-174421887 | C3 | 0,9562561 | 0,8716784 | 0,084577716 | 0,8866667 |
| Tgds | MmuALTA0018115-1/2 | chr14:118511133-118512281 | Alt3 | 0,1117969 | 0,0271826 | 0,0846143 | 0,818 |
| 2410004B18Rik | MmuINT0002460 | chr3:145601293-145601861 | IR-S | 0,1260991 | 0,0411118 | 0,084987255 | 0,866 |
| Dgcr8 | MmuINT0048675 | chr16:18283377-18283789 | IR-S | 0,1415659 | 0,056571 | 0,084994946 | 0,8113333 |
| Eri3 | MmuEX0017312 | chr4:117237232-117237503 | C2 | 0,9151893 | 0,8300318 | 0,085157499 | 0,89 |
| Usp21 | MmuINT0171428 | chr1:173212934-173213145 | IR-C | 0,2220605 | 0,1368749 | 0,085185582 | 0,844 |
| Rhebl1 | MmuINT0134588 | chr15:98708966-98709438 | IR-S | 0,3315201 | 0,2459796 | 0,085540479 | 0,8566667 |
| Anapc1 | MmuEX0004845 | chr2:128505845-128505945 | S | 0,9583478 | 0,8726952 | 0,085652594 | 0,8053333 |
| Pum2 | MmuEX0038043 | chr12:8740079-8740315 | S | 0,7883708 | 0,7023811 | 0,085989703 | 0,824 |
| Sco2 | MmuINT0139878 | chr15:89204034-89204113 | IR-C | 0,191198 | 0,1050229 | 0,086175063 | 0,842 |
| Sft2d2 | MmuEX0041979 | chr1:167118100-167118185 | S | 0,8708461 | 0,7846283 | 0,086217824 | 0,9286667 |
| 1810032O08Rik | MmuINT0001725 | chr11:116535386-116535613 | IR-S | 0,1929923 | 0,1067458 | 0,086246472 | 0,832 |
| Gatad2a | MmuEX0020135 | chr8:72474552-72474653 | C2 | 0,9537086 | 0,8673296 | 0,086379009 | 0,9666667 |
| C230081A13Rik | MmuEX0008505 | chr9:56166470-56166627 | S | 0,9692038 | 0,8827756 | 0,086428181 | 0,8606667 |
| Cspp1 | MmuEX0012969 | chr1:10106099-10106165 | C2 | 0,9776924 | 0,8906823 | 0,087010093 | 0,892 |
| Cald1 | MmuINT0028216 | chr6:34696016-34703442 | IR-S | 0,2719601 | 0,1849078 | 0,087052242 | 0,8433333 |
| Exosc2 | MmuINT0059851 | chr2:31531425-31532187 | IR-S | 0,1318593 | 0,0443316 | 0,087527667 | 0,8186667 |
| Safb | MmuALTA0015782-1/2 | chr17:56744980-56745080 | Alt3 | 0,901788 | 0,8140731 | 0,08771492 | 0,854 |
| Erf | MmuINT0058824 | chr7:26030551-26030652 | IR-S | 0,2487525 | 0,1609929 | 0,087759561 | 0,9473333 |
| Taf1d | MmuINT0156671 | chr9:15117281-15118433 | IR-S | 0,2630985 | 0,175081 | 0,088017433 | 0,8086667 |
| Ift52 | MmuINT0080570 | chr2:162843270-162848110 | IR-S | 0,1462855 | 0,0580342 | 0,088251374 | 0,8386667 |
| Arhgap23 | MmuALTA0001927-1/2 | chr11:97309484-97309579 | Alt3 | 0,9553613 | 0,8669664 | 0,088394971 | 0,828 |
| BC003331 | MmuALTA0002510-1/2 | chr1:152235653-152235749 | Alt3 | 0,145169 | 0,0565731 | 0,088595888 | 0,8086667 |
| Ing4 | MmuINT0082105 | chr6:124997606-124997810 | IR-S | 0,3048046 | 0,2157425 | 0,089062049 | 0,814 |
| Aim2 | MmuEX0004445 | chr1:175357890-175358029 | C1 | 0,7940641 | 0,7049465 | 0,08911757 | 0,8686667 |
| Dbr1 | MmuEX0013797 | chr9:99478860-99478940 | S | 0,9662237 | 0,877066 | 0,089157762 | 0,8386667 |
| Scaf1 | MmuINT0139153 | chr7:52268645-52268849 | IR-S | 0,1864567 | 0,0972741 | 0,089182594 | 0,814 |
| Arfip2 | MmuINT0017796 | chr7:112784908-112785324 | IR-S | 0,2420649 | 0,152533 | 0,089531868 | 0,8386667 |
| Dph1 | MmuINT0052839 | chr11:74994447-74994794 | IR-S | 0,1390794 | 0,0490201 | 0,090059323 | 0,828 |
| Clk4 | MmuEX0011723 | chr11:51080673-51080833 | C1 | 0,9448811 | 0,8543343 | 0,090546867 | 0,9093333 |
| Irak1 | MmuINT0083219 | chrX:71267589-71267811 | IR-S | 0,2509778 | 0,1602676 | 0,090710203 | 0,87 |
| Dhx30 | MmuINT0049139 | chr9:109988383-109988453 | IR-S | 0,1598615 | 0,0690619 | 0,0907996 | 0,8486667 |
| Cby1 | MmuINT0029856 | chr15:79496182-79496260 | IR-S | 0,1393244 | 0,0484905 | 0,090833899 | 0,8186667 |
| Zfml | MmuALTD0016024-1/2 | chr6:-83864473 | Alt5 | 0,9446024 | 0,8537172 | 0,090885239 | 0,852 |
| Chrnb1 | MmuEX0011383 | chr11:69609097-69609175 | S | 0,810087 | 0,7191448 | 0,090942246 | 0,8153333 |
| Rhebl1 | MmuINT0134585 | chr15:98709919-98710207 | IR-S | 0,2268013 | 0,1358334 | 0,090967862 | 0,8653333 |
| St6galnac6 | MmuINT0153256 | chr2:32463617-32464745 | IR-C | 0,1738341 | 0,0823231 | 0,091510956 | 0,8086667 |
| Yes1 | MmuEX0052527 | chr5:32942704-32942976 | S | 0,9723264 | 0,8806271 | 0,091699301 | 0,8973333 |
| Igsf9 | MmuEX0023843 | chr1:174420420-174420574 | C3 | 0,9636968 | 0,871959 | 0,091737806 | 0,954 |
| Dcaf6 | MmuEX0013850 | chr1:167341595-167341730 | S | 0,9815461 | 0,8894828 | 0,092063249 | 0,8673333 |
| 9430008C03Rik | MmuINT0006146 | chr2:158181778-158182933 | IR-S | 0,1929969 | 0,1008868 | 0,092110101 | 0,9186667 |
| Hook2 | MmuINT0078678 | chr8:87522424-87525053 | IR-C | 0,174999 | 0,0828396 | 0,092159457 | 0,8366667 |
| Sart1 | MmuINT0138866 | chr19:5382360-5382714 | IR-S | 0,2410462 | 0,1485549 | 0,092491247 | 0,8073333 |
| Golph3l | MmuEX0021465 | chr3:95395639-95395827 | S | 0,9707232 | 0,8779859 | 0,092737235 | 0,8873333 |
| Sirt6 | MmuINT0143981 | chr10:81085618-81086736 | IR-S | 0,1554679 | 0,0626925 | 0,092775484 | 0,8253333 |
| BC037034 | MmuINT0023193 | chr5:138701936-138702062 | IR-S | 0,1875847 | 0,094662 | 0,092922638 | 0,8086667 |
| Midn | MmuALTA0010964-1/2 | chr10:79617079-79617217 | Alt3 | 0,379792 | 0,2867663 | 0,09302569 | 0,87 |
| Rab24 | MmuINT0130449 | chr13:55421576-55421672 | IR-S | 0,1364975 | 0,0433509 | 0,093146604 | 0,962 |

| | | | | | | | |
|---|---|---|---|---|---|---|---|
| Ssbp4 | MmuINT0152818 | chr8:73121668-73121911 | IR-S | 0,7622352 | 0,6689352 | 0,093300001 | 0,9273333 |
| Magi1 | MmuEX0027545 | chr6:93644002-93644169 | S | 0,9494457 | 0,8560893 | 0,093356378 | 0,8153333 |
| Kdm1a | MmuEX0025261 | chr4:136137955-136138014 | S | 0,5856106 | 0,4921069 | 0,093503718 | 0,854 |
| Atp9b | MmuEX0006985 | chr18:80933551-80933583 | S | 0,2344563 | 0,14063 | 0,09382627 | 0,804 |
| Trip6 | MmuINT0165591 | chr5:137754318-137754578 | IR-S | 0,2701666 | 0,1763321 | 0,093834545 | 0,864 |
| Arhgap22 | MmuINT0017989 | chr14:34172512-34175628 | IR-S | 0,1379991 | 0,0439578 | 0,094041352 | 0,86 |
| Ift81 | MmuALTD0006923-1/2 | chr5:123064447- | Alt5 | 0,9455176 | 0,8513383 | 0,094179295 | 0,8013333 |
| Rce1 | MmuINT0133172 | chr19:4624182-4624684 | IR-S | 0,1786244 | 0,0843429 | 0,094281528 | 0,9006667 |
| Athl1 | MmuINT0020568 | chr7:148132328-148132400 | IR-S | 0,1619887 | 0,067607 | 0,094381683 | 0,856 |
| Mettl10 | MmuEX0028729 | chr7:140023043-140023259 | C1 | 0,9783788 | 0,8835486 | 0,094830268 | 0,9293333 |
| Dtx3 | MmuALTA0005895-1/2 | chr10:126631466-126631529 | Alt3 | 0,1891188 | 0,0941445 | 0,094974278 | 0,818 |
| Use1 | MmuALTA0019551-1/2 | chr8:73893031-73893202 | Alt3 | 0,8983651 | 0,80256 | 0,095805073 | 0,9033333 |
| Gpr172b | MmuINT0073135 | chr15:76371010-76371101 | IR-S | 0,2280601 | 0,132235 | 0,095825161 | 0,824 |
| Tns3 | MmuEX0048432 | chr11:8500558-8500595 | S | 0,9742752 | 0,8778489 | 0,096426235 | 0,8113333 |
| Tmem87a | MmuEX0048105 | chr2:120200224-120200295 | C3 | 0,4604897 | 0,3640101 | 0,096479639 | 0,8046667 |
| Tecpr1 | MmuINT0158726 | chr5:144957121-144957220 | IR-S | 0,1714745 | 0,0748174 | 0,0966571 | 0,8813333 |
| C230081A13Rik | MmuEX0008507 | chr9:56222183-56222246 | S | 0,9636418 | 0,8666551 | 0,096986622 | 0,8786667 |
| Cchcr1 | MmuINT0031500 | chr17:35666092-35667092 | IR-S | 0,2331386 | 0,1361514 | 0,096987208 | 0,848 |
| Skiv2l | MmuINT0144279 | chr17:34978164-34978386 | IR-S | 0,1720031 | 0,0745359 | 0,097467166 | 0,8153333 |
| Gtf2ird1 | MmuEX0022126 | chr5:134842706-134842765 | S | 0,5664876 | 0,4690168 | 0,097470791 | 0,8206667 |
| 2310037I24Rik | MmuEX0000927 | chr15:98352067-98352193 | S | 0,2616243 | 0,1639597 | 0,097664583 | 0,852 |
| Lzts2 | MmuALTA0010402-1/3 | chr19:45096096-45096326 | Alt3 | 0,1546912 | 0,05645 | 0,098241174 | 0,8766667 |
| Gcat | MmuINT0068051 | chr15:78867305-78867572 | IR-C | 0,135199 | 0,0367941 | 0,098404923 | 0,8486667 |
| Git2 | MmuEX0020455 | chr5:115193785-115193889 | S | 0,7486963 | 0,6502748 | 0,098421568 | 0,8533333 |
| Toe1 | MmuINT0163718 | chr4:116478351-116478589 | IR-S | 0,1489127 | 0,0498154 | 0,099097306 | 0,8713333 |
| Ercc2 | MmuINT0058738 | chr7:19979050-19979124 | IR-S | 0,1685199 | 0,0692327 | 0,099287269 | 0,8266667 |
| Entpd2 | MmuINT0057671 | chr2:25253034-25253520 | IR-S | 0,1347206 | 0,0349637 | 0,099756899 | 0,8493333 |
| Dnm2 | MmuINT0051794 | chr9:21310160-21310779 | IR-S | 0,1411197 | 0,0413379 | 0,099781807 | 0,86 |
| Stxbp1 | MmuEX0045617 | chr2:32650108-32650233 | S | 0,2147982 | 0,1149387 | 0,099859482 | 0,8486667 |
| 5730419I09Rik | MmuINT0005332 | chr6:143048317-143048452 | IR-C | 0,150036 | 0,0498591 | 0,100176896 | 0,832 |
| Ercc5 | MmuALTD0004996-1/2 | chr1:44214989-44215104 | Alt5 | 0,9464284 | 0,8458808 | 0,100547622 | 0,8213333 |
| Tmem168 | MmuEX0047810 | chr6:13555787-13555959 | C1 | 0,2677 | 0,1671394 | 0,100560535 | 0,8086667 |
| Kirrel | MmuEX0025651 | chr3:86944899-86945041 | S | 0,9713259 | 0,8706877 | 0,100638128 | 0,9133333 |
| Pgm5 | MmuINT0118482 | chr19:24783853-24802158 | IR-S | 0,3529745 | 0,2515704 | 0,10140409 | 0,824 |
| Ogg1 | MmuALTD0009774-1/2 | chr6:113278403-113278473 | Alt5 | 0,1851295 | 0,0834096 | 0,101719858 | 0,8166667 |
| Papd5 | MmuALTA0012735-1/2 | chr8:90776426-90776472 | Alt3 | 0,2474741 | 0,1454458 | 0,102028281 | 0,808 |
| Lnp | MmuEX0026643 | chr2:74366149-74366231 | S | 0,3745931 | 0,2712384 | 0,10335467 | 0,8113333 |
| Dcaf8 | MmuALTD0003997-1/3 | chr1:-174078299 | Alt5 | 0,6396813 | 0,5362079 | 0,103473438 | 0,83 |
| Malt1 | MmuEX0027588 | chr18:65623695-65623769 | S | 0,972198 | 0,867452 | 0,104745998 | 0,818 |
| Nf2 | MmuEX0031553 | chr11:4680581-4680625 | S | 0,9215042 | 0,8166129 | 0,10489134 | 0,8166667 |
| Lpin2 | MmuINT0092041 | chr17:71588494-71590151 | IR-S | 0,1756235 | 0,0706669 | 0,104956639 | 0,8186667 |
| Gpr124 | MmuINT0072935 | chr8:28221071-28221523 | IR-S | 0,2105634 | 0,1055446 | 0,105018803 | 0,9146667 |
| 1300001I01Rik | MmuALTA0000080-1/2 | chr11:74477289-74477390 | Alt3 | 0,2242024 | 0,1183646 | 0,10583779 | 0,81 |
| Strn4 | MmuINT0154497 | chr7:17416488-17416813 | IR-C | 0,1577957 | 0,0515256 | 0,106270147 | 0,8286667 |
| Tle6 | MmuALTA0018241-1/2 | chr10:81061822-81061883 | Alt3 | 0,9663804 | 0,8600717 | 0,10630874 | 0,8026667 |
| Canx | MmuINT0028590 | chr11:50110640-50110775 | IR-S | 0,1805933 | 0,0736397 | 0,106953564 | 0,8313333 |
| Rif1 | MmuALTA0015254-1/2 | chr2:51972362-51972470 | Alt3 | 0,1754467 | 0,0683777 | 0,107068984 | 0,826 |
| Pitpnm1 | MmuINT0120267 | chr19:4108194-4108422 | IR-S | 0,1835022 | 0,0761087 | 0,107393525 | 0,818 |
| Nub1 | MmuALTD0009618-1/3 | chr5:-24191840 | Alt5 | 0,9580616 | 0,8501953 | 0,107866337 | 0,8506667 |
| Usp15 | MmuEX0050842 | chr10:122590097-122590183 | S | 0,6627488 | 0,5533333 | 0,109415539 | 0,8386667 |
| 9130404D08Rik | MmuINT0006049 | chr8:72552674-72553106 | IR-S | 0,222681 | 0,1126689 | 0,110012095 | 0,9013333 |
| Acot8 | MmuINT0009931 | chr2:164620680-164621151 | IR-S | 0,2323245 | 0,1221607 | 0,110163839 | 0,82 |
| Ncor2 | MmuALTA0011820-1/2 | chr5:125531281-125531428 | Alt3 | 0,7831923 | 0,6728603 | 0,11033206 | 0,828 |
| Wee1 | MmuEX0052129 | chr7:117274343-117274489 | S | 0,9713598 | 0,8591169 | 0,112242889 | 0,9333333 |
| Dlg4 | MmuEX0014813 | chr11:69840425-69840490 | S | 0,8749818 | 0,7623945 | 0,112587282 | 0,888 |
| D10Ertd610e | MmuINT0045684 | chr10:126621269-126621447 | IR-S | 0,2000382 | 0,0874467 | 0,112591502 | 0,8733333 |
| Ccnl1 | MmuINT0031750 | chr3:65752713-65754401 | IR-C | 0,3029351 | 0,1898807 | 0,113054448 | 0,8806667 |
| 2310022A10Rik | MmuINT0002202 | chr7:28365711-28366369 | IR-S | 0,9722806 | 0,8589698 | 0,113310779 | 0,8386667 |
| Fibp | MmuINT0064520 | chr19:5463305-5463603 | IR-S | 0,2823538 | 0,168753 | 0,113600788 | 0,8153333 |
| Safb2 | MmuINT0138607 | chr17:56715436-56716300 | IR-S | 0,2607887 | 0,1453462 | 0,115442501 | 0,872 |
| Pxk | MmuALTA0014604-1/2 | chr14:8987800-8987859 | Alt3 | 0,5422425 | 0,4265622 | 0,115680244 | 0,822 |
| Igsf9 | MmuINT0081000 | chr1:174420904-174421745 | IR-C | 0,187597 | 0,0718666 | 0,115730388 | 0,8526667 |
| Rere | MmuALTA0015114-1/2 | chr4:149990868-149991588 | Alt3 | 0,6983648 | 0,5819608 | 0,11640408 | 0,836 |

| | | | | | | | |
|---|---|---|---|---|---|---|---|
| Kdm4c | MmuEX0025318 | chr4:73926801-73926915 | S | 0,9657154 | 0,8491409 | 0,116574524 | 0,8866667 |
| Ubr2 | MmuEX0050335 | chr17:47117269-47117367 | C3 | 0,7578447 | 0,6406627 | 0,117181976 | 0,8106667 |
| 2810403A07Rik | MmuINT0003063 | chr3:88514775-88515551 | IR-S | 0,5684822 | 0,451092 | 0,117390155 | 0,9366667 |
| Gas2l1 | MmuINT0067671 | chr11:4964482-4964839 | IR-C | 0,7843065 | 0,6665696 | 0,117736889 | 0,9253333 |
| Slc25a16 | MmuEX0042839 | chr10:62395467-62395530 | S | 0,9206415 | 0,802713 | 0,117928564 | 0,8786667 |
| Mrpl40 | MmuALTA0011257-1/2 | chr16:18875417-18875500 | Alt3 | 0,9482699 | 0,8278741 | 0,120395877 | 0,862 |
| Tatdn1 | MmuALTD0014009-1/2 | chr15:58741132-58741204 | Alt5 | 0,1856037 | 0,0648008 | 0,120802818 | 0,8226667 |
| Zfp740 | MmuINT0179805 | chr15:102039308-102039585 | IR-S | 0,1983438 | 0,0770936 | 0,12125026 | 0,868 |
| Prrc2b | MmuEX0037403 | chr2:32084783-32084864 | S | 0,3711154 | 0,2496312 | 0,121484224 | 0,9066667 |
| Clcn7 | MmuEX0011607 | chr17:25292313-25292482 | S | 0,8700041 | 0,7483302 | 0,121673948 | 0,9206667 |
| Rnf123 | MmuALTA0015329-1/2 | chr9:107964085-107964190 | Alt3 | 0,1653079 | 0,0434658 | 0,121842109 | 0,8766667 |
| Eml3 | MmuINT0057212 | chr19:9013995-9015068 | IR-S | 0,1789372 | 0,0565621 | 0,122375069 | 0,9093333 |
| Wdr74 | MmuALTD0015694-1/2 | chr19:8812338-8812352 | Alt5 | 0,1324167 | 0,0100313 | 0,122385417 | 0,8413333 |
| Asb8 | MmuALTA0002103-1/2 | chr15:97973082-97973241 | Alt3 | 0,3442245 | 0,2202541 | 0,123970448 | 0,8253333 |
| D2Wsu81e | MmuINT0045983 | chr2:30030810-30031233 | IR-S | 0,2014134 | 0,0766717 | 0,124741716 | 0,8706667 |
| Ets1 | MmuEX0017458 | chr9:32545684-32545944 | S | 0,994463 | 0,869087 | 0,125375954 | 0,832 |
| Ldb1 | MmuINT0090373 | chr19:46107832-46108521 | IR-C | 0,6474918 | 0,5220538 | 0,12543804 | 0,846 |
| Ints3 | MmuINT0082557 | chr3:90195834-90195914 | IR-S | 0,9084923 | 0,7829711 | 0,125521204 | 0,878 |
| Ncor2 | MmuINT0106748 | chr5:125506175-125507171 | IR-S | 0,4271202 | 0,3006092 | 0,126511037 | 0,8233333 |
| Susd1 | MmuINT0155279 | chr4:59337780-59342334 | IR-C | 0,2463954 | 0,1197297 | 0,126665739 | 0,8326667 |
| Cdc42bpa | MmuEX0010395 | chr1:182024075-182024317 | S | 0,7681745 | 0,6413013 | 0,12687319 | 0,8373333 |
| Rbbp5 | MmuALTD0011614-1/2 | chr1:134394510-134394584 | Alt5 | 0,8433908 | 0,7162279 | 0,127162942 | 0,8506667 |
| Zbtb38 | MmuEX0052674 | chr9:96631711-96632161 | C2 | 0,8434165 | 0,7161021 | 0,127314333 | 0,8873333 |
| Stard3 | MmuINT0153680 | chr11:98240313-98241286 | IR-S | 0,2115694 | 0,0826069 | 0,128962506 | 0,8346667 |
| Gcfc1 | MmuINT0068106 | chr16:91027575-91030677 | IR-S | 0,3318565 | 0,2023473 | 0,129509154 | 0,8253333 |
| Zfp68 | MmuEX0053548 | chr5:139057698-139057787 | S | 0,7114357 | 0,5818803 | 0,129555397 | 0,86 |
| Mknk2 | MmuALTD0008560-1/2 | chr10:80138868- | Alt5 | 0,9028697 | 0,7721994 | 0,13067025 | 0,846 |
| Klc3 | MmuINT0087636 | chr7:19980517-19980587 | IR-C | 0,3840167 | 0,252842 | 0,131174768 | 0,8193333 |
| Ssh3 | MmuINT0152874 | chr19:4264109-4264313 | IR-S | 0,2672813 | 0,1354968 | 0,131784462 | 0,8193333 |
| Mef2a | MmuEX0028560 | chr7:74413155-74413286 | C3 | 0,6135948 | 0,4799287 | 0,133666101 | 0,8253333 |
| Rnpepl1 | MmuINT0136064 | chr1:94813838-94814210 | IR-S | 0,2184971 | 0,08467 | 0,133827152 | 0,9386667 |
| Klc3 | MmuINT0087637 | chr7:19980298-19980409 | IR-C | 0,4569387 | 0,3229695 | 0,133969165 | 0,874 |
| Msh3 | MmuEX0029790 | chr13:93123180-93123297 | S | 0,7662206 | 0,6314531 | 0,134767525 | 0,8633333 |
| Pcm1 | MmuEX0033988 | chr8:42360349-42360522 | S | 0,9082845 | 0,7734773 | 0,134807191 | 0,87 |
| 5430435G22Rik | MmuEX0001980 | chr1:133602173-133602298 | C1 | 0,5940483 | 0,4592097 | 0,134838606 | 0,8206667 |
| Rab11fip5 | MmuEX0038272 | chr6:85290337-85290491 | C2 | 0,3989762 | 0,263807 | 0,135169208 | 0,898 |
| Nup88 | MmuINT0112037 | chr11:70758488-70758704 | IR-C | 0,2157204 | 0,0801019 | 0,135618504 | 0,8133333 |
| Fibp | MmuINT0064523 | chr19:5464216-5464333 | IR-S | 0,2709194 | 0,1350897 | 0,13582966 | 0,916 |
| Irf3 | MmuINT0083304 | chr7:52253279-52254074 | IR-C | 0,2095747 | 0,0722046 | 0,137370092 | 0,9746667 |
| Clasp1 | MmuEX0011507 | chr1:120438252-120438275 | S | 0,1891587 | 0,0515258 | 0,137632923 | 0,894 |
| Atxn7l3 | MmuALTA0002430-1/2 | chr11:102154590-102154635 | Alt3 | 0,8978321 | 0,7585421 | 0,139290048 | 0,8446667 |
| Igsf9 | MmuEX0023842 | chr1:174419644-174419796 | C2 | 0,951005 | 0,8106519 | 0,140353094 | 0,936 |
| Trim3 | MmuINT0165094 | chr7:112759984-112761120 | IR-S | 0,3676723 | 0,2268679 | 0,140804465 | 0,8853333 |
| Wrb | MmuEX0052257 | chr16:96374583-96374645 | S | 0,9652827 | 0,8234751 | 0,141807608 | 0,8426667 |
| Trim11 | MmuINT0164965 | chr11:58795726-58802226 | IR-S | 0,3649832 | 0,2230205 | 0,141962647 | 0,8713333 |
| Slc25a37 | MmuINT0145663 | chr14:69865695-69867450 | IR-S | 0,6885801 | 0,5436866 | 0,144893435 | 0,928 |
| Zfp598 | MmuINT0179513 | chr17:24817977-24818053 | IR-S | 0,2628259 | 0,1168237 | 0,146002154 | 0,862 |
| Fam133b | MmuALTA0006793-1/4 | chr5:3559138-3559159 | Alt3 | 0,1939526 | 0,0476906 | 0,146261986 | 0,8013333 |
| Kif2a | MmuALTD0007462-1/2 | chr13:107778090-107778155 | Alt5 | 0,5517139 | 0,4044529 | 0,147260936 | 0,802 |
| Syvn1 | MmuINT0156329 | chr19:6052480-6052547 | IR-S | 0,2517794 | 0,1044071 | 0,147372357 | 0,9626667 |
| Usp1 | MmuEX0050811 | chr4:98593146-98593365 | S | 0,7694125 | 0,6213254 | 0,148087087 | 0,8786667 |
| Pitpnm1 | MmuINT0120274 | chr19:4111699-4111865 | IR-S | 0,3148845 | 0,1660081 | 0,148876406 | 0,942 |
| Dbnl | MmuALTA0005093-1/4 | chr11:5697290-5697338 | Alt3 | 0,6311117 | 0,4800965 | 0,15101517 | 0,89 |
| Stoml1 | MmuEX0045502 | chr9:58102741-58102847 | C1 | 0,9162442 | 0,7649631 | 0,151281131 | 0,8786667 |
| Slc25a22 | MmuINT0145559 | chr7:148617813-148617919 | IR-S | 0,3365945 | 0,185057 | 0,151537426 | 0,8713333 |
| Terf2 | MmuALTA0018073-1/2 | chr8:109600779-109600896 | Alt3 | 0,3666741 | 0,2137821 | 0,152892001 | 0,8686667 |
| Rbm15 | MmuINT0132595 | chr3:107129223-107133132 | IR-S | 0,4087696 | 0,25561 | 0,153159601 | 0,8893333 |
| Ncor2 | MmuALTA0011823-1/2 | chr5:125517644-125517746 | Alt3 | 0,2406819 | 0,0859379 | 0,154743994 | 0,854 |
| Sipa1 | MmuINT0143835 | chr19:5661112-5663603 | IR-C | 0,2440785 | 0,0869072 | 0,157171367 | 0,9386667 |
| 1810063B07Rik | MmuALTD0000212-1/2 | chr14:20902100- | Alt5 | 0,8932792 | 0,735326 | 0,157953265 | 0,96 |
| Dhrsx | MmuEX0014516 | NT_166291:31370-31471 | C1 | 0,8170732 | 0,658967 | 0,158106197 | 0,8566667 |
| Nup88 | MmuALTA0012434-1/2 | chr11:70758347-70758472 | Alt3 | 0,7839401 | 0,6244217 | 0,159518461 | 0,94 |
| Zfp7 | MmuINT0179739 | chr15:76718778-76720436 | IR-S | 0,3536458 | 0,1929014 | 0,16074432 | 0,9186667 |

| | | | | | | | |
|---|---|---|---|---|---|---|---|
| Diap1 | MmuEX0014636 | chr18:38066042-38066068 | S | 0,8375693 | 0,6747997 | 0,162769615 | 0,85 |
| Nat10 | MmuEX0030922 | chr2:103566873-103567045 | S | 0,8339928 | 0,6682544 | 0,165738376 | 0,8486667 |
| Ttll4 | MmuEX0049665 | chr1:74719418-74719584 | S | 0,9673515 | 0,799379 | 0,16797248 | 0,9946667 |
| Mink1 | MmuEX0029044 | chr11:70422910-70422933 | S | 0,362034 | 0,1936096 | 0,168424447 | 0,8886667 |
| Gpr124 | MmuINT0072933 | chr8:28220245-28220412 | IR-S | 0,455733 | 0,2858032 | 0,169929796 | 0,8353333 |
| Atp6v1e1 | MmuEX0006911 | chr6:120768352-120768417 | C1 | 0,870825 | 0,6998187 | 0,171006315 | 0,9713333 |
| Ogg1 | MmuINT0112996 | chr6:113278607-113279222 | IR-S | 0,4419171 | 0,2656175 | 0,176299521 | 0,9006667 |
| Mfap3 | MmuALTD0008457-1/3 | chr11:-57332214 | Alt5 | 0,7919435 | 0,6150726 | 0,176870868 | 0,8186667 |
| Pcnxl3 | MmuALTA0012873-1/3 | chr19:5681432-5681553 | Alt3 | 0,6526103 | 0,4741171 | 0,178493252 | 0,8386667 |
| Plekhg3 | MmuEX0035767 | chr12:77672591-77672629 | S | 0,9442094 | 0,7645407 | 0,179668755 | 0,8953333 |
| Sec61g | MmuALTD0012549-1/3 | chr11:16408103- | Alt5 | 0,4500681 | 0,2665522 | 0,183515941 | 0,908 |
| Tank | MmuALTA0017805-1/2 | chr2:61464963-61465071 | Alt3 | 0,7587042 | 0,5747137 | 0,183990568 | 0,8933333 |
| Amn1 | MmuEX0004800 | chr6:149133514-149133646 | C1 | 0,9366184 | 0,7522014 | 0,184417056 | 0,8726667 |
| Zfp428 | MmuALTA0020637-1/2 | chr7:25300095-25300121 | Alt3 | 0,6151911 | 0,4261129 | 0,189078225 | 0,8373333 |
| Chpf2 | MmuINT0036144 | chr5:24095480-24096126 | IR-S | 0,4342334 | 0,2423701 | 0,191863266 | 0,946 |
| 6330503K22Rik | MmuALTA0000628-1/2 | chr7:125867972-125868099 | Alt3 | 0,2586236 | 0,0654822 | 0,193141493 | 0,8693333 |
| AU019823 | MmuEX0002794 | chr9:50424763-50424789 | S | 0,3287905 | 0,1264678 | 0,202322706 | 0,9473333 |
| Setd6 | MmuINT0142186 | chr8:98240942-98241820 | IR-S | 0,3510388 | 0,1458754 | 0,205163405 | 0,936 |
| Nat10 | MmuEX0030920 | chr2:103568157-103568253 | S | 0,7848231 | 0,5707078 | 0,214115364 | 0,8066667 |
| Igdcc4 | MmuALTA0009019-1/2 | chr9:64974565-64974735 | Alt3 | 0,7301875 | 0,4493135 | 0,280873977 | 0,8453333 |
| Taf4a | MmuALTD0013961-1/3 | chr2:179666721-179666930 | Alt5 | 0,7643013 | 0,4588023 | 0,305499017 | 0,95 |

| | Considered | Pdiff>0.8 | Volcano plot (all non redundant) | Violin plot (IR,EX Pdiff>0.8) | Differentially spliced (Pdiff>0.8, \|ΔPSI\|>0.5) |
|---|---|---|---|---|---|
| IR | 63180 | 849 | 63180 | 849 | 262 |
| EX | 22160 | 741 | 22160 | 741 | 358 |
| Alt 5' | 17103 | 434 | 7752 | | 65 |
| Alt 3' | 23892 | 240 | 10957 | | 89 |
| TOTAL | 126335 | 2264 | 104049 | 1590 | 774 |
