## Supplemental figure 1 for "The Splicing Factor XAB2 interacts with ERCC1-XPF and XPG for RNA-loop processing during mammalian development"

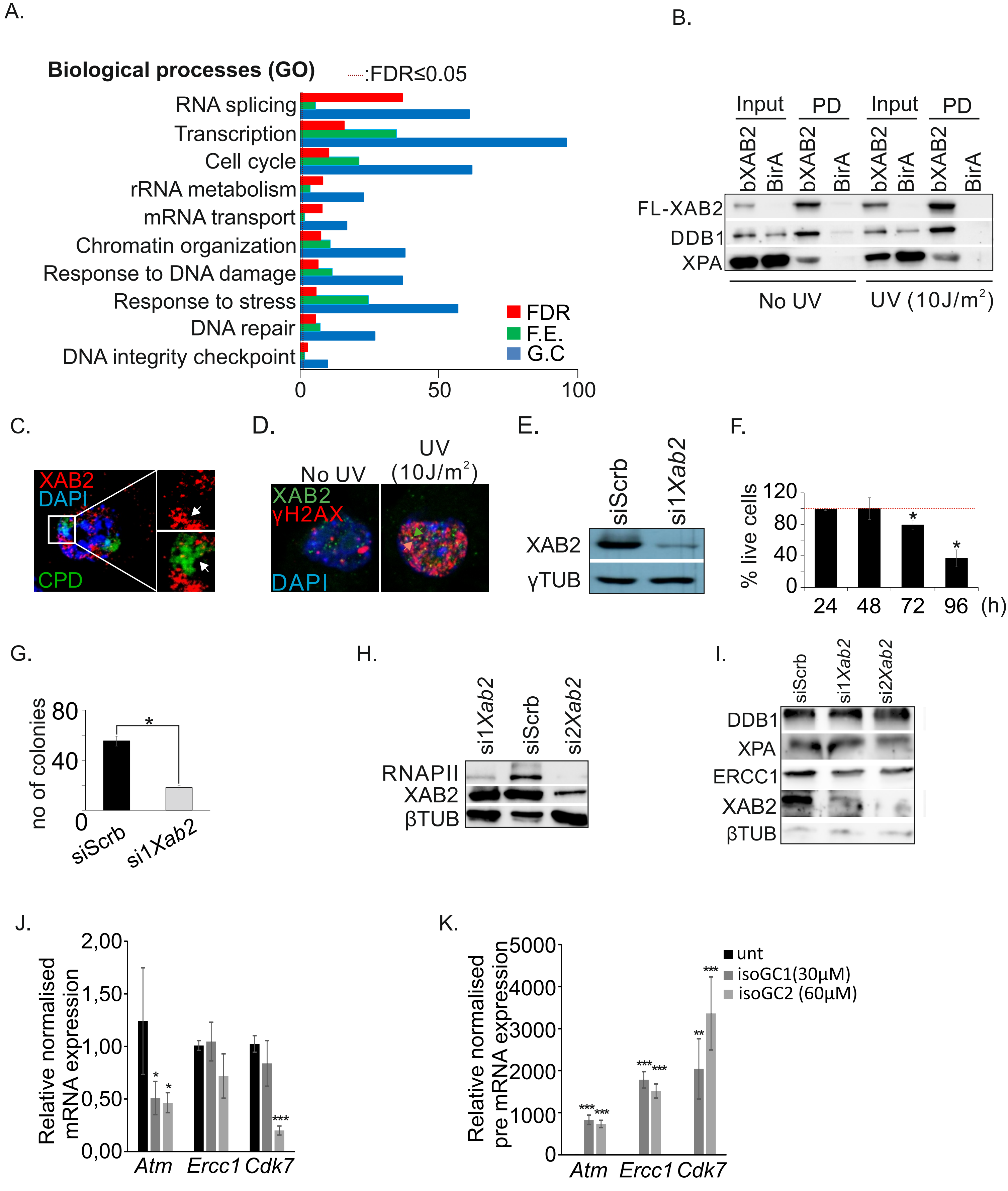


L.

| | Considered | Pdiff>0.8 | Volvano (all non redudant) | Violin plots (IR, EX, Pdiff >0.8) | Differentially spliced Pdiff>0.8 and \|ΔPSI\|>5 |
|---|---|---|---|---|---|
| IR | 63180 | 849 | 63180 | 849 | 262 |
| EX | 22160 | 741 | 22160 | 741 | 358 |
| Alt3' | 23892 | 434 | 10957 | | 89 |
| Alt5' | 17103 | 240 | 7752 | | 65 |
| Total | 126335 | 2264 | 104049 | 1590 | 774 |
