## Supplemental figure 2 for "The Splicing Factor XAB2 interacts with ERCC1-XPF and XPG for RNA-loop processing during mammalian development"

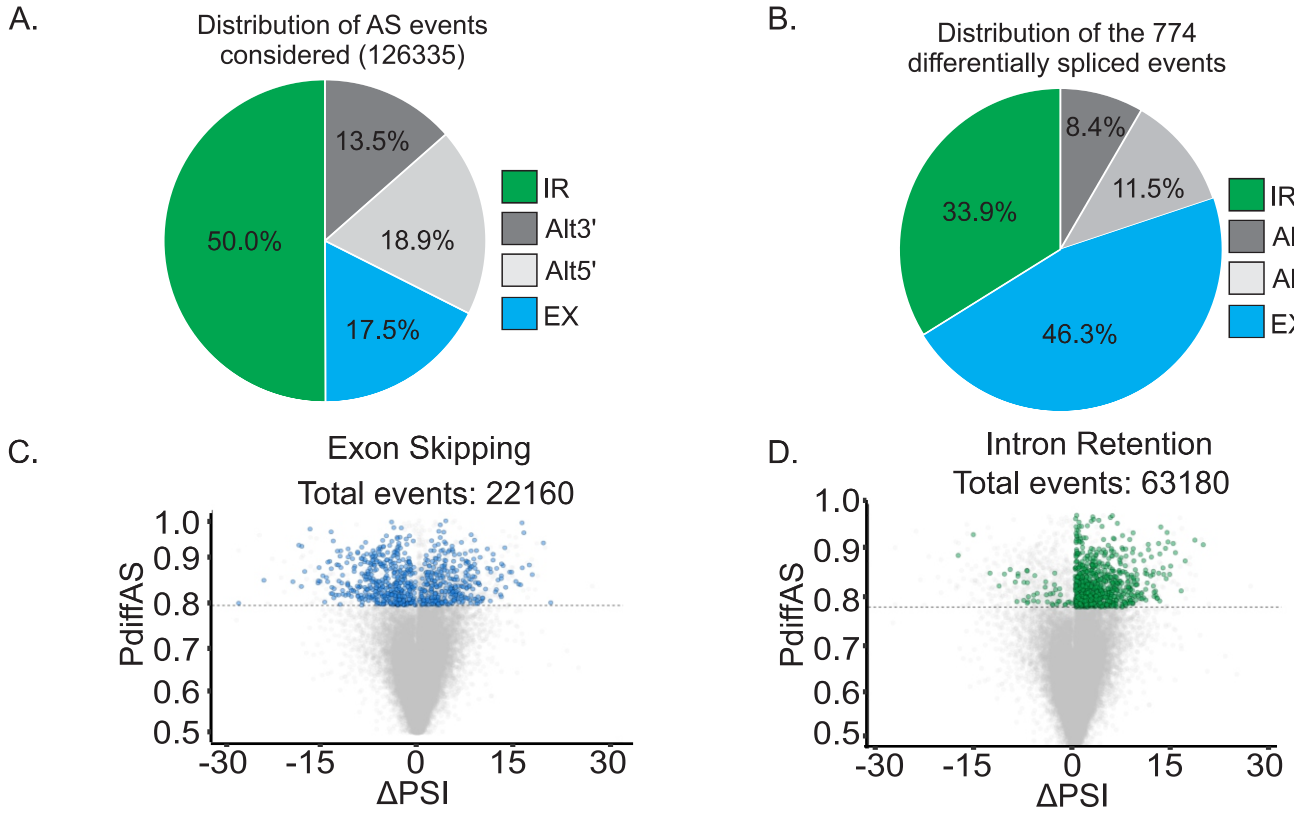


E.

| | Considered | Pdiff>0.8 | Volvano (all non redudant) | Violin plots (IR, EX, Pdiff >0.8) | Differentially spliced Pdiff>0.8 and \|ΔPSI\|>5 |
|---|---|---|---|---|---|
| **IR** | 28917 | 1558 | 28917 | 1558 | 748 |
| **EX** | 7468 | 347 | 7468 | 347 | 207 |
| **Alt3'** | 12156 | 412 | 5545 | | 123 |
| **Alt5'** | 8136 | 182 | 3676 | | 56 |
| **Total** | 56677 | 2499 | 45606 | 1905 | 1134 |

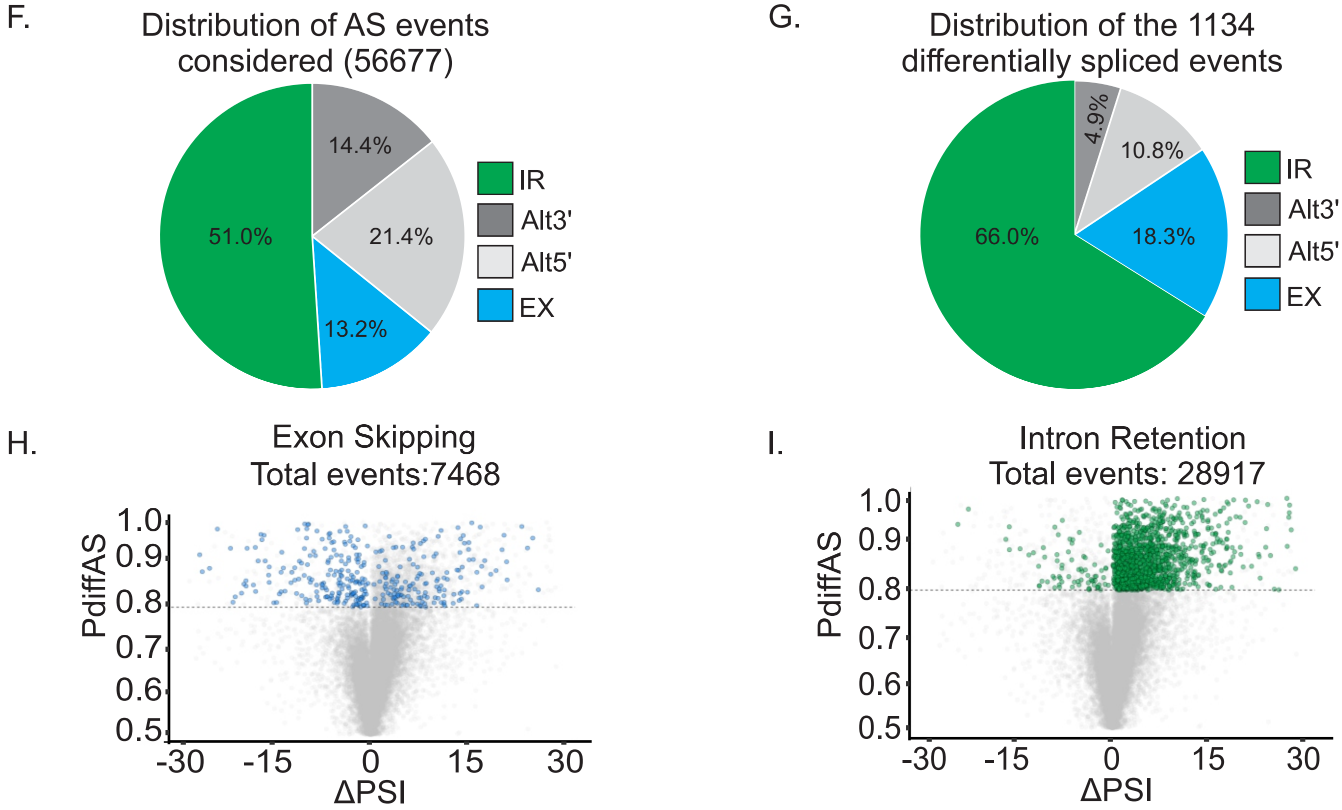


Goulielmaki et al., Fig. S2
