## Supplementary figures and images for "The Splicing Factor XAB2 interacts with ERCC1-XPF and XPG for RNA-loop processing during mammalian development"

### Supplemental figure 3

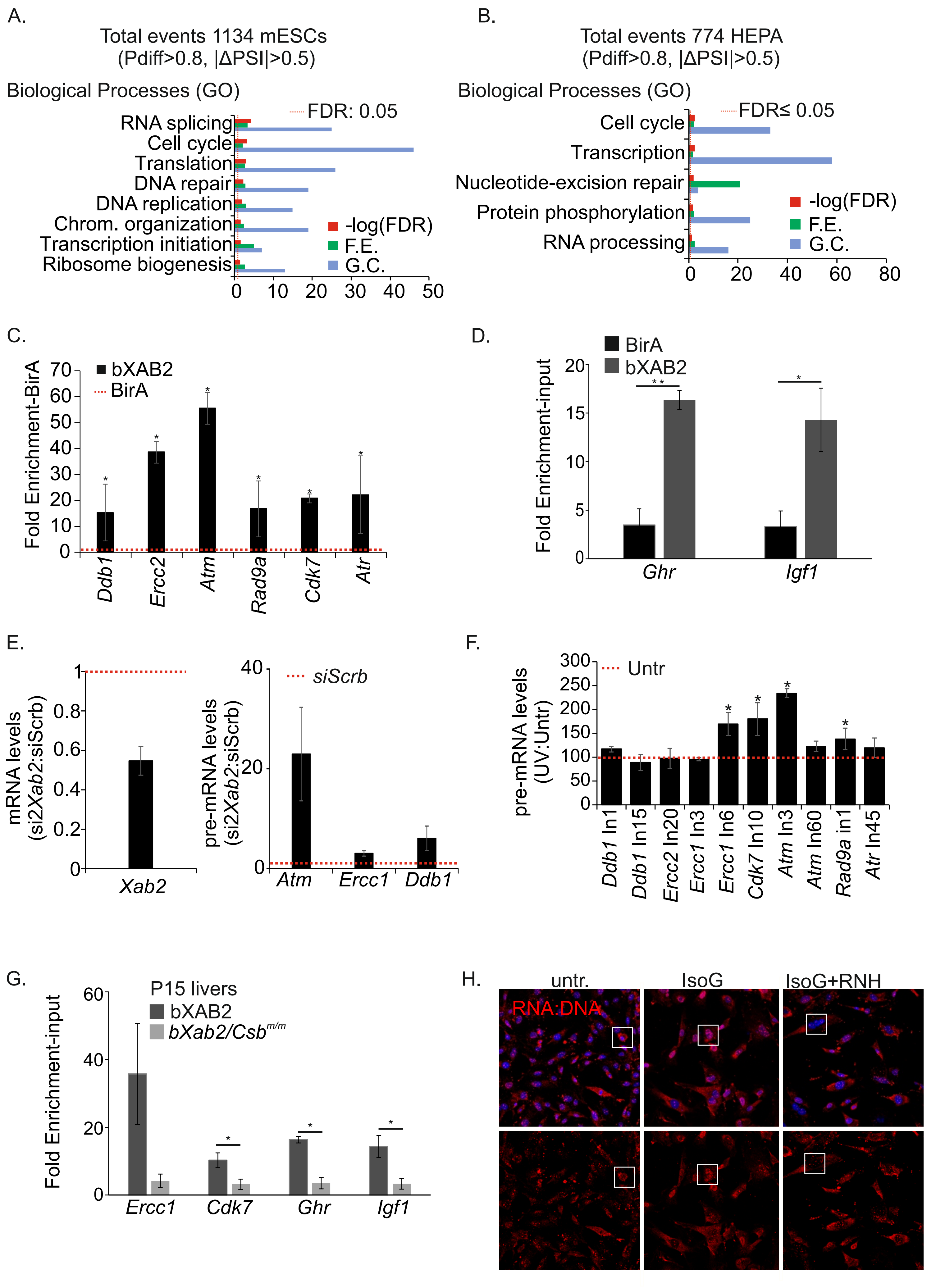


Goulielmaki et al., Fig. S3

### Supplemental figure 4

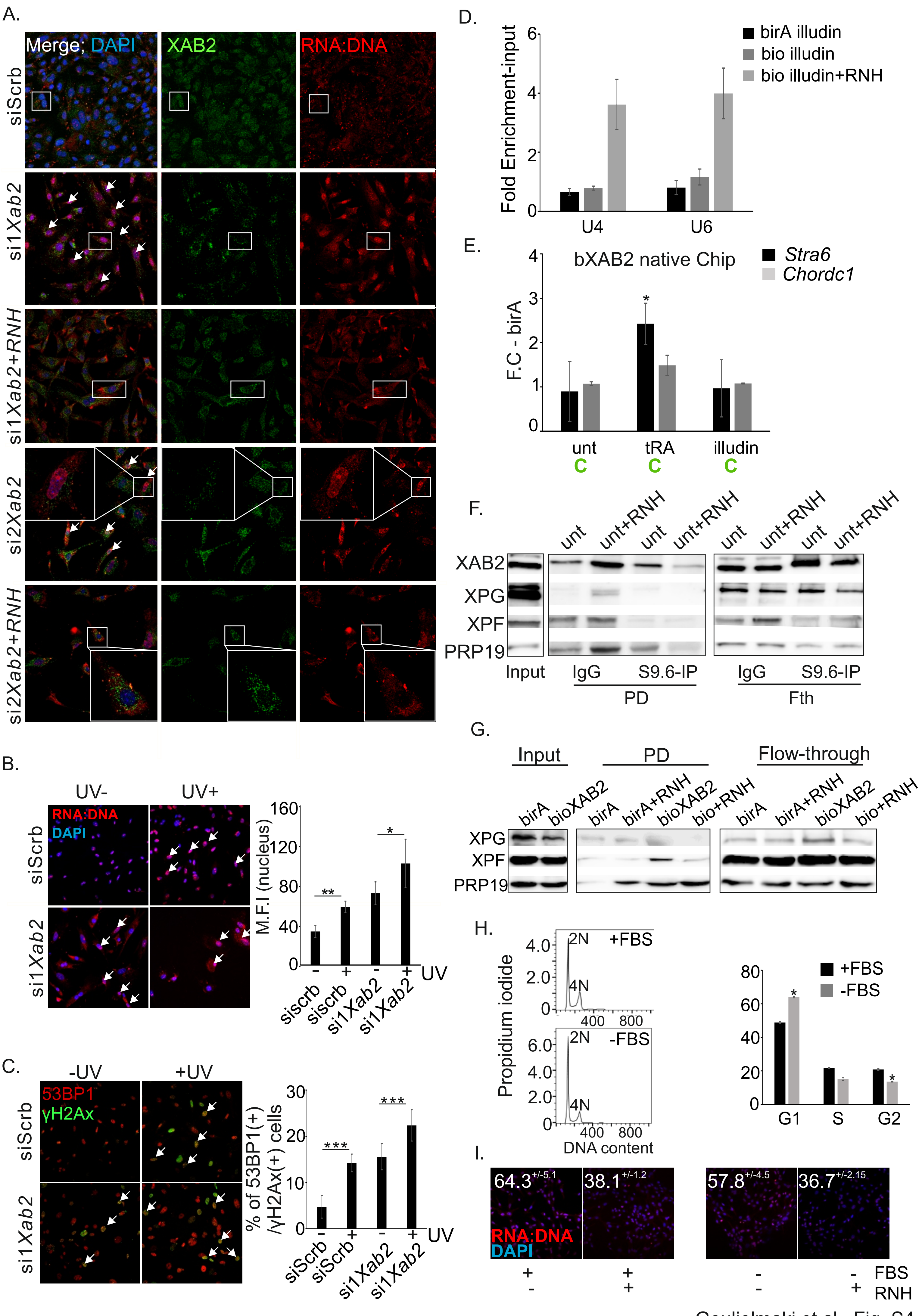


Goulielmaki et al., Fig. S4

### Supplemental figure 5

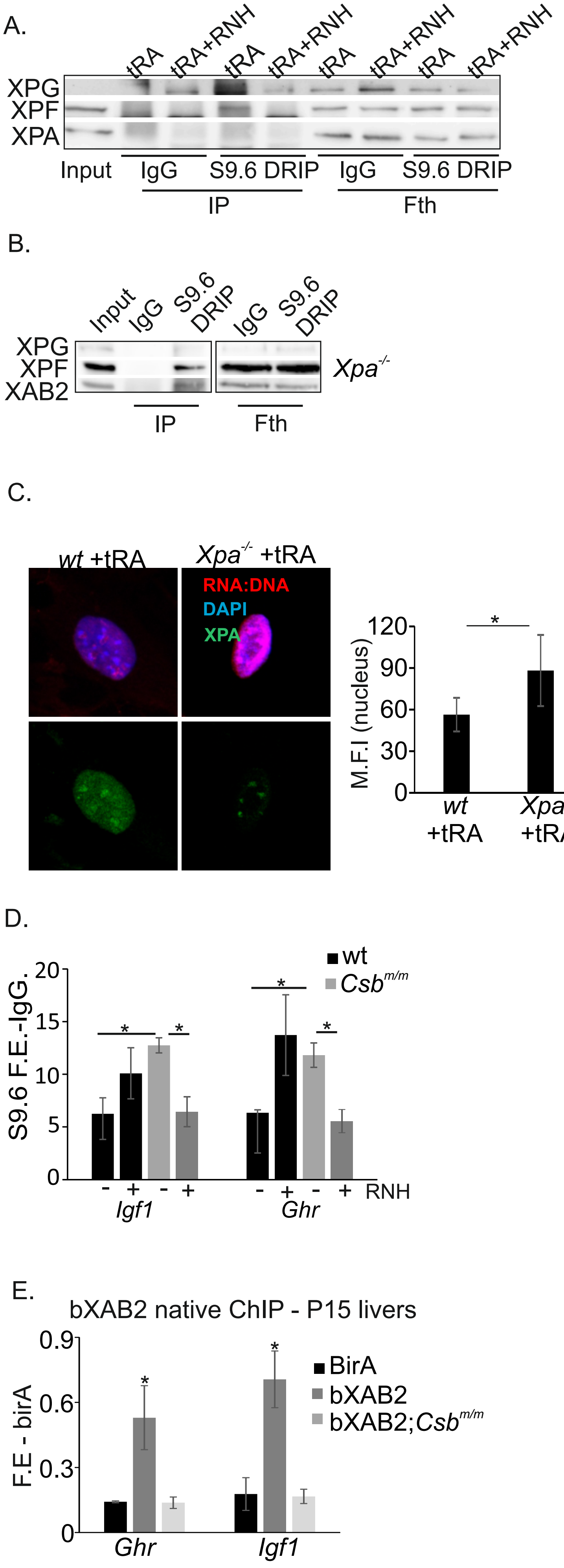
